# Unique Effects of Sedatives, Dissociatives, Psychedelics, Stimulants, and Cannabinoids on Episodic Memory: A Review and Reanalysis of Acute Drug Effects on Recollection, Familiarity, and Metamemory

**DOI:** 10.1101/2022.05.20.492842

**Authors:** Manoj K. Doss, Jason Samaha, Frederick S. Barrett, Roland R. Griffiths, Harriet de Wit, David A. Gallo, Joshua D. Koen

## Abstract

Despite distinct classes of psychoactive drugs producing putatively unique states of consciousness, there is surprising overlap in terms of their effects on episodic memory and cognition more generally. Episodic memory is supported by multiple subprocesses that have been mostly overlooked in psychopharmacology and could differentiate drug classes. Here, we reanalyzed episodic memory confidence data from 10 previously published datasets (28 drug conditions total) using signal detection models to estimate 2 conscious states involved in episodic memory and 1 consciously-controlled metacognitive process of memory: the retrieval of specific details from one’s past (recollection), noetic recognition in the absence of retrieved details (familiarity), and accurate introspection of memory decisions (metamemory). We observed that sedatives, dissociatives, psychedelics, stimulants, and cannabinoids had unique patterns of effects on these mnemonic processes dependent on which phase of memory (encoding, consolidation, or retrieval) was targeted. All drugs at encoding except stimulants impaired recollection, and sedatives, dissociatives, and cannabinoids at encoding impaired familiarity. The effects of sedatives on metamemory were mixed, whereas dissociatives and cannabinoids at encoding tended to enhance metamemory. Surprisingly, psychedelics at encoding tended to enhance familiarity and did not impact metamemory. Stimulants at encoding and retrieval enhanced metamemory, but at consolidation, they impaired metamemory. Together, these findings may have relevance to mechanisms underlying unique subjective phenomena under different drug classes, such as blackouts from sedatives or *déjà vu* from psychedelics. This study provides a framework for interrogating drug effects within a domain of cognition beyond the global impairments on task performance typically reported in psychopharmacology.

**Public significance statement:** This systematic review and reanalysis of several datasets indicate that sedatives (alcohol, zolpidem, triazolam), dissociatives (ketamine, dextromethorphan), psychedelics (psilocybin, MDMA), stimulants (dextroamphetamine, dextromethamphetamine), and cannabinoids (THC) can each have idiosyncratic effects on episodic memory, differentially impairing certain mnemonic processes while sparing or even facilitating others. Such findings inform how different drugs can produce unique subjective phenomena and provide a framework for future work to differentiate the effects of psychoactive drugs within a domain of cognition.

## Introduction

Despite anecdotal reports attributing unique “altered states of consciousness” to different psychoactive drugs, pharmacologically distinct drugs can have largely similar effects on cognitive processes relevant to different components of consciousness, such as episodic memory and metacognition (LeDoux and Lau, 2020). Episodic memory is the conscious reexperiencing of information from the past, and metacognition is the awareness of one’s own thought processes. Several classes of psychoactive drugs are found to impair the formation (i.e., encoding) of episodic memories (Carter, Kleykamp, *et al*., 2013; e.g., Barrett *et al*., 2018; Doss, Weafer, Gallo, *et al*., 2018a; Doss, Weafer, *et al*., 2020), have little impact on the post-encoding stabilization (i.e., consolidation) of memories (Parker *et al*., 1980; Ballard *et al*., 2015), and increase false memories when remembering (i.e., retrieving) memories (Ballard *et al*., 2014; Doss, Weafer, Gallo, *et al*., 2018a; b). Most drugs also seemingly impair metacognition (Carter *et al*., 2009; Mintzer *et al*., 2010; Carter, Kleykamp, *et al*., 2013; Adam *et al*., 2020). Finally, sedatives (Weber *et al*., 2014), dissociatives (Bonhomme *et al*., 2016), psychedelics (Mason *et al*., 2020), stimulants (Schrantee *et al*., 2016), cannabinoids (Wall *et al*., 2019), and drugs of other classes (Bajo *et al*., 2015; Doss, May, *et al*., 2020) attenuate task-free functional connectivity within the default mode network, a network that supports episodic memory and consciousness (Vanhaudenhuyse *et al*., 2010; Ranganath and Ritchey, 2012).

Although psychopharmacology has extensively investigated non-episodic memory (e.g., basal ganglia-mediated reward and procedural learning and amygdala-mediated conditioning; Koob and Volkow, 2016; Scofield *et al*., 2016), the utility of episodic memory to the understanding of how different drugs might uniquely produce addictive behaviors has been underexplored (Müller, 2013; but see Bornstein and Pickard, 2020; Kloft *et al*., 2021). Discovering selective drug effects on different mnemonic processes and other behaviors will be important to the development of novel therapeutics. With the recent push for medicalizing drugs that were previously thought to have no utility in psychiatry (e.g., dissociatives, psychedelics, and cannabinoids), understanding how these drugs impact episodic memory may be crucial for maximizing their benefits (e.g., altering maladaptive mnemonic representations) while minimizing their harms (e.g., avoiding false memory induction).

Although pharmacologically distinct drugs may be differentiated across domains of cognition (e.g., dissociatives impair episodic memory more than psychedelics but vice versa for working memory; Barrett *et al*., 2018), within episodic memory there exists the potential for unique patterns of effects on mnemonic subprocesses that have been overlooked due to the use of standardized tasks and measures. One approach to differentiating drug effects on memory is to use specific task manipulations on which drugs may have selective effects. For example, the sedative midazolam impairs the encoding of contextual information as indicated by a lack of a boost in memory performance when a context is reinstated during memory retrieval (Reder *et al*., 2013). In contrast, the cannabinoid Δ^9^-tetrahydrocannabinol (THC) has been found to spare context reinstatement effects on memory (Doss, Weafer, *et al*., 2020).

Another approach to differentiating drug effects on episodic memory is using computational modeling to estimate the contribution of different mnemonic subprocesses to performance on a memory task. Not only might this approach highlight differences between drugs, it can be more sensitive to drug effects than standard measures of cognition that do not differentiate key mnemonic subprocesses (see Doss, Weafer, Gallo, *et al*., 2018a). Despite the existence of several drug studies in which confidence ratings for memory decisions were collected (i.e., confidence on a Likert scale during a recognition memory test), few of these have interrogated these data with established computational models. Here, we utilized two computational models of memory confidence to explore unique patterns of drug effects on recollection, familiarity, and metamemory. Whereas recollection involves the retrieval of specific details and associations of a past experience, familiarity is a general feeling of knowing that a stimulus has been experienced without recalling any specific details about the prior encounter. In contrast, metamemory, while not a direct form of memory *per se*, refers to one’s ability to introspect on their own episodic memory (see below for in-depth descriptions). Not only can all of these mnemonic processes be estimated from a single memory test, they may support distinct aspects of consciousness and be selective targets of different psychoactive drugs. We begin by briefly introducing recollection, familiarity, and metamemory and how they may be useful in understanding psychiatric illness and drug effects. Then we introduce how these mnemonic processes can be estimated from confidence ratings on a recognition memory test using the dual process signal detection (DPSD) model and the meta-*d*’ model and review drug effects on episodic memory of the five classes of drug for which we were able to acquire data (sedatives, dissociatives, psychedelics, stimulants, and cannabinoids). This review is followed by a reanalysis of 10 datasets with 28 different drug manipulations using the reviewed signal detections models. Finally, we end with a discussion of novel drug effects, what might be predicted from the unique effects of different drug classes on specific memory processes, and guidance for future work on the effects of psychoactive drugs on episodic memory. Although we largely focus on human data, we also describe animal literature including relevant neurobiology, behavioral proxies to human paradigms, and drug effects for which human data is nonexistent.

## Recollection, Familiarity, and Metamemory

Recent neurocognitive models have explored interactions between the mnemonic representations and metacognitive operations that contribute to conscious experience (Dehaene and Changeux, 2011; Bastin *et al*., 2019; LeDoux and Lau, 2020). For example, recollection and familiarity have been proposed to support autonoetic consciousness (the ability to place oneself in the past or future) and noetic consciousness (awareness of objects and events), respectively (Tulving, 1985, 2002; Gardiner, 2001; Conway, 2005), and metacognition has been argued to be central to conscious awareness (Koriat, 2007; Lau and Rosenthal, 2011; Brown *et al*., 2019). A recent model postulates that distinct conscious experiences require qualitatively different memories and corresponding meta-representations (LeDoux and Lau, 2020). Therefore, measuring the effects of drugs on recollection, familiarity, and metamemory may allow us to capture idiosyncratic “altered states of consciousness.” Additionally, of relevance to models of episodic memory, dissociable effects of different drugs on measures corresponding to mnemonic representations and metamemory would provide evidence that memory and metamemory processes are separable.

### Recollection and Familiarity

Dual process models of memory propose that two qualitatively distinct processes contribute to memory performance, namely recollection and familiarity (Yonelinas, 2002; Yonelinas *et al*., 2010). Recollection reflects the retrieval of specific details and associations from past experiences including “where” and “when” information (e.g., recalling that you saw your barista yesterday on the bus). Typically, recollection is associated with a subjective reliving of an experience and is dependent on the function of the hippocampus. In contrast, familiarity is a feeling of knowing that a stimulus has been experienced in the absence of any corroborating evidence (e.g., recognizing your barista’s face while on the bus but not remembering from where you know them). Such familiarity-based memories are thought to be supported by the ease or fluency with which information is processed through sensory hierarchies and semantic memory stores (Johnston *et al*., 1985; Wang and Yonelinas, 2012; Ozubko and Yonelinas, 2014) with particular support from perirhinal cortex (Yonelinas *et al*., 2010). Of relevance, recollection and familiarity are thought to lead to varying levels of confidence (i.e., on a Likert scale) in memory decisions on a recognition test. Recollection generally leads to higher confidence in one’s memory and more accurate memory decisions than familiarity, whereas familiarity leads to a lower level of confidence on average and can make one more likely to erroneously attribute familiarity to a novel stimulus. In animal models, behavioral proxies of recollection include hippocampally-dependent tasks involving temporal order, spatial locations, or memory guided by contextual cues such as maze tasks (Eichenbaum *et al*., 2005, 2007). For familiarity, novel object recognition has served as a behavioral proxy in animal models with the perirhinal cortex supporting this function (Eichenbaum *et al*., 2007; Cohen and Stackman Jr., 2015).

There is a large literature demonstrating that recollection and familiarity are dissociable across a variety of non-pharmacological manipulations and conditions. For example, changing the perceptual modality on a recognition memory test (e.g., first presenting a word visually and then testing memory for that word auditorily) impairs familiarity more so than recollection, whereas requiring a response deadline (e.g., <1 second) on a memory test impairs recollection more so than familiarity (reviewed in Yonelinas, 2002). Consistent with this latter effect, event-related potentials associated with familiarity emerge before those of recollection (Rugg and Curran, 2007). The hippocampus rapidly encodes memories (i.e., one-shot learning) in contrast to the slow learning cortex (Norman and O’Reilly, 2003), and recollection is thought to be more flexible than familiarity such that recollection can be subsequently used to update memories with new associations (Ozubko *et al*., 2017). Familiarity, on the other hand, can lead to intrusions in memory that may no longer be relevant, especially when unconstrained by recollection. Such data might speak to the cognitive rigidity in psychiatric disorders such as depression and schizophrenia that exhibit disproportionate impairments in recollection (Hertel and Milan, 1994; Libby *et al*., 2013; Stange *et al*., 2017; Waltz, 2017), which may preclude one from updating their mental model. Under laboratory conditions in which familiarity is high but recollection fails, healthy participants report *déjà vu* (feeling that one has experienced an event despite not having actually experienced it; Cleary *et al*., 2012; Cleary and Claxton, 2018). *Déjà vu* is also experienced for prolonged periods in patients with schizophrenia (Adachi *et al*., 2006) and can be evoked by psychoactive drugs (especially hallucinogens; Singh, 2007; Siomopoulos, 1972; Sno, 2000; Wells *et al*., 2014; for discussion on its relatively rare occurence, see Honey *et al*., 2006, 2003; Peden *et al*., 1981)

Although dual process models typically focus on measuring the frequency of accurate recollection, these processes can interact, and recollection can become distorted resulting in false memories (Gallo and Roediger, 2003; see Doss *et al*., 2016). False memory induction may be particularly prominent when recollection is reduced and less able to constrain feelings of familiarity or partial recollections of an event (Brainerd *et al*., 2014; Doss *et al*., 2016, 2019). For example, a picture in a never-before-visited house may feel familiar, but if one can recollect from where they originally saw the picture, they may be less likely to misattribute the familiarity of the picture to having visited the house. In addition to more selectively impairing recollection (as discussed below), some drugs such as psychedelics and THC might enhance other cognitive processes that that can distort memory such as mental imagery (Ames, 1958; Tart, 1970; Carhart-Harris *et al*., 2012, 2014). Thus, multiple mechanisms of false memory induction may be driven by some drugs under certain conditions (reviewed in Kloft *et al*., 2021).

### Metamemory

Metamemory is a domain of metacognition that refers to the understanding of one’s own memory abilities, including prospective metamemory, how well one knows what they will remember, and retrospective metamemory, how well one knows a retrieved memory (e.g., monitoring a memory for its veracity). We focus on retrospective metamemory here, as drug studies have typically implemented retrospective memory judgements, and unless otherwise stated, we use “metamemory” in the retrospective sense. Metamemory, and metacognition in general, can be measured as the degree to which confidence tracks performance (e.g., confident that you did well on a test when in fact you did do well on a test). Although one may be highly confident when recognizing a previously seen stimulus based on the recollection of thoughts one had when first seeing that stimulus or the experienced familiarity, confidence in one’s memory can also be influenced by how well they understands their own memory system and their resulting expectations (e.g., Wong *et al*., 2012). One who is confident in their memory when they accurately remember a stimulus or event and not very confident when their memory is less accurate would be said to have good metamemory. In contrast, someone who is less confident when their memory is accurate but overly confident when their memory is inaccurate would have a poor metamemory. Such misunderstandings of one’s memory can result in memory distortions (Gallo and Lampinen, 2015). There appears to be a domain general metacognitive brain network including medial and lateral prefrontal cortex (Fleming and Dolan, 2012; Vaccaro and Fleming, 2018), and disengagement of lateral prefrontal cortex can produce memory distortions (cf. Gallo *et al*., 2006, 2010; McDonough *et al*., 2013; Zhu *et al*., 2019).

Recently, there has been increased interest in metacognitive processes in psychiatry, as metacognition can be enhanced through training (Carpenter *et al*., 2019), and treatments aimed at improving metacognition may have some success in improving psychiatric illnesses (Cella *et al*., 2015; Hamonniere and Varescon, 2018; Moritz *et al*., 2019; Philipp *et al*., 2019). Anosognosia, or the lack of insight into one’s own condition or cognitive deficits, is an extreme case of metacognitive failure that could be a factor in the discontinuation of medication in schizophrenia (David *et al*., 2012; Lehrer and Lorenz, 2014). Patients with schizophrenia are impaired in both prospective (Chiu *et al*., 2015) and retrospective (Moritz *et al*., 2008; Peters *et al*., 2013; Eifler *et al*., 2015) metamemory, and higher doses of antipsychotics, drugs that typically attenuate dopaminergic function, have been associated with less impairment in retrospective metamemory specifically (Moritz *et al*., 2008). In contrast, enhancing dopaminergic signaling in healthy participants can impair metamemory (Clos *et al*., 2019). Such effects of dopamine may not apply to all metacognitive processes, however, as metacognition of agency can be enhanced with methamphetamine (Kirkpatrick *et al*., 2008).

A lack of insight into one’s deficits may also play a role in different addictions (Goldstein *et al*., 2009; Spada *et al*., 2015; Le Berre and Sullivan, 2016; Hamonniere and Varescon, 2018). Patients with cocaine addiction (Moeller *et al*., 2016) and patients with opioid addiction who are receiving methadone maintenance treatment (Sadeghi *et al*., 2017) were found to have impaired perceptual metacognition. Although these patients with opioid addiction also had an impairment in memory, their retrospective metamemory was not impaired. Patients with alcohol use disorder were also not found to have impairments in retrospective metamemory, but they were impaired in prospective metamemory (Le Berre *et al*., 2010, 2016).

Interestingly, anxiety and depression are associated with better perceptual metacognition (NSPN Consortium *et al*., 2017; Rouault *et al*., 2018), but compulsion and intrusive thoughts are associated with worse perceptual metacognition. An impairment in metamemory could underlie obsessive checking even if memory is intact. Consistent with this idea, several pieces of evidence point to impairments in retrospective metamemory in obsessive compulsive disorder (Tolin *et al*., 2001; Tuna *et al*., 2005; Exner *et al*., 2009; Radomsky *et al*., 2014; Moritz and Jaeger, 2018)(but see Moritz, Kloss, *et al*., 2009; Moritz, Ruhe, *et al*., 2009) with persistent checking leading to impairments in metamemory but not memory accuracy (van den Hout and Kindt, 2003b; a). Although some anxious and depressive symptomology may give one more realistic expectations of their performance (similar to reductions in an optimism bias, Sharot, 2011; Rouault *et al*., 2018), subjective cognitive failures better predict depression than objective cognitive failures (David *et al*., 2002). Believing that one is performing poorly in the absence of an actual deficit may give rise to a distorted sense of self that could precede or even contribute to illness-related cognitive deficits (Eisenacher *et al*., 2015; Eisenacher and Zink, 2017). Although speculative, extended use of cognitively impairing drugs could produce the belief that one has a poor memory even when one is sober. Alternatively, certain drugs (e.g., stimulants) may enhance memory and cognition to better match with one’s inflated expectations (cf. van den Hout and Kindt, 2003b), thereby driving subsequent use.

## Phases of Episodic Memory: Encoding, Consolidation, and Retrieval

Standard episodic memory paradigms contain an encoding phase followed by a retrieval phase with most work in psychopharmacology administering a drug prior to the encoding phase. During the encoding phase of a typical memory paradigm, participants are presented with a randomized list of stimuli such as individual words or pictures (Figure Ia). For each stimulus, a judgement is typically made to ensure that participants are paying attention and also to ensure that the “depth” of processing is held constant across experimental conditions (Craik and Lockhart, 1972). Depth of processing refers to whether an individual is attending to “shallow” features of a stimulus (e.g., “Are the letters of the word in lower or upper case?”) or “deep” features of a stimulus (e.g., “Is the word pleasant or not?”) during encoding. Because semantic (i.e., deep) judgements typically improve memory over perceptual (i.e., shallow) judgements, holding encoding judgements constant minimizes differences in cognitive operations that could spontaneously vary across experimental conditions. This may be especially important for pharmacological studies in which drug effects take place during memory encoding, as certain drugs may make one more or less likely to engage in semantic encoding strategies that are known to benefit memory.

After the encoding phase but before testing memory, a delay period is sometimes implemented to allow memories to stabilize or “consolidate.” Whereas the stabilization of newly formed synapses, or cellular consolidation, is thought to take place within hours, the transfer of episodic memories from the hippocampus to the cortex, or systems consolidation, can take up to years (Dudai, 2004; Mednick *et al*., 2011; Kandel *et al*., 2014). The existence of systems consolidation has recently been debated (Yonelinas *et al*., 2019), but importantly, the ability to retroactively enhance or impair recently encoded memories appears to be time-dependent. For example, interference effects from new learning degrade within minutes after initial learning (reviewed in Wixted, 2004). Here, we refer to pharmacological manipulations of consolidation as those that take place immediately post-encoding when drugs may be able to retroactively modulate memory for stimuli that were encoded prior to drug administration (cf. post-encoding enhancements with stress; Shields *et al*., 2017). As will be discussed, such post-encoding effects can drastically differ from pre-encoding effects.

The testing of previously encoded memories is referred to as the retrieval phase. Perhaps the most common test of memory for verbal stimuli in psychopharmacology is free recall. In this test, participants simply recall (via writing, typing, or saying aloud) as many words as they can from the encoding phase in whatever order. Performance on this task predominantly relies on recollection, as participants must self-generate each word instead of evaluating a stimulus for subjective novelty (for which familiarity can be used). Hippocampally-dependent temporal context plays an important role in free recall with the retrieval of a given word increasing the likelihood to subsequently retrieve proximate words (Polyn *et al*., 2009).

## Signal Detection Models of Recognition Memory

Another common way to assess memory is to use recognition memory tests (Figure Ib). On a typical recognition test, stimuli from the encoding phase (known as “targets”) are randomly intermixed with stimuli that were not previously presented in the encoding phase (known as “lures” or “foils”). For each stimulus, participants are asked to determine whether it is “old” or “new” (previously observed or not observed, respectively). Importantly, recognition tests can lead to different conclusions about memory relative to recall tests. For instance, whereas hippocampal amnesic patients typically perform at floor on tests of verbal free recall, they can perform above chance on recognition memory tests because of spared familiarity (Yonelinas, 2002; Yonelinas *et al*., 2010). Recognition memory tests, which are less frequently used in psychopharmacology research, have the potential to shed light on multiple mnemonic processes to which recall tests may be less sensitive.

There are four outcomes, or classifications, of trials that can occur on a recognition test depending on the status of the stimuli (target or lure) and the participant’s response (“old” or “new”). Correctly identifying a stimulus from encoding as previously observed is known as a “hit,” whereas misidentifying a stimulus that was not previously presented as “old” is known as a “false alarm.” These two types of trials contribute to the main measures of recognition memory accuracy, namely the hit rate (i.e., the proportion of target stimuli attracting an “old” response or *p*[“old”|target]) and the false alarm rate (i.e., the proportion of lure stimuli attracting an “old” response or *p*[“old”|lure]). Note that incorrectly identifying a target as “new” is considered a “miss” and correctly identifying a lure as “new” is known as a “correct rejection.” However, these measures are typically of no interest, as miss and correction rejection rates are equivalent to 1 minus the hit and false alarm rates, respectively.

It is now well-established that at least two processes, namely recollection and familiarity, support accurate responding on recognition memory tests (Yonelinas, 2002; Yonelinas *et al*., 2010). However, simple recognition tests with binary “old” and “new” responses as those described in the preceding paragraph are unable to tease apart these processes (Yonelinas and Parks, 2007). To overcome this limitation, a variety of procedures have been developed to quantify the contribution of recollection and familiarity more readily. One such method is known as the process dissociation procedure (Jacoby, 1991; Yonelinas and Jacoby, 2012). Unlike standard recognition memory tests, this procedure typically implements an encoding phase with two types of stimuli, such as words presented in temporally separated lists, and two different memory tests that put recollection and familiarity in opposition to each other. One test, the exclusion test, requires accepting stimuli as “old” only if they appeared on one of the two lists, thereby requiring recollection of “when” information to exclude stimuli that are otherwise familiar (i.e., those on the non-criterial list). The other test requires accepting any stimulus from either list as “old,” thereby making one use relatively more familiarity than the exclusion test. Based on the independence of recollection and familiarity, the probabilities of accepting stimuli from encoding on either memory test can be used to estimate the contributions of recollection and familiarity. Another method is the remember/know procedure (Tulving, 1985), which asks participants to report subjective experiences of recollection and familiarity by making a “remember” or “know” response, respectively. Tulving (1985) originally conceived “remember” responses as capturing autonoetic consciousness and “know” responses capturing noetic consciousness. More specifically, a “remember” response is given for a stimulus thought to be old when details or associations made at the time of encoding a stimulus are recollected, whereas a “know” response is given if a participant simply feels that a stimulus is familiar in the absence of any corroborating evidence. Given that recollection and familiarity are thought to be independent and operate in parallel (i.e., they can co-occur or each occur in the absence of the other), “know” responses are an underestimate of familiarity (i.e., they measure familiarity in the absence of recollection). Thus, an independence correction was developed to better estimate familiarity (Yonelinas and Jacoby, 1995).

One last method for dissociating recollection and familiarity that obtains measures of response bias (responses made for non-mnemonic reasons; see below) is to have participants report their confidence in their old/new recognition decisions in a second response or in a single response on a hybrid scale (see Figure Ib for a hybrid yes/no confidence response on a 6-point Likert scale). These confidence data can then be analyzed with signal detection theory to estimate different mnemonic processes (Koen *et al*., 2017; Yonelinas and Parks, 2007). Confidence in one’s memory for an event can be shaped not only by the strength of memory representations and the quantity of unique episodic details but also by knowledge of one’s own episodic memory system (i.e., metamemory). Different signal detection models have been developed to tease apart such influences on memory confidence. The DPSD model (Yonelinas, 1994, 1999, 2001) was developed to dissociate the contributions of recollection and familiarity on a memory test, and the meta-*d*’ model (Maniscalco and Lau, 2014) was developed to model metacognition. Surprisingly, direct comparisons between dual process and metacognitive signal detection models of memory confidence have rarely been conducted within the same study (but see Clos *et al*., 2019). Before discussing these models, we first provide a brief overview of signal detection theory.

Signal detection theory has proven to be one of the most influential frameworks in experimental psychology (Macmillan and Creelman, 1991; Wixted, 2020). The main idea is that when one makes a recognition memory decision or any other decision, such as a perceptual decision, they must determine whether a stimulus has actually been observed in the context of an inherently noisy system. Observers make this decision by placing a response criterion whereby stimuli with strength (or evidence) values falling above the criterion are labeled as stimuli that have been observed (i.e., “old” on a memory test), and stimuli that evoke subjective strength below this criterion are labeled as not having been observed (i.e., “new” on a memory test). Response criterion (or response bias) can vary from one individual to another, from one context to another, and even from moment-to-moment. That is, how likely one is to respond “old” to a stimulus can have little to do with actual memory but rather biases in their decision making. For example, one may have a more liberal response bias (i.e., accept a higher number of stimuli as “old”) if the benefits for correctly identifying a target (e.g., $10 reward for every correct response) outweighs or exceeds the costs of incorrectly labeling a lure as “old” (e.g., $1 loss for every incorrect response). Reversing the contingencies in the above example would lead an individual to have a more conservative response bias (Snodgrass and Corwin, 1988; i.e., label only a few stimuli as targets; Koen and Yonelinas, 2011). Although in most cases, targets will evoke greater strength than lures, some targets may evoke weaker strength (e.g., due to poor encoding) and some lures may evoke stronger strength (e.g., due to an imperfect, noisy memory system). Signal detection theory conceptualizes the possible strengths evoked from targets and lures to fall along two Gaussian distributions (Figure Ic), a stronger “signal” distribution (corresponding to the varying degrees of strengths evoked by targets) and a weaker “noise” distribution (corresponding to the varying degrees of strengths evoked by lures). Greater distance between these two distributions is suggestive of better memory (i.e., better ability to discriminate between old and new stimuli), but even with high discriminability, an overly liberal response bias can result in misses and an overly conservative response bias can still result in false alarms. Signal detection theory attempts to identify the discriminability (i.e., the distance or *d*’) between the two distributions of internal evidence corresponding to two different mental states (i.e., memory or no memory for a given stimulus) while accounting for an individual’s response criterion.

Although the signal and noise distributions were originally conceptualized to be of equal variance in the context of episodic recognition memory, the analysis of receiver operator characteristic (ROC) curves suggests otherwise. ROC curves are constructed by plotting the probability of correctly accepting a target (i.e., hit rates) against the probability of incorrectly accepting a lure (i.e., false alarm rates) at multiple levels of response bias (Figures Id and Ie). Note that these are considered type 1 ROC curves because the function is based on the probability of accepting target or lure stimuli based on the number of trials within each class of stimuli (i.e., targets and lures). Multiple levels of response bias are typically obtained by having participants report their confidence in their old/new recognition decisions under the assumption that confidence reflects how participants would respond under different levels of response bias (Figure Ic; Yonelinas and Parks, 2007). This plotting happens cumulatively going from the most stringent criterion (i.e., the highest confidence level that a stimulus is a target, which will mostly be used for targets but not lures) to the most lax criterion (i.e., the highest confidence that a stimulus is a lure). In animals, discrete confidence bins can be obtained by examining the distribution of the time that they wait for a reward before initiating the next trial, a procedure used in the perceptual decision-making literature (e.g., Kepecs *et al*., 2008; Schmack *et al*., 2021)(for a related paradigm, see Sauvage *et al*., 2010). The assumption is that an animal that is confident in their decision will wait longer for a reward because they are expecting their response to be correct. In contrast, an animal will wait less time to initiate another trial when confidence is low, as the opportunity to obtain a reward on the current trial is perceived as unlikely. One can also explicitly manipulate the criterion of animals by varying the reward for correct trials or the difficulty to respond, as has been done in the memory literature (e.g., Fortin *et al*., 2004; Guderian *et al*., 2011). A large body of work has amassed showing that ROC curves from recognition tasks that tap episodic memory in humans (Yonelinas and Parks, 2007), non-human primates (Guderian *et al*., 2011), and rodents (Fortin *et al*., 2004) are typically asymmetrical, suggesting that signal and noise distributions in these data cannot be of equal variance, as is assumed in standard signal detection theory. This asymmetry suggests that performance on recognition memory is driven by at least two processes that influence confidence judgements in old/new memory decisions. There are, of course, other ways of modeling memory confidence such as those involving a single memory process or an attention parameter (Wixted, 2007; Yonelinas and Parks, 2007; for a debate on this issue, see Rotello, 2017), as well as alternative theories that may not even invoke episodic memory (see behaviorism, Skinner, 1991). For this reason, all of the data analyzed here are made available for further analyses under other frameworks (https://osf.io/ehsg4/).

### Dual Process Signal Detection (DPSD) Model

The DPSD model was developed in part to account for the observation that mnemonic ROC curves were asymmetrical (Yonelinas, 1994, 1999). To explain this pattern, a dual process framework was proposed in which familiarity is a continuous signal detection process that reflects the strength of a stimulus, whereas recollection is a threshold process that either occurs or does not (Figure Ic; Yonelinas, 2002; Yonelinas *et al*., 2010). Yonelinas (1994, 1999) proposed a model whereby recognition memory decisions are determined by a mixture of a high-threshold process and an equal-variance signal detection process. The threshold process, which determines the *y*-intercept of the ROC function, is proposed to reflect a recollection process, whereas the curvilinearity of the function is thought to reflect the strength of familiarity-based recognition (Figure Id; Yonelinas, 2001; Koen and Yonelinas, 2010, 2013; cf. Koen *et al*., 2013). These parameters for recollection and familiarity typically converge with those from other non-signal detection methods such as the remember/know and process dissociation procedures (Koen and Yonelinas, 2010; Yonelinas, 2001; but see Rotello *et al*., 2005). Although there has been considerable debate regarding dual vs. single process models (Wixted, 2007; e.g., Ingram *et al*., 2012), different sources of information in memory tend to be captured by recollection and familiarity parameters, respectively, such as context vs. item information, associative vs. unitized information, and hippocampal vs. cortical information (Diana *et al*., 2007; Yonelinas *et al*., 2010; Ranganath and Ritchey, 2012). That is, recollection estimates tend to be greater for memories of scenes or groups of objects, disparate associations such as a separate entity within an environment, and stimuli to which the hippocampus is more responsive (like scenes). In contrast, familiarity estimates are relatively greater for memories of discrete stimuli, stimuli or features that can be grouped into a single entity (like compound words), and stimuli to which the cortex, especially perirhinal cortex, is responsive (like objects).

### Meta-d’ Model

Perhaps the most common way of assessing the correspondence between memory confidence and accuracy (i.e., metamemory) has been with the rank order correlation gamma. In episodic memory research, gamma is the strength of association between levels of memory confidence and correct and incorrect memory decisions. Ideally, one with a good understanding of their own memory system would be highly confident when correct (i.e., responding “old” to targets and “new” to lures) and less confident when incorrect (i.e., responding “new” to targets and “old” to lures). However, as has been reviewed elsewhere (Maniscalco and Lau, 2012; Fleming and Lau, 2014), gamma is an imperfect measure. For one, accuracy in memory performance correlates with gamma. That is, using this measure, those individuals with a poor memory would also tend to have poor metamemory, yet it is possible that one can have a poor memory and recognize their deficit (and similarly, one can have a good memory but not realize this). Some have attempted to control for memory performance while estimating metamemory with gamma (e.g., using attention manipulations during memory encoding to match overall performance between younger and older adults; Wong *et al*., 2012), though the relationship between gamma and memory accuracy remains. Another problem with gamma is that it can be influenced by nonspecific response biases (e.g., using only half of a confidence scale), something that signal detection theory can handle.

To explore metacognition, some researchers have used type 2 signal detection models (Maniscalco and Lau, 2012; Fleming and Lau, 2014). Unlike type 1 ROC curves that plot hit rates as a function of false alarm rates, type 2 ROC curves plot the proportion of correct responses (i.e., *p*[“old”|targets + “new”|lures]) against the proportion of incorrect responses (i.e., *p*[“new”|targets + “old”|lures]) regardless of whether the test stimulus is old or new at different response criteria (or level of confidence; Figure Ie). As with type 1 ROC curves, type 2 ROC curves can be used to compute a discrimination score (i.e., area under the type 2 ROC curve) that quantifies how far the curve is from the chance diagonal. Examining type 2 ROC curves with signal detection theory provides a non-parametric way of mapping the relationship between memory accuracy and confidence which, unlike gamma, accounts for response bias.

Although the area under the type 2 ROC curve is unaffected by type 2 response bias, it is impacted, both mathematically (Galvin *et al*., 2003) and empirically (Higham *et al*., 2009), by type 1 *d*’ and criterion. As an intuition, if the type 1 task is easier (i.e., higher *d*’), it is often also easier to tell when one makes a mistake versus when *d*’ is low and one may always feel as if they are guessing. Therefore, the area under the type 2 ROC curve does not control for type 1 difficulty. A measure with face validity should be able to capture a poor memory of which one is unaware (e.g., many older adults) and a poor memory of which one is aware (e.g., many memory researchers).

A measure called meta-*d*’ was developed to better control for task difficulty (Maniscalco and Lau, 2012, 2014). For a given type 1 *d*’ and criterion, one can generate the type 2 ROC curve that would be expected if all the information available to the type 1 decision is available to metacognition. Similarly, given an observed type 2 ROC curve, one can find the type 1 *d*’ and criterion that would have given rise to the observed type 2 ROC curve. The *d*’ predicted from the type 2 ROC curve alone is referred to as meta-*d*’ and can serve as an index of metacognitive accuracy. Like the area under the type 2 ROC curve, meta-*d’* also correlates with performance (for a similar reason as noted above), but because meta-*d*’ is in units of *d*’, performance can be controlled for by dividing meta-*d*’ by the observed type 1 *d*’. This ratio measure is referred to as metacognitive efficiency (meta-*d*’/*d*’) and captures the degree to which metacognitive judgements discriminate between one’s own correct and incorrect responses relative to how well an ideal metacognitive observer could make such a discrimination given the actual difficulty of the task.

## Drug Effects on Episodic Memory

Before applying the DPSD and meta-*d*’ models to the acquired datasets, we briefly review what is currently known about the effects of different classes of drugs on episodic memory. Surprisingly, there are few reviews summarizing the effects of different psychoactive drugs on episodic memory (but see Kloft *et al*., 2021). When discussing the effects of drugs (or any manipulation) on episodic memory, in addition to specific processes, one must always consider the memory phase that is modulated: encoding, consolidation, or retrieval. As discussed, encoding refers to acquiring or learning information. This is typically the first phase of episodic memory paradigms in which a series of stimuli (e.g., words or pictures) are presented one by one. Consolidation refers to the post-encoding period in which memory traces are thought to be particularly susceptible to interference before they stabilize. Retrieval refers to the act of explicitly remembering and is typically the final phase of memory paradigms in which memory for the encoded stimuli are tested.

Most episodic memory studies with pharmacological manipulations aim to target the encoding phase by administering a drug prior to encoding and testing memory soon after. This approach can easily confound encoding effects with consolidation and retrieval effects given the duration that drugs remain active. However, with careful manipulations, such as separating encoding and retrieval phases with a ≥24-hour delay, certain drug effects can be attributed to different phases. For instance, although the amnestic effect of pre-encoding sedatives can be attributed to both encoding and consolidation, as drug effects persist after the encoding phase into consolidation, sedatives administered specifically at consolidation have the opposite effect (e.g., Doss, Weafer, Ruiz, *et al*., 2018). That is, administering sedatives immediately post-encoding and testing memory ≥24 hours later so that drug effects do not impact memory retrieval actually enhances memory, suggesting that the impairments of pre-encoding sedatives are due to specific modulation of encoding. In another example, THC’s effects on encoding or retrieval impair discrimination between targets and lures on a recognition memory test, but these impairments can be shown to be qualitatively distinct with proper isolation of drug effects to each phase. THC administered before encoding reduces hit rates (typically what is thought to be a memory impairment) with no impact on false alarm rates if a ≥24-hour delay is implemented between encoding and retrieval (Ballard *et al*., 2013). In contrast, THC administered prior to retrieval (with a ≥24-hour delay between encoding and retrieval to not impact consolidation) increases false alarm rates (an increase in response bias potentially due to the induction of false memories; Doss, Weafer, Gallo, *et al*., 2018b).

Whereas an effect on metamemory from a drug administered at retrieval could directly modulate the memory monitoring processes supporting metamemory, an effect on metamemory from drug effects isolated to encoding or consolidation may be less intuitive, as the acute drug effects are no longer apparent during metacognitive decisions (i.e., confidence judgements). When metamemory is impacted from drug effects during encoding or consolidation, one possibility is that memory and metamemory are not completely independent even when controlling for memory performance in metamemory (as is the case with the measure metacognitive efficiency). Another possibility is that knowledge that one was previously on a drug during encoding could impact their expectations and their subsequent confidence judgements. We return to this issue in the Discussion.

### Sedatives (GABA_A_ PAMs)

Although several drugs can be sedating, we reserve “sedative” here for classes of drugs that facilitate activity at the GABA_A_ receptor, specifically GABA_A_ positive allosteric modulators (PAMs). γ-aminobutyric acid (GABA) is the endogenous ligand for this receptor and the main inhibitory neurotransmitter in the brain, so it is perhaps no wonder why drugs that impact the GABA_A_ receptor, such as alcohol, benzodiazepines (drugs typically prescribed for anxiety or insomnia such as alprazolam and triazolam), and non-benzodiazepine hypnotics (or “Z drugs” typically prescribed for insomnia such as zolpidem), are particularly sedating. These different drugs certainly have their pharmacological idiosyncrasies such as binding to different locations on the GABA_A_ receptor, binding to GABA_A_ receptors with different subunits, or binding to other receptors altogether (alcohol is particularly promiscuous). However, as discussed below, they appear to have strikingly similar effects on episodic memory, suggesting that most of these mnemonic effects can be accounted for by GABA_A_ facilitation. More evidence supporting this conjecture could come from studies with other drugs that facilitate GABA_A_ activity (e.g., barbiturates such as amobarbital, quinazolinones such as methaqualone, or even hallucinogenic GABA_A_ agonists such as gaboxadol or the *Amanita muscaria*-derived muscimol). Because the datasets in the present report only include sedative manipulations of encoding and consolidation, we do not discuss retrieval effects, though we note the few studies that have explicitly tested sedative effects on retrieval do not tend to find effects (Curran, 1986).

Sedatives are particularly well-known for their amnestic effects, though such memory impairments are only present when these drugs are administered prior to memory encoding. Several studies using the remember/know procedure initially reported that sedatives administered at encoding impaired recollection but not familiarity (reviewed and reanalyzed in Doss, Weafer, Ruiz, *et al*., 2018). However, in a reanalysis of these data using a statistical correction for the co-occurrence of recollection and familiarity (the independence remember/know procedure; Yonelinas and Jacoby, 1995), it was found that most of these studies did in fact find sedatives to impair both recollection and familiarity. Moreover, this study included a DPSD analysis of separate confidence data, which also found alcohol to impair both recollection and familiarity, highlighting the convergence of the remember/know and DPSD methods (Yonelinas, 2001; Koen and Yonelinas, 2010). Additionally, GABA_A_ PAMs at encoding were shown to impair memory of both detail and gist for stories (Kamboj and Curran, 2006) and both item and context memory (Reder *et al*., 2013). Together, these global encoding impairments provide some explanation for why sedatives can cause “blackouts.”

Several studies have found that sedatives at encoding impair metamemory (Bacon *et al*., 1998; Mintzer and Griffiths, 2003a; b, 2007; Carter *et al*., 2009; Mintzer *et al*., 2010; Kleykamp *et al*., 2012; Carter, Kleykamp, *et al*., 2013; Carter, Reissig, *et al*., 2013), though not always (Mintzer and Griffiths, 2005; Izaute and Bacon, 2006; Kleykamp *et al*., 2010). However, all of these studies used gamma and are therefore susceptible to response bias confounds and memory accuracy scaling with this measure of metamemory. Considering the massive amnestic effects of these drugs, it should be unsurprising that a measure of metamemory that is correlated with memory accuracy would also exhibit a reduction.

Perhaps one of the most paradoxical and lesser known effects in psychopharmacology is that, despite causing dense amnesia when sedatives are administered at encoding, sedatives actually enhance memory if administered immediately post-encoding (i.e., during consolidation), an effect referred to as “retrograde facilitation” (Wixted, 2004; Mednick *et al*., 2011). Retrograde facilitation has been found by multiple groups, from different sedatives (alcohol, benzodiazepines, and zolpidem), on various stimuli (e.g., words, word pairs, pictures, word-picture pairs, emotional and neutral material), with distinct memory tests (e.g., recall, recognition, cued recollection), and across various delays between encoding and retrieval (Hinrichs *et al*., 1984; e.g., same day encoding/retrieval, sleep after encoding, one- and two-day delays between encoding and retrieval; e.g., Bruce and Pihl, 1997; Fillmore *et al*., 2001; Kaestner *et al*., 2013; Mednick *et al*., 2013; Weafer *et al*., 2016a). One explanation of retrograde facilitation is that these drugs prevent the encoding of new information that would normally retroactively interfere with information previously encoded in a drug-free state while having no effect on memory stabilization. In contrast, other amnestic drugs, such as γ-hydroxybutyric acid (GHB, a GABA_B_ agonist) and THC do not cause retrograde facilitation (Parker *et al*., 1980; Mednick *et al*., 2013), suggesting that GHB and THC impair memory stabilization processes or that sedatives may actively facilitate such consolidation processes. Evidence for the latter comes from studies that administered zolpidem immediately post-encoding followed by a nap (Kaestner *et al*., 2013; Mednick *et al*., 2013; Niknazar *et al*., 2015). These studies found that post-encoding zolpidem enhanced the power of slow waves and the occurrence of sleep spindles, processes important for memory consolidation (Latchoumane *et al*., 2017). Moreover, greater coupling between these oscillations was associated with larger memory enhancements of zolpidem at consolidation on recollection-based memory (Niknazar *et al*., 2015; Latchoumane *et al*., 2017). One DPSD analysis found alcohol at consolidation to enhance both recollection and familiarity, though this effect was mostly only apparent for neutral and not emotionally positive or negative stimuli (Doss, Weafer, Ruiz, *et al*., 2018; but see, Bruce and Pihl, 1997; Kaestner *et al*., 2013 for studies that found enhancements of emotional stimuli). In contrast, no study has measured metamemory in a retrograde facilitation paradigm.

### Dissociatives (NMDA Antagonists)

Dissociative hallucinogens (or simply “dissociatives”) are drugs that antagonize the *N*-methyl-D-aspartate (NMDA) receptor. NMDA receptors are widely distributed throughout the brain, and they are involved in synaptic learning mechanisms (i.e., long-term potentiation and depression). Conditions in which the function of these receptors are attenuated (i.e., anti-NMDA receptor encephalitis) can result in psychosis, and similarly, dissociatives produce effects that can model both positive and negative symptoms of psychosis (Krystal, 1994). Although these drugs are used recreationally in social settings (e.g., clubs, raves) in lower doses, at higher doses, they can induce an anesthetic state of high dissociation referred to as a “K-hole” (referring to this being a particularly common phenomenon of high doses of ketamine; Muetzelfeldt *et al*., 2008). Here, we discuss the dissociatives ketamine (an anesthetic used particularly in populations with respiratory and cardiac problems) and dextromethorphan (a medicine found in over-the-counter cough syrups). Although the effects of these drugs are likely not mediated solely by NMDA antagonism, the similar dissociative state evoked by these drugs, as well as other NMDA antagonists such as nitrous oxide (i.e., “laughing gas”), phencyclidine (or PCP), and emerging novel compounds like methoxetamine (or MXE) suggest that NMDA antagonism is a primary mechanism of action of these drugs. Because the present report only contains reanalysis of studies in which dissociatives were administered at encoding, we focus our discussion of these drugs on this phase. Nevertheless, it has been proposed that post-encoding NMDA antagonism (i.e., during consolidation) should cause retrograde facilitation like sedatives (Wixted, 2004; Mednick *et al*., 2011), though one study suggests if anything dissociatives may impair memory consolidation (Das *et al*., 2016). Moreover, studies on the effects of dissociatives at retrieval have administered drugs shortly after encoding and tested memory shortly after, thereby conflating consolidation and retrieval effects. These studies find dissociatives to have little impact on memory accuracy (Honey *et al*., 2005) with some evidence of diminished metamemory (Lehmann *et al*., 2021).

Because NMDA receptors are involved in synaptic learning, these drugs have strong amnestic effects when administered prior to encoding like sedatives. Whereas memory tasks requiring recollection, such as free recall, find clear impairments of dissociatives at encoding (Lofwall *et al*., 2006; Carter, Kleykamp, *et al*., 2013; Carter, Reissig, *et al*., 2013; Barrett *et al*., 2018), it is less clear if familiarity is also impacted. One study administered ketamine prior to encoding and used the remember/know procedure at retrieval (Hetem *et al*., 2000). Although ketamine was found to impair both recollection and familiarity, the independence correction (Yonelinas and Jacoby, 1995) was not performed, and the recognition test contained no lures. Another study used the DPSD model and found some evidence for dextromethorphan at encoding to impair familiarity (Barrett *et al*., 2018), but the modeling approach may have produced unreliable estimates because of low trial counts. It is also unclear whether these dissociatives at encoding impact metamemory, as some studies found no impact (Lofwall *et al*., 2006; Carter, Kleykamp, *et al*., 2013; Lehmann *et al*., 2021), whereas others found impairments of metamemory with higher doses of dextromethorphan (Carter, Reissig, *et al*., 2013). All of these studies measured metamemory with gamma except (Lehmann *et al*., 2021), which did measure metamemory with metacognitive efficiency (i.e., from the meta-*d*’ model) and did not find any impact of ketamine at encoding on metamemory. Thus, as was the case for sedatives, an impairment in metamemory may be an artefact of the covariation between memory accuracy and metamemory measured with gamma.

### Psychedelics (5-HT_2A_ Agonists)

Although research into the effects of psychedelics in humans has seen a recent resurgence, surprisingly little work has investigated their effects on episodic memory. Psychedelics have a range of effects on perception, particularly vision (Kometer and Vollenweider, 2016), and compared to dissociatives, the effects of psychedelics more closely resemble positive rather than negative symptoms of psychosis. Moreover, psychedelics appear to facilitate or distort semantic processing (Spitzer *et al*., 1996), which may be responsible for the oft-reported sense that some experiences are endowed with highly significant meaning (Griffiths *et al*., 2006). Although some use “psychedelic” to refer to other hallucinogens, we reserve the term here for drugs with serotonin 2A (5-HT_2A_) agonism such as psilocybin (derived from *Psilocybe* species of “magic mushrooms”), lysergic acid diethylamide (LSD), and *N*,*N*-dimethyltryptamine (DMT; found in the brew ayahuasca that is made orally active via monoamine oxidase inhibitors). When co-administered with a 5-HT_2A_ antagonist, all of these drugs produce few subjective effects (Vollenweider *et al*., 1998; Valle *et al*., 2016; Preller *et al*., 2018). Similarly, the *R*-enantiomer of ±3,4-methylenedioxymethamphetamine (MDMA) is a 5-HT_2A_ agonist, and co-administration of a 5-HT_2A_ antagonist attenuates the subjective effects (Liechti, 2000), as well as the mnemonic effects of MDMA (van Wel *et al*., 2011). Moreover, when the isolated *R*-enantiomer of a similar drug (3,4-methylenedioxyethylamphetamine or MDEA) has been administered in humans, it appears to produce psychedelic effects distinct from the *S*-enantiomer (Spitzer *et al*., 2001). For these reasons, here, we group MDMA with psychedelics. Although we recognize that racemic MDMA also has prominent stimulant and entactogenic effects (owing to the *S*-enantiomer’s affinity for catecholamine and serotonin transporters, respectively), as will be discussed, the effects of MDMA on episodic memory resemble those of psychedelics. We limit the discussion of psychedelics to encoding and retrieval effects, as the reanalyzed data from studies of psychedelics in the present report only manipulated these two studies. Nevertheless, one study in humans (Wießner *et al*., 2022) and one study in mice (Zhang *et al*., 2013) found evidence for post-encoding psychedelics to enhance memory that may be particularly familiarity-based.

One recent study explored the effects of psilocybin on episodic memory encoding, and a few studies have looked at MDMA’s effects on memory encoding. Psilocybin administered prior to encoding was found to impair verbal free recall, and a DPSD analysis found some evidence for an impairment of recollection with higher doses, but simultaneously, familiarity was numerically enhanced (Barrett *et al*., 2018). As discussed previously, the analysis of this study likely suffered from poor model fits due to low trial counts. Studies have consistently found MDMA at encoding to impair free recall (Kuypers and Ramaekers, 2005; Kuypers *et al*., 2008, 2011, 2013; van Wel *et al*., 2011; de Sousa Fernandes Perna *et al*., 2014), even though typical stimulants can enhance memory encoding (Soetens *et al*., 1995; Zeeuws *et al*., 2010; Linssen *et al*., 2012). Moreover, this impairment of MDMA can be abolished with 5-HT_2A_ antagonism (van Wel *et al*., 2011), suggesting that despite MDMA’s stimulant effects (i.e., facilitating catecholaminergic transmission), it acts more like a psychedelic in terms of its effects on episodic memory. Further evidence for this assertion comes from a DPSD analysis that found MDMA to impair recollection with a trend toward enhanced familiarity (Doss, Weafer, Gallo, *et al*., 2018a), though these effects were only found for emotional (negative and positive) but not neutral stimuli. This study also implemented a subsequent recognition test with the remember/know procedure and found further evidence for impaired recollection. Despite psychedelic users anecdotally reporting that these drugs give a better understanding of one’s mind (the word psychedelic means “mind manifesting”), psychedelics have not been tested on any measure of metacognition.

Only one study has isolated the effects of a psychedelic to memory retrieval (Doss, Weafer, Gallo, *et al*., 2018a; for studies using non-objective measures, see Carhart-Harris *et al*., 2012, 2014). Using the DPSD model, MDMA numerically impaired recollection but not familiarity, and thus, these findings were interpreted as MDMA potentially inhibiting the retrieval of details from memory. However, there did not appear to be any impact on recollection measured from the remember/know procedure, and MDMA at retrieval somewhat increased high confidence false alarms, an effect that can particularly reduce recollection estimates of the DPSD model. With more false alarms using the highest level of confidence, the first point of a ROC curve is plotted further away from the *y*-axis, thereby pulling the *y*-intercept of a DPSD ROC curve (i.e., the recollection estimate) toward zero. This known zero-bias of the DPSD model is typically not an issue, as there are rarely high confidence false alarms, though manipulations of false memory (e.g., the Deese-Roediger-McDermott task, Roediger and McDermott, 1995) can produce bizarre results like zero estimates of recollection, a finding exhibited by hippocampal amnesic patients (Yonelinas *et al*., 1998). Such findings suggest that when false memories are high, different task and modeling procedures are necessary to dissociate other processes (e.g., the “phantom ROC” model to estimate false memory parameters; Lampinen *et al*., 2006).

### Stimulants (Catecholamine Transporter Inhibitors)

Although the term “stimulant” can be used for drugs like caffeine, modafinil, and nicotine, here we reserve this term for drugs whose major effects stem from their inhibition of catecholamine (i.e., dopamine and norepinephrine) transporters such as amphetamine (specifically the dextro-/*S*-enantiomer, which is far more potent than the levo-/*R*-enantiomer), methamphetamine (also the dextro-/*S*-enantiomer), cocaine, methylphenidate, and cathinones (or “bath salts,” amphetamines with a ketone group). Amphetamine and methylphenidate are prescribed for attention deficit hyperactivity disorder (ADHD), as they appear to enhance various aspects of cognition, and methamphetamine is still a prescribed drug for obesity despite its high abuse potential. The pharmacology of these stimulants vary slightly with some of these drugs (e.g., dextroamphetamine, dextromethamphetamine, and cocaine) also inhibiting the serotonin transporter to a lesser degree, though they mostly lack entactogenic effects like MDMA. Furthermore, many of the amphetamines not only block monoamine transporters but also reverse their action that causes a release of neurotransmitters from presynaptic neurons.

Stimulants at encoding can enhance recollection-based memory (e.g., measured with free recall), especially when there is a longer delay between encoding and retrieval (Soetens *et al*., 1995; Zeeuws *et al*., 2010; Linssen *et al*., 2012). However, enhancement of memory encoding or other cognitive processes with these drugs has been somewhat inconsistent, with cognitively impaired populations (e.g., patients with ADHD) sometimes seeing more of a benefit (Advokat, 2010; Bagot and Kaminer, 2014). Because these drugs tend to prevent sleep, subsequent memory enhancements can be canceled out by impaired sleep (Ballard *et al*., 2015; Whitehurst and Mednick, 2020). The effects of stimulants at encoding on familiarity or metamemory have not been tested, though some work found that the impairments of metamemory (measured with gamma) from GABA_A_ PAMs at encoding can be attenuated by co-administration of dextroamphetamine (Mintzer and Griffiths, 2003b, 2007).

Because encoding enhancements from stimulants are more prominent after a delay, this would suggest that stimulants enhance post-encoding consolidation processes. Although few human studies have directly manipulated consolidation with stimulants (by administering drugs immediately post-encoding and testing memory after a delay), one possibility is that stimulants at consolidation enhance memory via dopamine-dependent replay (Gruber *et al*., 2016). Administration of stimulants after animals learn various behaviors can enhance learning (Krivanek and McGaugh, 1969; Janak and Martinez, 1992; Simon and Setlow, 2006), though these forms of memory are not thought to be episodic-like. One study found post-encoding intramuscular dextroamphetamine to enhance free recall on a test one day later, but there was an initial memory test shortly after drug administration that could have served as another instance of memory encoding (Soetens *et al*., 1995). Thus, this second instance of memory encoding may have actually been enhanced rather than memory consolidation. In contrast, a direct test of memory consolidation with post-encoding dextromethamphetamine found no effect on a picture recognition test (Ballard *et al*., 2015). Thus, it is unclear what effects, if any, stimulants at consolidation have on recollection, familiarity, or metamemory.

Most studies on the effects of stimulants at retrieval are in the context of state-dependent learning (i.e., facilitation of memory because of being in the same state during encoding and retrieval). Like MDMA, one study found dextroamphetamine at retrieval to increase false memories (Ballard *et al*., 2014), but another study did not replicate this effect (Weafer *et al*., 2014). Studies of state-dependency are mixed, sometimes finding that stimulants at retrieval on a delayed test with sober encoding conditions or vice versa can result in reductions of recollection-based memory (Bustamante *et al*., 1970; Shea, 1982), but others finding no such evidence (Aman and Sprague, 1974; Steinhausen and Kreuzer, 1981; Weingartner *et al*., 1982; Becker-Mattes *et al*., 1985). Finally, the effects of stimulants at retrieval or both encoding and retrieval on familiarity or metamemory have not been tested.

### Cannabinoids (CB_1_ Agonists)

Although all compounds in cannabis (e.g., cannabidiol) can be referred to as “cannabinoids,” here we reserve this term for agonists of the cannabinoid 1 receptor (CB_1_), specifically THC. THC and synthetic CB_1_ agonists (e.g., the medication nabilone and “spice” drugs) have subjective and perceptual effects that can mimic psychotic symptoms, sometimes resulting in these drugs being labeled as hallucinogens. Despite recent ongoing studies looking into the utility of these drugs in the treatment of a plethora of conditions (e.g., pain, insomnia, anxiety), the evidence for clinical utility is still rather limited, and currently, the only approved use of CB_1_ agonists is for nausea and appetite stimulation in patients with cancer and acquired immunodeficiency syndrome (i.e., AIDS). We focus the discussion of cannabinoids on encoding and retrieval effects, as the data analyzed in the present report have no consolidation manipulation. We note, however, that one study found no impact of THC at consolidation on free recall (Parker *et al*., 1980).

THC is known to be amnestic when administered at encoding. Such studies tend to find impairments in verbal free recall, suggesting that recollection is impaired (Miller and Cornett, 1978; Hart *et al*., 2010; Ranganathan *et al*., 2017). However, because these studies do not always find impairments in recognition, THC administered during encoding might not impact familiarity. It is unclear what THC at encoding does to metamemory, though one study found THC to impair metacognition on a working memory task in which metacognition was derived by correlating the number of correct responses on each trial (out of six) with the number of high confident responses on each trial (Adam *et al*., 2020). This measure of metamemory likely suffers from the same problems as gamma, namely response bias and scaling with performance.

Interestingly, THC was previously thought to not impact memory retrieval (Curran *et al*., 2016) because in contrast to THC at encoding, hit rates and verbal free recall appear to be unaffected when THC is administered strictly at retrieval (Ranganathan *et al*., 2017; Doss, Weafer, Gallo, *et al*., 2018b). However, recently it was rediscovered (see Abel, 1971) that that THC at retrieval robustly increases false memories as measured by increased high confidence false alarms on a cued recollection test and false recognition with “remember” and “know” responses on a recognition test (Doss, Weafer, Gallo, *et al*., 2018b). One reason this effect was somewhat evasive was that in many studies, cannabinoids were administered prior to encoding and memory was tested soon after encoding (i.e., drug effects on both encoding and retrieval; Ilan *et al*., 2004; Hart *et al*., 2010). Such a study design can result in both reductions in hit rates and elevations in false alarm rates and thus, overall attenuations of corrected memory scores (e.g., hits minus false alarms and *d*’). Although this pattern of memory performance can be exhibited when memory is near chance, memory is not typically at chance in these studies, suggesting that these are qualitatively different memory errors to which signal detection models are agnostic. Such an increase in false memories might implicate an impairment of metamemory, though it may simply be reflected as a criterion shift in signal detections models.

## Analytic Methods

Data from 10 separate double-blind, placebo-controlled drug administration studies were re-analyzed to determine the effects of a variety of drug classes on recollection, familiarity, and metamemory (Table I). For estimating recollection and familiarity from the DPSD model, we used the ROC Toolbox for Matlab (Koen *et al*., 2017). This toolbox plots hit rates against false alarm rates cumulatively, going from the most stringent criterion (i.e., highest confidence “old” response) to the most lax criterion (i.e., lowest confidence “new”). Then the DPSD model is fit to the observed data using maximum likelihood estimation to obtain estimates of recollection (i.e., the *y*-intercept) and familiarity (i.e., the curvature of the function modeled as an equal-variance signal detection process).

**Table I.** *Reanalyzed Datasets* MDMA = ±3,4-methylenedioxymethamphetamine, THC = Δ^9^-tetrahydrocannabinol.

| Study | Original publication(s) | Drug(s) | Targeted Phase(s) |
| --- | --- | --- | --- |
| 1 | (Weafer *et al.*, 2016a; b) | alcohol | encoding, consolidation |
| 2 | (Kleykamp *et al.*, 2012) | zolpidem | encoding |
| 3 | (Carter, Kleykamp, *et al.*, 2013) | triazolam, ketamine | encoding |
| 4 | (Carter, Reissig, *et al.*, 2013) | triazolam, dextromethorphan | encoding |
| 5 | (Barrett *et al.*, 2018) | dextromethorphan, psilocybin | encoding |
| 6 | (Doss, Weafer, Gallo, *et al.*, 2018a) | MDMA | encoding, retrieval |
| 7 | (Weafer *et al.*, 2014) | dextroamphetamine | encoding, retrieval, both<br>encoding and retrieval |
| 8 | (Ballard *et al.*, 2015) | dextromethamphetamine | encoding, consolidation |
| 9 | (Doss, Weafer, *et al.*, 2020) | THC | encoding |
| 10 | (Doss, Weafer, Gallo, *et al.*, 2018b) | THC | retrieval |

Whereas studies estimating single subject parameters usually implement tasks with more than 100 targets and lures per condition (e.g., Koen *et al*., 2013), some of the datasets analyzed contained as few as 36 trials per drug condition. Low trial counts can obscure drug effects (e.g., THC not impairing working memory, Adam *et al*., 2020), and this can be particularly problematic with computational models including meta-*d*’ (Fleming, 2017). Additionally, repeated measures studies with missing data points (e.g., when a dose was too strong for a given participant) would require excluding all of a participant’s data, thereby reducing power. Thus, we performed the following bootstrapping procedure (Lampinen *et al*., 2006; Doss, Weafer, Gallo, *et al*., 2018a; Kim *et al*., 2019). For a given condition within a dataset with *N* participants, we randomly drew data from *N* participants with replacement and fit each model. This procedure was repeated 10,000 times to generate distributions of recollection and familiarity parameters. Comparisons between conditions can then be made by subtracting distributions and inspecting whether the 95% confidence interval (CI) does not contain 0 or whether 95% of the difference distribution lies above or below 0 (for α = .05, one-tailed). Because our intent was to explore unique patterns produced by different classes of drugs that could guide future work, we did not correct for multiple comparisons, and we discuss recurring trends, especially those that may be unique to a class of drugs.

For estimating metamemory from the meta-*d*’ model, we used meta-*d*’ Matlab functions that are described elsewhere (Maniscalco and Lau, 2012). These functions plot the cumulative proportion of correct against incorrect responses going from the most stringent criterion (i.e., highest confidence correct response) to the most lax criterion (i.e., lowest confidence incorrect response). A curve is then fit to these points using maximum likelihood estimation. From this type 2 ROC curve, a type 1 *d*’ can be interpolated assuming a metacognitively ideal observer (i.e., perfect correspondence between confidence and accuracy). This particular type 1 *d*’ is referred to as meta-*d*’. Because meta-*d*’ can scale with one’s actual memory accuracy, metacognitive efficiency is calculated by taking the ratio of meta-*d*’ to *d*’. Values closer to 1 therefore indicate more efficient metacognition (less loss of information between the type 1 and type 2 response). Unless otherwise stated, we refer to metacognitive efficiency as metamemory. We performed the same bootstrapping procedure described above using identical random samples of participants.

Each drug condition was individually modeled and compared separately to the placebo condition of a given dataset. For simplicity, all stimulus conditions (e.g., negative, neutral, and positive emotional stimuli) were collapsed (i.e., confidence counts were aggregated) before fitting models. Parameter distributions and statistics from models fit to separate stimulus conditions can be found in the Supplemental Material (SM) and are discussed when relevant. All datasets analyzed here can be found at https://osf.io/ehsg4/.

### Study 1: The Effects of Alcohol on Encoding and Consolidation

The dataset from the current analysis comes from (Weafer *et al*., 2016a; b) and was subsequently submitted to a DPSD analysis (Doss, Weafer, Ruiz, *et al*., 2018). In this reanalysis, alcohol at encoding impaired both recollection and familiarity, though these effects were mostly only apparent for negative and positive stimuli but not neutral stimuli (Doss, Weafer, Ruiz, *et al*., 2018). Impairments of both recollection and familiarity were supported by a reanalysis of remember/know data from several studies with GABA_A_ PAM manipulations at encoding that had previously failed to use the independence correction to derive proper familiarity estimates (Yonelinas and Jacoby, 1995). Although a meta-*d*’ analysis was not previously run on the current dataset, several studies with GABA_A_ PAM manipulations at encoding have found metamemory impairments on verbal recognition tests using gamma (Mintzer and Griffiths, 2003b; a, 2007; Carter *et al*., 2009; Mintzer *et al*., 2010; Kleykamp *et al*., 2012; Carter, Kleykamp, *et al*., 2013; Carter, Reissig, *et al*., 2013). As discussed, not only is gamma problematic as a measure of metacognition (Maniscalco and Lau, 2012; Fleming and Lau, 2014), these studies tested memory within the same day, thereby not controlling for potential drug effects on memory retrieval. Moreover, drug-induced amnesia can become larger after a delay, during which time recognition of a memory impairment could become more apparent to an individual. This might especially be true of the present sample, who were social drinkers that have potentially acquired metacognitive knowledge regarding the effects of alcohol on their memory. Thus, it is possible that longer delays between encoding and retrieval may attenuate metamemory impairments.

Whereas alcohol and other GABA_A_ modulators at encoding are known to impair memory, GABA_A_ modulators administered immediately post-encoding (i.e., at consolidation) enhance memory (i.e., retrograde facilitation). Previously, it was found in the current dataset that alcohol at consolidation enhanced both recollection and familiarity, though these effects were mostly only apparent for neutral material (Doss, Weafer, Ruiz, *et al*., 2018). Others have also found post-encoding alcohol or zolpidem to enhance memory on recollection-based tasks (i.e., free recall, paired-associates; Bruce and Pihl, 1997; Knowles and Duka, 2004; Mednick *et al*., 2013). In contrast to recollection and familiarity, it is unclear what the effects of alcohol at consolidation will be on metamemory. Because retrograde facilitation is a relatively unknown phenomenon, alcohol users might be unaware of any memory enhancements from post-encoding alcohol. In such a scenario, metamemory, as measured by metacognitive efficiency, might be attenuated.

#### Study Methods

This study is described in-depth elsewhere (Weafer *et al*., 2016a; b). Briefly, a double-blind, between-subjects design was used in which 59 young adults (*M* = 24.1-years-old, range = 21-30) were randomized to one of 3 demographically similar groups: placebo (*N* = 19, 11 males), encoding (*N* = 20, 11 males), or consolidation (*N* = 20, 11 males). The placebo group received placebo prior to encoding stimuli and placebo immediately post-encoding (i.e., at consolidation). The encoding group received alcohol prior to encoding and placebo at consolidation. The consolidation group received placebo prior to encoding and alcohol at consolidation. Doses of alcohol (.7 and .8 g/kg for females and males, respectively) and placebo were administered orally via gelatin.

Participants came in for 2 sessions separated by 48 hours. During the first session, participants consumed the first serving of gelatin and 20 minutes later completed the encoding phase of a 240-trial cued recollection task. This encoding phase consisted of the presentation of emotional (negative, neutral, positive) and beverage-related (non-alcoholic, alcoholic) labels (48 of each condition), half of which were followed by a picture of that label. During the presentation of each label, participants were to rate on a five-point scale how much they would like to see a picture of the label. On trials in which a picture was presented, they were to rate the picture’s valence on a five-by-five grid with negative and positive valence ratings on orthogonal axes and arousal on a five-point scale. After the encoding phase, participants consumed the second serving of gelatin and remained in the lab for three and half hours. To minimize retroactive interference, participants were only permitted to listen to music without lyrics during the first two hours. During the second session, participants’ memory for the pictures from the encoding phase was tested (under sober conditions) using a cued recollection test. Participants were presented with all 240 labels in random order and asked whether they had seen a picture of each label (yes/no) followed by a confidence rating (5-point scale). Therefore, labels for which a picture had been presented were targets and labels presented without a corresponding picture were lures.

#### Results

Figures IIa-c display the distributions of parameter estimates from the DPSD and meta-*d*’ models. As previously found, compared to placebo, alcohol at encoding impaired both recollection (*M* = .16, *SD* = .04, CI = [.08, .24], *p* < .001) and familiarity (*M* = .28, *SD* = .11, CI = [.06, .49], *p* = .007), and consistent with past work, alcohol at encoding impaired metamemory (*M* = .16, *SD* = .09, CI = [-.02, .34], *p* = .047). However, this effect was driven by the beverage stimuli (see SM), which have much more overlapping conceptual and perceptual features than the negative, neutral, and positive stimuli. In contrast to the original analyses of these data separated by stimulus type, alcohol’s enhancement of consolidation was only trending for recollection (*M* = .07, *SD* = .05, CI = [-.03, .16], *p* = .072) and negligible for familiarity (*M* = .06, *SD* = .10, CI = [-.13, .26], *p* > .250). This reduction of retrograde facilitation compared to the original study was because enhancements of recollection and familiarity were only found for neutral material (see SM). Alcohol at consolidation had no significant impact on metamemory (*M* = .03, *SD* = .12, CI = [-.22, .25], *p* > .250).

**Figure II.**
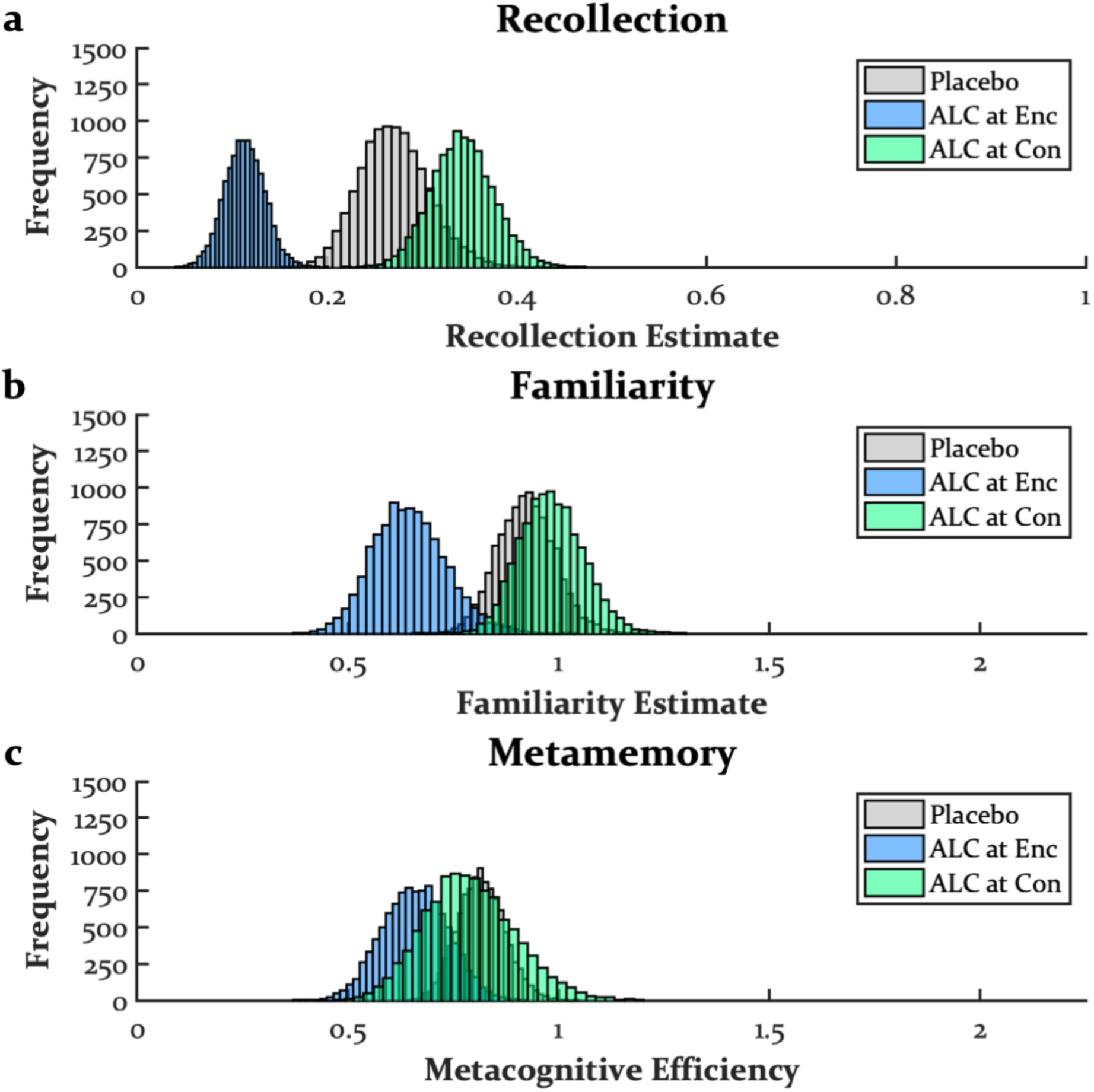
*The Effects of Alcohol at Encoding and Consolidation* Alcohol at encoding impaired recollection, familiarity, and metamemory, though the metamemory impairment may have been due to idiosyncratic stimuli (see Supplemental Material and Discussion), and this impairment did not replicate across sedatives. Alcohol at consolidation enhanced recollection but had no impact on familiarity or metamemory. Doses used here were .7 and .8 g/kg for females and males, respectively. ALC = alcohol, Enc = encoding, Con = consolidation.

### Study 2: The Effects of Zolpidem on Encoding

The dataset from the current analysis comes from (Kleykamp *et al*., 2012). Like alcohol, zolpidem is a GABA_A_ PAM. In the original analysis of these data, zolpidem at encoding impaired memory on verbal free recall tests (i.e., a recollection-based task), as well metamemory on verbal recognition tests measured with gamma. Based on previous work (Doss, Weafer, Ruiz, *et al*., 2018) and the previous analysis, it would be expected for zolpidem at encoding to impair recollection, familiarity, and perhaps metamemory.

Although the complex design of the current dataset (outlined below) has several potential confounds (e.g., drug effects on retrieval, practice effects, interactions between drug effects and circadian rhythms, withdrawal effects), it also has many advantages because of the implementation of multiple memory tests within a session. For example, the second memory test assesses drug effects on encoding when there is a short delay between encoding and retrieval, and the third memory test allows for the assessment of drug effects with a longer delay between encoding and retrieval. Whereas impairments on memory encoding can sometimes become larger with longer delays between encoding and retrieval, it is possible that metamemory impairments may become smaller because larger memory impairments and more time could better allow one to realize the degree of their impairment. Additionally, because the zolpidem used in this study was an extended-release formulation, the fourth memory test could act as a lower dose condition because of lingering drug effects. Finally, there was a period of several days between the two zolpidem administrations in which participants self-administered zolpidem at home each night. This allows for the potential development of tolerance to zolpidem between these sessions, in which case the second zolpidem administration could also act as a lower dose.

#### Study Methods

This study is described in-depth elsewhere (Kleykamp *et al*., 2012). Briefly, a double-blind, within-subjects design was used in which 15 young adults (*M* = 30.0-years-old, range = 21-42, all males) came in for 4 sessions that each involved the oral administration of a capsule containing placebo or 12.5 mg of extended-release zolpidem. During the first and fourth sessions, participants orally consumed a capsule containing placebo, and during the second and third sessions, they orally consumed a capsule containing zolpidem. Between sessions two and three, participants took zolpidem nightly for 22 to 30 days.

Each session contained four verbal memory tests. During the encoding phase of each memory test, participants were presented with 36 concrete nouns, each of which they were to read aloud. During the retrieval phase, participants first completed a free recall test followed by a recognition test, which contained the 36 previously presented words (targets) and 36 new words (lures). Participants used a single response, hybrid old/new six-point confidence scale (i.e., definitely old, probably old, maybe old, maybe new, probably new, definitely new) to indicate whether each word was previously presented. The first memory test was always conducted prior to drug administration with the encoding and retrieval phases separated by 60 minutes during which other cognitive tasks were completed. Afterward, participants consumed a capsule, and 30 minutes later, they went to sleep. At 80 minutes post-capsule administration, the lights were turned on, and participants were awoken. At 90 minutes post-capsule administration, they completed the second memory test, again with the encoding and retrieval phases separated by 60 minutes of other cognitive tasks. After the retrieval phase of the second memory test, participants completed the encoding phase of the third memory test. At 180 minutes post-capsule administration, participants went back to sleep for another 210 to 270 minutes. At 390 minutes post-capsule administration, the lights were turned back on, and at 450 minutes post-capsule administration, participants completed the retrieval phase of the third memory test, followed by the fourth memory test, again with the encoding and retrieval phases separated by 60 minutes of other cognitive tasks. We separately compared both zolpidem sessions to both placebo sessions for each memory test at a given timepoint, producing four contrasts. See SM for results of the first memory test (i.e., prior to capsule administration) and results of the fourth memory test (i.e., when there was little to no effect of zolpidem on memory the morning after capsule administration).

#### Results

Figures IIIa-c display the distributions of parameter estimates from the DPSD and meta-*d*’ models for the second memory test when the effects of zolpidem were near peak, and memory retrieval occurred shortly after encoding. Three of the four contrasts found zolpidem at encoding to impair recollection (first placebo vs. first zolpidem: *M* = .13, *SD* = .08, CI = [-.03, .27], *p* = .048; first placebo vs. second zolpidem: *M* = .09, *SD* = .08, CI = [-.09, .24], *p* = .147; second placebo vs. first zolpidem: *M* = .29, *SD* = .07, CI = [.15, .42], *p* < .001; second placebo vs. second zolpidem: *M* = .24, *SD* = .07, CI = [.10, .38], *p* = .001). Because drug conditions were always administered in the same order and participants took zolpidem for 22 to 30 weeks before their second zolpidem sessions, practice effects and drug tolerance may account for the non-significant finding between the first placebo and second zolpidem sessions. All contrasts found zolpidem at encoding to impair familiarity (first placebo vs. first zolpidem: *M* = .71, *SD* = .12, CI = [.48, .94], *p* < .001; first placebo vs. second zolpidem: *M* = .69, *SD* = .13, CI = [.45, .94], *p* < .001; second placebo vs. first zolpidem: familiarity: *M* = .57, *SD* = .17, CI = [.23, .91], *p* < .001; second placebo vs. second zolpidem: *M* = .56, *SD* = .17, CI = [.23, .91], *p* < .001). In contrast, none of the contrasts found zolpidem at encoding to impair metamemory (first placebo vs. first zolpidem: *M* = .34, *SD* = .57, CI = [-.55, 1.65], *p* > .250; first placebo vs. second zolpidem: *M* = .67, *SD* = 1.42,CI = [-.41, 3.02], *p* = .192; second placebo vs. first zolpidem: *M* = .36, *SD* = .59, CI = [-.51, 1.73], *p* > .250; second placebo vs. second zolpidem: *M* = .69, *SD* = 1.46 CI = [-.48, 3.23], *p* = .213), though these statistics should be interpreted with caution as the distributions were extremely wide (and therefore not depicted in Figure IIIc).

**Figure III.**
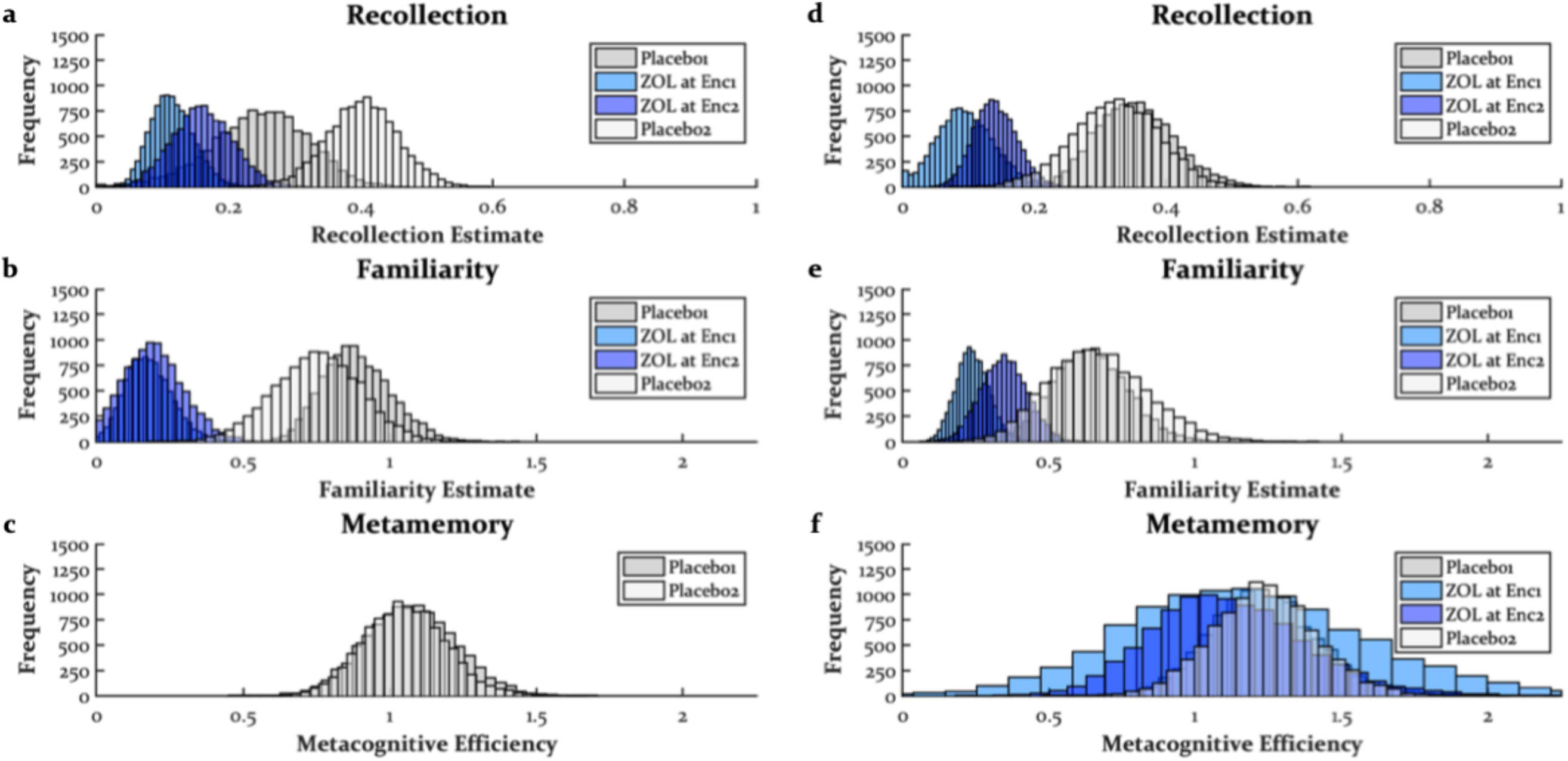
*The Effects of Zolpidem at Encoding* Zolpidem (12.5 mg extended-release) at encoding impaired recollection and familiarity but had no impact on metamemory. Ordering in legend is the sequence in which placebo/zolpidem sessions took place, and the numbers in legend indicate the first and second placebo/zolpidem sessions. Both metamemory distributions for zolpidem on the first memory test (panel c) were too wide to plot on the scale common to all plots. ZOL = zolpidem, Enc = encoding.

Figures IIId-f display the distributions of parameter estimates from the DPSD and meta-*d*’ models for the third memory test when the effects of zolpidem were still near peak, but memory retrieval occurred 270 minutes post-encoding with sleep between these phases. All contrasts found zolpidem at encoding to impair recollection (first placebo vs. first zolpidem: *M* = .26, *SD* = .03, CI = [.20, .33], *p* < .001; first placebo vs. second zolpidem: *M* = .21, *SD* = .05, CI = [.11, .32], *p* < .001; second placebo vs. first zolpidem: *M* = .23, *SD* = .06, CI = [.11, .34], *p* < .001; second placebo vs. second zolpidem: *M* = .18, *SD* = .07, CI = [.04, .31], *p* = .008) and familiarity (first placebo vs. first zolpidem: *M* = .40, *SD* = .17, CI = [.07, .74], *p* = .009; first placebo vs. second zolpidem: *M* = .30, *SD* = .13, CI = [.05, .54], *p* = .010; second placebo vs. first zolpidem: *M* = .43, *SD* = .17, CI = [.12, .77], *p* = .002; second placebo vs. second zolpidem: *M* = .32, *SD* = .17, CI = [.01, .67], *p* = .021). Similar to the second memory test, none of the contrasts found zolpidem at encoding to impair metamemory (first placebo vs. first zolpidem: *M* = .06, *SD* = .36, CI = [-.67, .77], *p* > .250; first placebo vs. second zolpidem: *M* = .12, *SD* = .37, CI = [-.59, .85], *p* > .250; second placebo vs. first zolpidem: *M* = .06, *SD* = .41, CI = [-.77, .85], *p* > .250; second placebo vs. second zolpidem: *M* = .12, *SD* = .32, CI = [-.51, .76], *p* > .250).

### Study 3: The Effects of Triazolam and Ketamine on Encoding

The dataset from the current analysis comes from (Carter, Kleykamp, *et al*., 2013). Because triazolam is a GABA_A_ PAM like alcohol and zolpidem, the previous analyses suggest that recollection, familiarity, and perhaps metamemory should be impaired by triazolam at encoding. However, it is less clear what the effects of ketamine (an NMDA antagonist) at encoding will be. In the original analysis of the current dataset, ketamine at encoding impaired free recall, less consistently impaired recognition, and did not change metamemory, as measured with gamma. In contrast, the NMDA antagonist dextromethorphan was found to impair free recall, recognition, and metamemory using the same methods (Carter, Reissig, *et al*., 2013), and another study with dextromethorphan at encoding found evidence for impairments of recollection and familiarity using the DPSD model (Barrett *et al*., 2018). Therefore, ketamine at encoding would be expected to be impair recollection with possible impairments on familiarity and metamemory.

#### Study Methods

This study is described in-depth elsewhere (Carter, Kleykamp, *et al*., 2013). Briefly, a double-blind, within-subjects design was used in which 20 young to middle-aged adults (median = 24-years-old, range = 19-42, 10 males) came in for 5 sessions during each of which they were orally administered a capsule followed 75 minutes later by an intramuscular infusion. In one session, both administrations were placebo. For two of the sessions, the first administration was triazolam (.2 and .4 mg/70 kg) and the second administration was placebo. For the other two sessions, the first administration was placebo and the second was ketamine (.2 and .4 mg/kg). Ordering of drug conditions was counterbalanced across participants. Data for 5 participants were unable to be acquired leaving an *N* of 15 for the present analysis.

Approximately 80-85 minutes post-capsule ingestion and 5-10 minutes post-infusion, participants completed the encoding phase of a verbal memory test. During encoding, participants were presented with 70 concrete nouns, and they indicated whether each noun was artificial or natural. After the encoding phase, participants completed other cognitive tasks for 120 minutes and then completed the retrieval phase. During retrieval, participants completed a free recall test followed by a recognition test. On the recognition test, participants were presented with the 70 nouns they had seen previously (targets) along with 70 non-presented words (lures). For each word, they used a single response, hybrid old/new six-point confidence scale to indicate whether each stimulus had been previously presented.

#### Results

Figures IVa-c display the distributions of parameter estimates from the DPSD and meta-*d*’ models for the triazolam conditions. Compared to placebo, .2 mg/70 kg of triazolam at encoding impaired recollection (*M* = .10, *SD* = .06, CI = [.00, .23], *p* = .031) but not familiarity (*M* = .04, *SD* = .13, CI = [-.22, .28], *p* > .250) or metamemory (*M* = .05, *SD* = .11, CI = [-.16, .27], *p* > .250). In contrast, the .4 mg/ 70 kg of triazolam impaired recollection (*M* = .43, *SD* = .05, CI = [.33, .54], *p* < .001) and familiarity (*M* = .39, *SD* = .13, CI = [.12, .64], *p* = .003) but not metamemory (*M* = .14, *SD* = .18, CI = [-.22, .50], *p* = .213).

**Figure IV.**
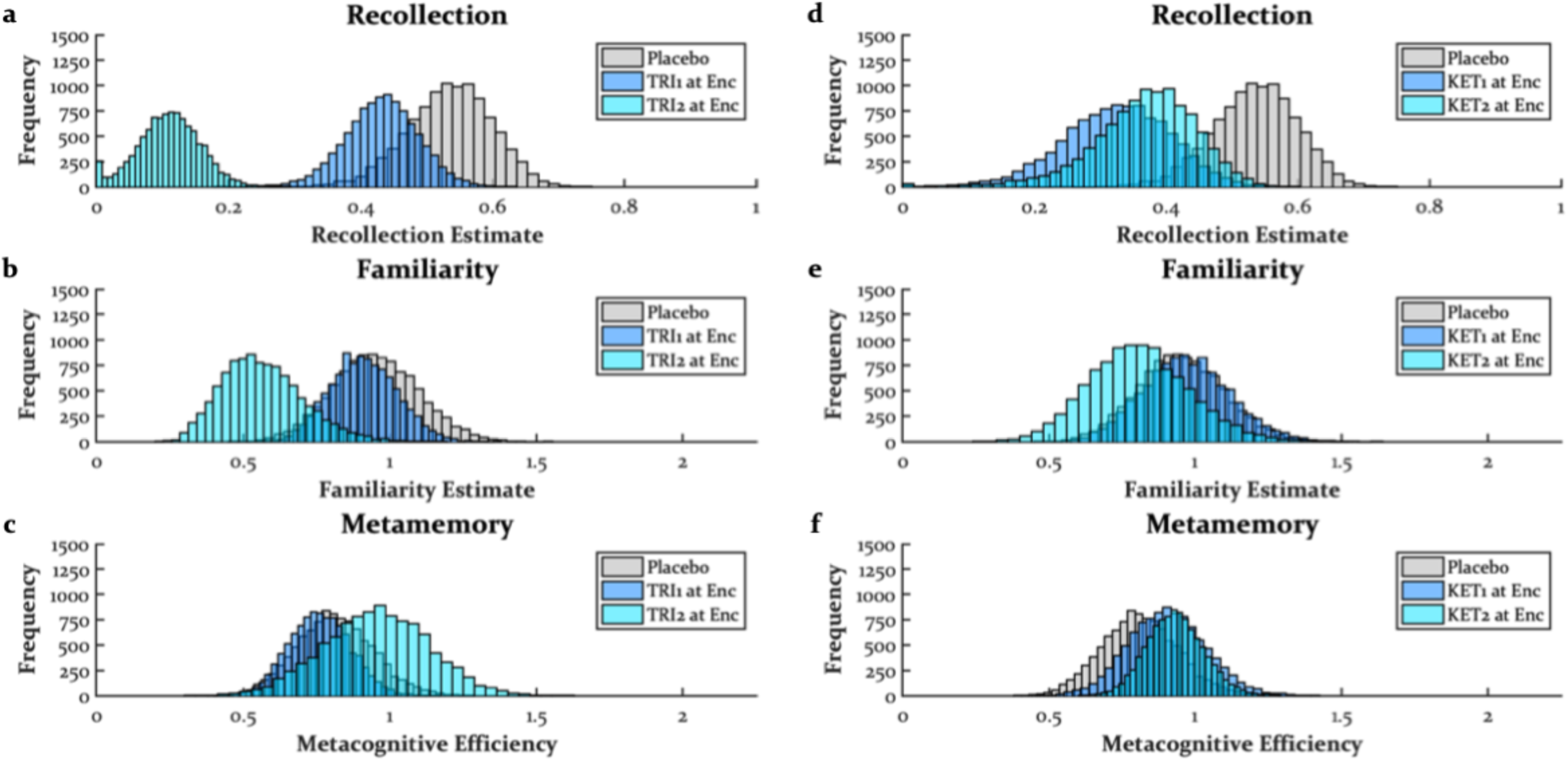
*The Effects of Triazolam and Ketamine at Encoding* Triazolam at encoding impaired recollection and familiarity (at the higher dose) but had no impact on metamemory. Ketamine at encoding impaired recollection and had no impact on familiarity or metamemory, though in other datasets, higher doses of dissociatives (specifically dextromethorphan) at encoding tended to impair familiarity. Also note that ketamine seemed to numerically enhance metamemory with increasing doses, and Study 5 found dextromethorphan at encoding to enhance metamemory. The placebo distributions in the triazolam (a-c) and ketamine (d-f) panels are the same. TRI1 = .2 mg/70 kg triazolam, TRI2 = .4 mg/70 kg triazolam, KET1 = .2 mg/kg ketamine, KET2 = .4 mg/kg ketamine, Enc = encoding.

Figures IVd-f display the distributions of parameter estimates from the DPSD and meta-*d*’ models for the ketamine conditions. Compared to placebo, the lower dose of ketamine at encoding impaired recollection (*M* = .21, *SD* = .08, CI = [.06, .39], *p* = .002) but not familiarity (*M* = .02, *SD* = .18, CI = [-.32, .36], *p* > .250) or metamemory (*M* = .09, *SD* = .12, CI = [-.14, .34], *p* = .228). Similarly, the higher dose of ketamine impaired recollection (*M* = .17, *SD* = .07, CI = [.06, .33], *p* < .001) but not familiarity (*M* = .14, *SD* = .14, CI = [-.14, .42], *p* = .161) or metamemory (*M* = .12, *SD* = .12, CI = [-.13, .34], *p* = .155). Although metamemory was not impacted by ketamine at encoding, it is worth noting that there was a tendency for ketamine to numerically enhance metamemory, especially with the higher dose. In Study 5 but not Study 4, there was a similar trend with the dissociative dextromethorphan at encoding.

### Study 4: The Effects of Triazolam and Dextromethorphan on Encoding

The dataset from the current analysis comes from (Carter, Reissig, *et al*., 2013) and is similar to the previous dataset comparing triazolam at encoding with the dissociative dextromethorphan at encoding. In the original analysis, it was found that both triazolam and dextromethorphan at encoding impaired free recall, recognition, and metamemory measured with gamma. Although it would be expected for triazolam to impair recollection and familiarity, as was the case for most of the previous sedative manipulations, thus far the effects of sedatives on metamemory using metacognitive efficiency have mostly been null. Based on the analysis of the previous dataset, dextromethorphan at encoding would be expected to impair recollection but leave familiarity and metamemory relatively intact.

#### Study Methods

This study is described in-depth elsewhere (Carter, Reissig, *et al*., 2013). Briefly, a double-blind, within-subjects design was used in which 12 young to middle-aged adults (*M* = 27.5-years-old, range = 20-40, 9 males) came in for up to 11 sessions during each of which they were orally administered a capsule containing either placebo, triazolam (.25 and .5 mg/70 kg), or dextromethorphan (100, 200, 300, 400, 500, 600, 700, and 800 mg/70 kg). The order of placebo, triazolam, and dextromethorphan were counterbalanced across participants, but once a given drug was administered, an ascending dose run-up was used across sessions until the maximum tolerable dose was reached for a given participant. 300 mg/70 kg was the maximum dose of dextromethorphan at which all 12 participants were able to complete the memory test with a steep rate of dropout at higher doses. We, therefore, limited analyses of dextromethorphan to the first three doses.

At 120 minutes post-capsule ingestion, participants completed the encoding phase of a verbal memory test. Participants were presented with 36 concrete nouns during each of which they were to decide if it was artificial or natural. After the encoding phase, participants completed several other cognitive tasks, and at 320 minutes post-capsule ingestion, they completed the retrieval phase. First, they completed a free recall test followed by a recognition memory test. Participants were presented in random order with the 36 nouns they had seen previously (targets) along with 36 non-presented words (lures). On each trial, they were to indicate whether they had previously seen the word during the encoding phase using a single response, hybrid old/new six-point confidence scale.

#### Results

Figures Va-c display the distributions of parameter estimates from the DPSD and meta-*d*’ models for the triazolam conditions. Compared to placebo, .25 mg/70 kg of triazolam at encoding impaired recollection (*M* = .28, *SD* = .05, CI = [.17, .38], *p* < .001) and familiarity (*M* = .39, *SD* = .20, CI = [.00, .77], *p* = .026). Unexpectedly, there was a trend for .25 mg/70 kg of triazolam at encoding to enhance metamemory (*M* = .29, *SD* = .19, CI = [-.07, .69], *p* = .056). This pattern was also found with .5 mg/70 kg of triazolam at encoding with larger impairments of recollection (*M* = .53, *SD* = .08, CI = [.38, .68], *p* < .001) and familiarity (*M* = .52, *SD* = .21, CI = [.18, .99], *p* = .001) and a trending enhancement of metamemory (*M* = .79, *SD* = .76, CI = [-.10, 2.59], *p* = .054). However, these enhancements of metamemory should be treated with caution, as the distribution of metamemory estimates for .5 mg/70 kg of triazolam were abnormally wide (and therefore not plotted in Figure Vc).

**Figure V.**
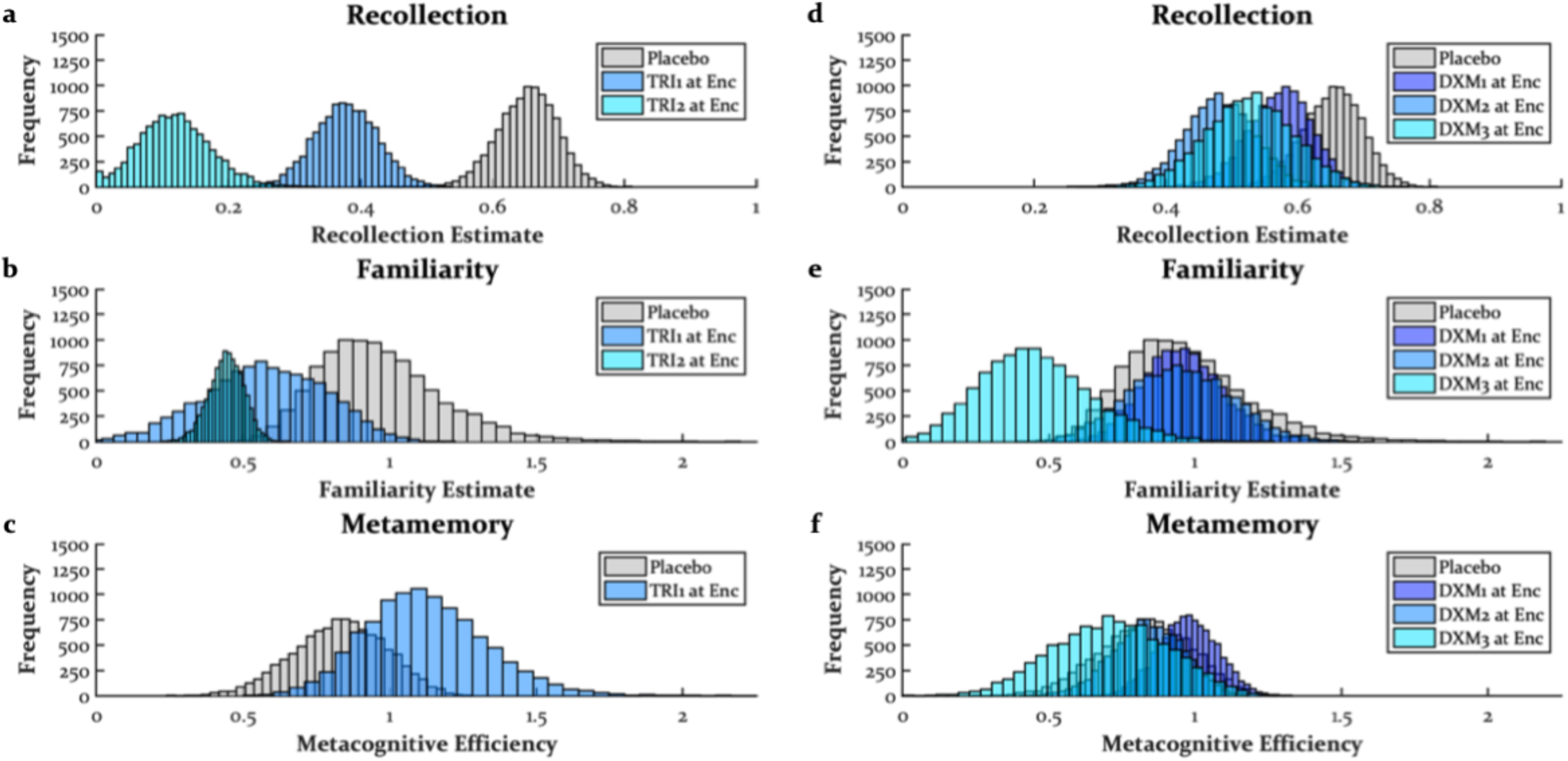
*The Effects of Triazolam and Dextromethorphan at Encoding* Triazolam at encoding impaired recollection and familiarity, and there was a trend for enhancement of metamemory, though this enhancement of metamemory did not replicate for any other sedative manipulation (including similar doses of triazolam in Study 3). Dextromethorphan at encoding impaired recollection and familiarity (at higher doses) but had no impact on metamemory. The placebo distributions in the triazolam (a-c) and dextromethorphan (d-f) panels are the same. The metamemory distribution for .5 mg/70 kg of triazolam was too wide to plot on the scale common to all plots. TRI1 = .25 mg/70 kg triazolam, TRI2 = .5 mg/70 kg triazolam, DXM1 = 100 mg/70 kg dextromethorphan, DXM2 = 200 mg/70 kg dextromethorphan, DXM3 = 300 mg/70 kg dextromethorphan, Enc = encoding.

Figures Vd-f display the distributions of parameter estimates from the DPSD and meta-*d*’ models for the dextromethorphan conditions. Compared to placebo, 100 mg/70 kg of dextromethorphan at encoding impaired recollection (*M* = .08, *SD* = .05, CI = [-.01, .18], *p* = .046) but not familiarity (*M* = .00, *SD* = .21, CI = [-.40, .44] *p* > .250), and it numerically but non-significantly increased metamemory (*M* = .15, *SD* = .17, CI = [-.18, .48], *p* = .194) similar to the previous analysis with ketamine. 200 mg/70 kg of dextromethorphan at encoding also impaired recollection (*M* = .17, *SD* = .08, CI = [.04, .33], *p* = .005) but not familiarity (*M* = .02, *SD* = .20, CI = [-.39, .40], *p* > .250) with no impact on metamemory (*M* = .01, *SD* = .21, CI = [-.40, .41], *p* > .250). Finally, 300 mg/70 kg of dextromethorphan at encoding impaired both recollection (*M* = .13, *SD* = .08, CI = [-.02, .28], *p* = .043) and familiarity (*M* = .51, *SD* = .18, CI = [.17, .88], *p* = .002). In contrast to 100 mg/70 kg of dextromethorphan, 300 mg/70 kg of dextromethorphan at encoding numerically, though non-significantly, decreased metamemory (*M* = .12, *SD* = .14, CI = [-.16, .38], *p* = .199).

### Study 5: The Effects of Dextromethorphan and Psilocybin on Encoding

The dataset from the current analysis comes from (Barrett *et al*., 2018). In the original analysis of the current dataset, a DPSD analysis found that dextromethorphan impaired recollection compared to placebo and impaired familiarity compared to the lowest dose of psilocybin, with this dose of psilocybin not differing appreciably from placebo. However, because the DPSD model was fit to individual participant data (instead of the bootstrapping procedure used here) and the number of trials per participant was low, it is possible that these estimates of recollection and familiarity were unreliable. Based on the previous analyses, it is expected that dextromethorphan will impair recollection and perhaps familiarity, though the effects on metamemory are unclear.

Like dissociatives, psychedelics are also classified as hallucinogens, but their main mechanism of action comes from activation of the 5-HT_2A_ receptor. The previous analysis of this dataset found that all doses of psilocybin at encoding impaired free recall, and the DPSD analysis (fit to individual participant data) found some evidence for psilocybin at encoding to impair recollection at the two higher doses, though these effects were non-significant. In contrast, there was no significant impact of psilocybin at encoding on familiarity, but interestingly, there were numerical enhancements. This is peculiar, as most impairments of recollection in the previous analyses, particularly at higher doses, were accompanied by familiarity impairments. This study did not run a metamemory analysis, but there have been anecdotal claims that classic psychedelics can enhance awareness of internal processes in which case psilocybin at encoding might be expected to enhance metamemory.

#### Study Methods

This study is described in-depth elsewhere (Barrett *et al*., 2018). Briefly, a double-blind, within-subjects design was used in which 20 young to middle-aged adults (*M* = 28.5-years-old, range = 22-43, 9 males) were orally administered a capsule containing placebo, a moderately high dose of dextromethorphan (400 mg/70 kg), a moderate dose of psilocybin (10 mg/70 kg), a moderately high dose of psilocybin (20 mg/70 kg), and a high dose of psilocybin (30 mg/70 kg) prior to encoding stimuli in 4 separate experimental sessions with dose order counterbalanced. One participant did not complete the dose of dextromethorphan, another did not complete the highest dose of psilocybin, and another participant did not complete either of these doses.

During each experimental session, participants consumed a capsule and after 180 minutes, completed the encoding phase of a verbal memory task. Participants were presented with 36 concrete nouns during each of which they were to decide if it was artificial or natural. After the encoding phase, participants completed several other cognitive tasks followed by a free recall test for the verbal stimuli from the encoding phase. At 390 minutes post-dosing, participants completed a recognition memory test. Participants were presented with the 36 nouns they had seen previously (targets) along with 36 non-presented words (lures). On each trial, they were to indicate whether they had previously seen the word during the encoding phase using a single response, hybrid old/new six-point confidence scale.

#### Results

Figures VIa-c display the distributions of parameter estimates from the DPSD and meta-*d*’ models for the dextromethorphan condition. Dextromethorphan at encoding impaired recollection (*M* = .18, *SD* = .07, CI = [.06, .32], *p* = .002). Although the effect of dextromethorphan at encoding on familiarity was not significant (*M* = .23, *SD* = .20, CI = [-.19, .59], *p* = .125), the wide distribution of estimates in the dextromethorphan condition suggests that at least some participants may have experienced familiarity impairments similar to the previous analysis with dextromethorphan. Interestingly, it appeared that dextromethorphan at encoding enhanced metamemory (*M* = .40, *SD* = .24, CI = [-.04, .91], *p* = .039) similar to the numerical increases in the previous analyses with ketamine and 100 mg/70 kg of dextromethorphan at encoding.

**Figure VI.**
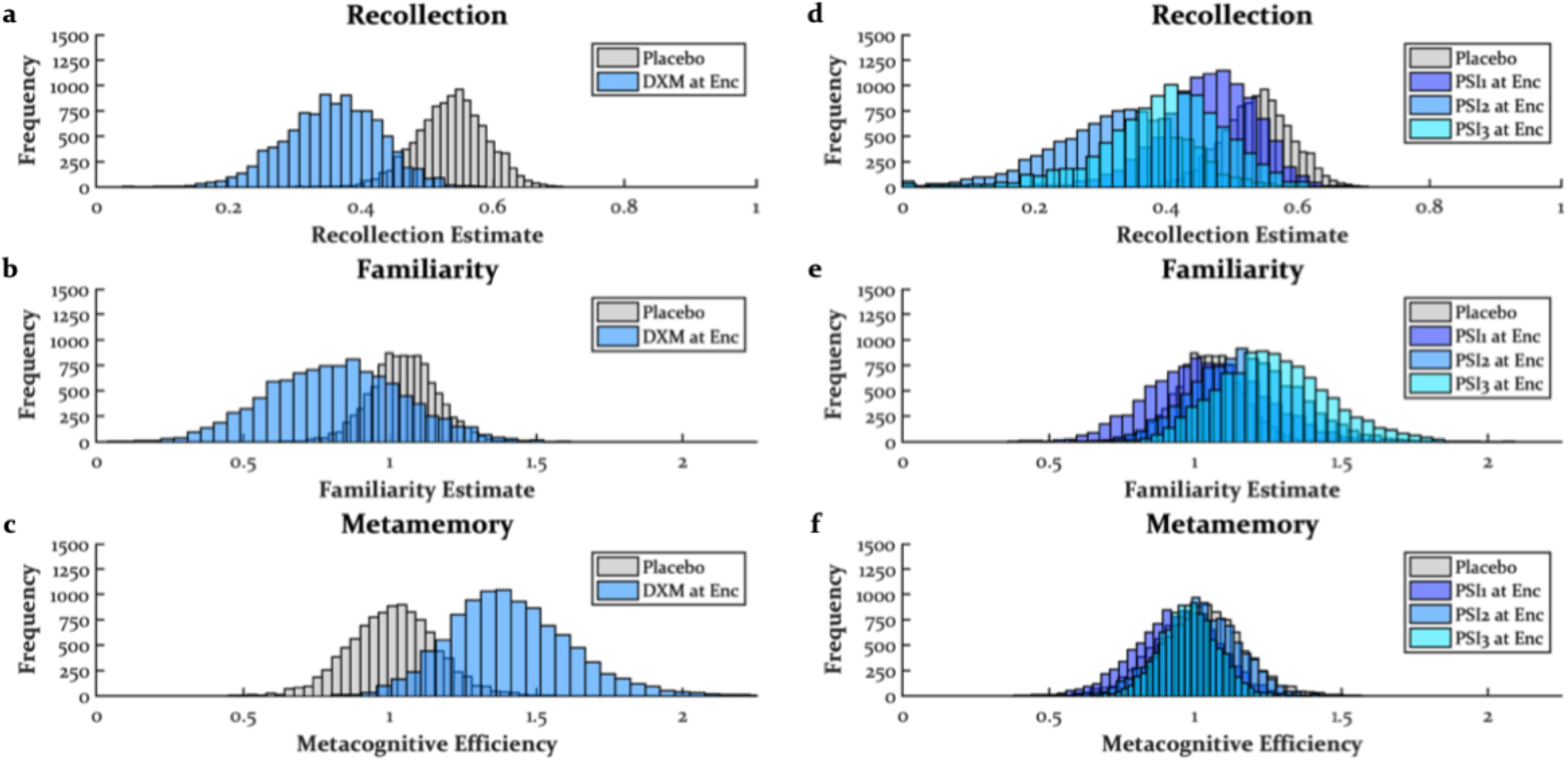
*The Effects of Dextromethorphan and Psilocybin at Encoding* Dextromethorphan at encoding impaired recollection, and although the impairment of familiarity was not significant, a lower dose of dextromethorphan in Study 4 did find a significant impairment. Dextromethorphan at encoding enhanced metamemory, and although this effect did not replicate in Study 4, such numerical enhancements were found with ketamine at encoding in Study 3. Psilocybin at encoding impaired recollection and spared or even enhanced familiarity at the higher doses, a trend observed in Study 6 with MDMA. Psilocybin had no impact on metamemory. The placebo distributions in dextromethorphan (a-c) and psilocybin (d-f) panels are the same. DXM = 400 mg/70 kg dextromethorphan, PSI1 = 10 mg/70 kg psilocybin, PSI2 = 20 mg/70 kg, PSI3 = 30 mg/70 kg, Enc = encoding.

Figures VId-f displays the distributions of parameter estimates from the DPSD and meta-*d*’ models for the psilocybin conditions. The 10 mg/70 kg (*M* = .07, *SD* = .05, CI = [-.01, .19], *p* = .049), 20 mg/70 kg (*M* = .21, *SD* = .10, CI = [.05, .44], *p* = .004), and 30 mg/70 kg (*M* = .14, *SD* = .09, CI = [-.02, .36], *p* = .046) doses of psilocybin at encoding impaired recollection with numerically larger effects from the 20 and 30 mg/70 kg doses. Although there was no impact of the 10 mg/70 kg of psilocybin at encoding on familiarity (*M* = .01, *SD* = .16, CI = [-.33, .30], *p* > .250), the 20 mg/70 kg (*M* = .12, *SD* = .15, CI = [-.15, .44], *p* = .212) and 30 mg/70 kg (*M* = .22, *SD* = .18, CI = [-.12, .59], *p* = .106) doses of psilocybin at encoding numerically enhanced familiarity with a marginal trend of the highest dose. Contrary to the idea that psychedelics could enhance awareness of mnemonic processes, there was no evidence that psychedelics impacted metamemory (10 mg/70 kg: *M* = .07, *SD* = .11, CI = [-.13, .29], *p* > .250; 20 mg/70 kg: *M* = .01, *SD* = .11, CI = [-.22, .23], *p* > .250; 30 mg/70 kg: *M* = .03, *SD* = .13, CI = [-.23, .27], *p* > .250).

### Study 6: The Effects of MDMA on Encoding and Retrieval

The dataset from the current analysis comes from (Doss, Weafer, Gallo, *et al*., 2018a). In the original analysis of the current dataset, a DPSD analysis (using a bootstrapping procedure) found that MDMA at encoding impaired recollection but only for negative and positive stimuli with similar trends on a remember/know test. Interestingly, there were trends for MDMA at encoding to enhance DPSD familiarity estimates, though again only for emotional stimuli. This study did not examine the effects of MDMA on metamemory, but like psilocybin, MDMA activates the 5-HT_2A_ receptor (despite not always being considered a psychedelic). Thus, the previous analysis would suggest that MDMA at encoding might not impact metamemory. Given MDMA’s facilitation of monoaminergic transmission, however, it is also possible that MDMA at encoding could have similar effects as stimulants such as dextroamphetamine and dextromethamphetamine. Based on the analyses of stimulant data below, it will become clear that MDMA’s mnemonic effects more closely resemble a psychedelic.

In the original analysis of these data, MDMA at retrieval did not significantly impact recollection or familiarity (Doss, Weafer, Gallo, *et al*., 2018a). There were numerical decreases from MDMA at retrieval on recollection, but similar to the effects of dextroamphetamine and THC at retrieval (Ballard *et al*., 2014; Doss, Weafer, Gallo, *et al*., 2018b), there was some evidence for MDMA at retrieval to increase high confidence false alarms. Such false memory effects can bias recollection estimates toward zero but might be qualitatively different from actual impairments of recollection. Although no study has tested the effects of a psychedelic at retrieval on metamemory, it is worth noting that drug effects during retrieval should directly modulate metacognitive operations in contrast to drug manipulations of encoding or consolidation before metamemory decisions are ever made. Therefore, MDMA at retrieval might enhance metamemory considering anecdotal reports claiming insights into one’s mind.

#### Study Methods

This study is described in-depth elsewhere (Doss, Weafer, Gallo, *et al*., 2018a). Briefly, a double-blind, between-subjects design in which 60 young adults (*M* = 23.7-years-old, range = 18-34) were randomized to one of three groups: placebo (*N* = 20, 10 males), encoding (*N* = 20, 10 males), or retrieval (*N* = 20, 10 males). The placebo group received placebo prior to encoding and placebo prior to retrieval. The encoding group received MDMA prior to encoding and placebo prior to retrieval. The retrieval group received placebo prior to encoding and MDMA prior to retrieval. Doses of MDMA (1 mg/kg) were administered orally via capsules.

Participants came in for 2 sessions separated by 48 hours. During the first session, participants consumed their first capsule and 90 minutes later completed the encoding phase of a 180-trial cued recollection task. On each trial, participants were presented with a negative, neutral, or positive label during which they were to indicate on a five-point scale how much they would like to see a picture of the label. On trials in which a picture was presented, they were to rate the picture’s valence (five-by-five grid with negative and positive valence ratings on orthogonal axes) and arousal (five-point scale). After the encoding phase, participants remained in the lab until three and a half hours post-capsule administration during the first two hours of which they were only permitted to listen to music without lyrics to minimize retroactive interference. At the beginning of the second session, participants consumed their second capsule, and 90 minutes later, they completed the retrieval phase of the cued recollection task. Participants were presented with all 180 labels from the encoding phase and were to decide if they had seen a picture of each label (yes/no) followed by a confidence rating (five-point scale). Therefore, labels for which a picture had been presented were targets, and labels presented without a corresponding picture were lures.

#### Results

Figures VIIa-c display the distributions of parameter estimates from the DPSD and meta-*d*’ models. MDMA at encoding impaired recollection (*M* = .09, *SD* = .05, CI = [-.02, .19], *p* = .057), though this effect was marginal because of collapsing across emotional conditions (effects were only apparent for negative and positive stimuli; SM). Like psilocybin, MDMA at encoding numerically boosted familiarity (*M* = .12, *SD* = .10, CI = [-.09, .32], *p* = .131) with stronger effects on emotional stimuli (see SM). MDMA at encoding did not significantly impact metamemory (*M* = .10, *SD* = .09, CI = [-.07, .28], *p* = .143), though there was a trend for an enhancement of positive stimuli (SM). MDMA at retrieval did not impact recollection (*M* = .04, *SD* = .06, CI = [-.07, .16], *p* = .224), familiarity (*M* = .08, *SD* = .10, CI = [-.11, .29], *p* = .204), or metamemory (*M* = .04, *SD* = .09, CI = [-.13, .22], *p* > .250). Thus, once again, there was no evidence for a psychedelic enhancement of the awareness of mnemonic processes.

**Figure VII.**
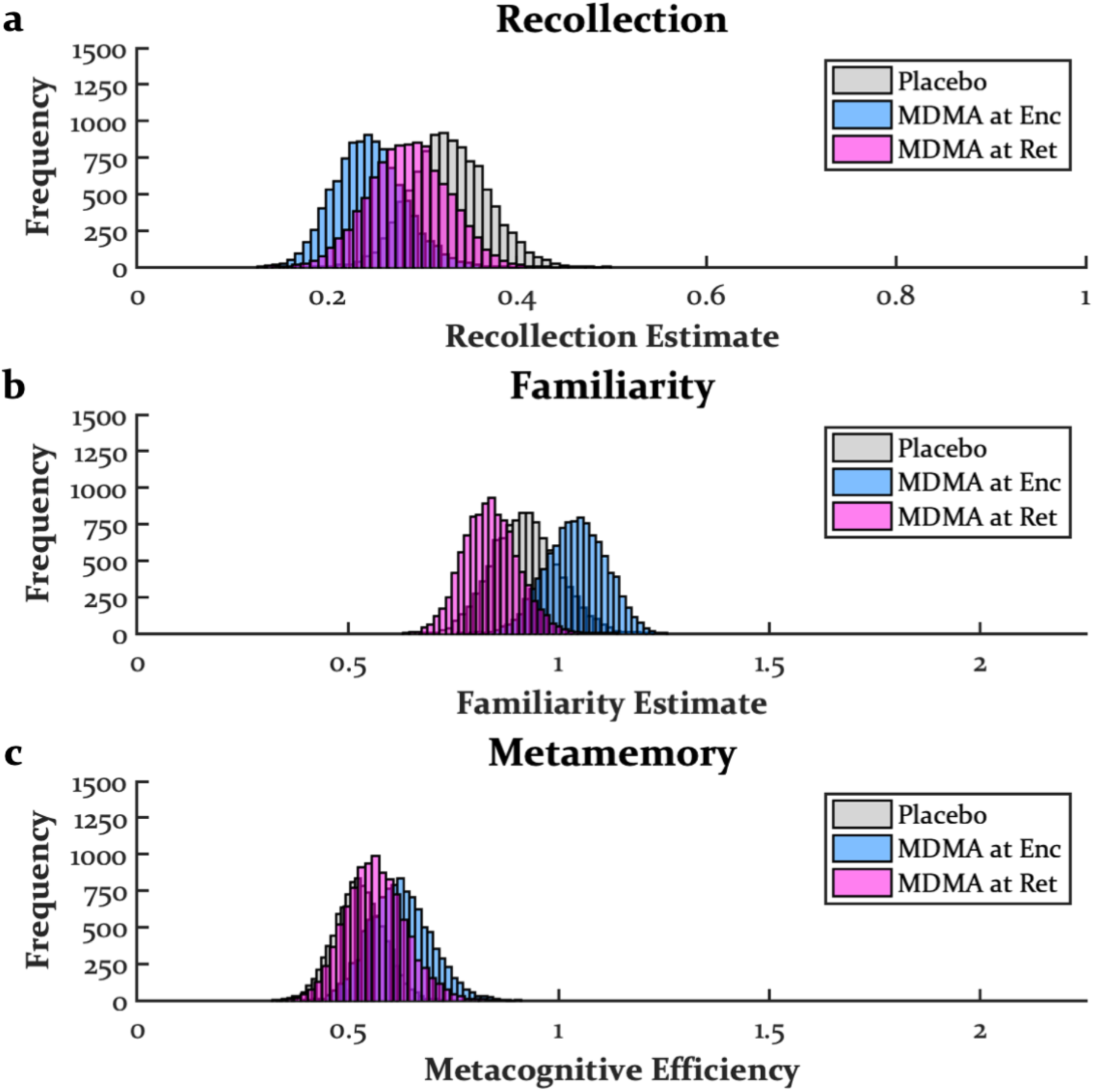
*The Effects* ±*3,4-Methylenedioxymethamphetamine at Encoding and Retrieval* MDMA at encoding impaired recollection and enhanced familiarity, particularly for emotional stimuli (see Supplemental Material) but had no impact on metamemory. These effects are consistent with its 5-HT_2A_ agonism effects (i.e., psilocybin’s effects in Study 5) but not its stimulant effects (i.e., dextroamphetamine and dextromethamphetamine’s effects in Studies 7 and 8). MDMA at retrieval had no impact on recollection, familiarity, or metamemory, again not exhibiting the stimulant effects observed in Study 7. Doses used here were 1 mg/kg. MDMA = ±3,4-methylenedioxymethamphetamine. Enc = encoding, Ret = retrieval.

### Study 7: The Effects of Dextroamphetamine on Encoding, Retrieval, and Both Encoding and Retrieval

The dataset from the current analysis comes from (Weafer *et al*., 2014). Like MDMA, dextroamphetamine increases synaptic monoamines, but as a typical stimulant, its effects are most prominent on dopamine and norepinephrine, and it does not activate 5-HT_2A_ receptors. In the original analysis of the current dataset, dextroamphetamine at encoding, retrieval, or encoding and retrieval did not impact any measure of memory on a cued recollection task. However, others have found dextroamphetamine and methylphenidate at encoding to improve memory on recollection-based tasks (Linssen *et al*., 2012; Soetens *et al*., 1995; Zeeuws *et al*., 2010). One possibility is that by dissociating recollection and familiarity with the DPSD model, we may also observe enhancements from dextroamphetamine at encoding. No study has examined the effects of dextroamphetamine on familiarity or metamemory, though it might be expected for dextroamphetamine at encoding to enhance metamemory, as stimulants can reverse the metamemory (measured with gamma) impairment induced by sedatives (Mintzer and Griffiths, 2003b, 2007).

Like MDMA at retrieval (Doss, Weafer, Gallo, *et al*., 2018a), as well as THC at retrieval (Doss, Weafer, Gallo, *et al*., 2018b), dextroamphetamine at retrieval has been found to increase recall intrusions and false recognition (Ballard *et al*., 2014). Therefore, recollection estimates may be attenuated with dextroamphetamine at retrieval because of the known zero-inflation when false alarms, particularly with high confidence, are elevated. However, the original study did not find evidence for an increase in false alarms.

Some studies have found administration of dextroamphetamine at both encoding and retrieval with a ≥24 delay between the two phases resulted in recollection-based memory enhancements that were greater than encoding enhancements alone (Bustamante *et al*., 1970; Shea, 1982). These state-dependent effects typically require an initial encoding enhancement from stimulants, but other studies have found no evidence for enhancements when a stimulant is given during both encoding and retrieval on recollection-based tasks (Aman and Sprague, 1974; Steinhausen and Kreuzer, 1981, p. 1; Weingartner *et al*., 1982; Becker-Mattes *et al*., 1985). It is worth noting that if both encoding and retrieval enhancements are found in isolation, then a larger enhancement when dextroamphetamine is administered at both encoding and retrieval may simply be additive effects from the individual encoding and retrieval enhancements rather than state-dependency.

#### Study Methods

This study is described in-depth elsewhere (Weafer *et al*., 2014). Briefly, a double-blind, between-subjects design was used in which 59 young adults (*M* = 23.9-years-old, range = 18-35) were randomized to one of four demographically similar groups: placebo (*N* = 20, 10 males), encoding (*N* = 20, 10 males), retrieval (*N* = 20, 10 males), or both encoding and retrieval (*N* = 20, 10 males). Participants came in for an encoding session followed 48 hours later by a retrieval session. The placebo group received placebo at encoding and placebo at retrieval. The encoding group received dextroamphetamine at encoding and placebo at retrieval. The retrieval group received placebo at encoding and dextroamphetamine at retrieval. The both encoding and retrieval group (or simply, “both” group) received dextroamphetamine at encoding and dextroamphetamine at retrieval. Doses of dextroamphetamine (20 mg) were administered orally via capsules.

During the first session, participants consumed the first capsule and 90 minutes later completed the encoding phase of a 144-trial cued recollection task. On each trial, participants were presented with a negative, neutral, or positive label (48 of each), half of which were followed by a picture of that label. During the presentation of each label, participants were to indicate on a five-point scale how likely they would be to remember a picture of the label. On trials in which a picture was presented, they were to rate the picture’s valence (five-by-five grid with negative and positive valence ratings on orthogonal axes) and arousal (five-point scale). During the second session, participants consumed the second capsule and 90 minutes later completed the retrieval phase of the cued recollection task. On each trial, participants were presented with a label from the encoding phase and asked whether they had seen a picture of that label (yes/no) followed by a confidence rating on a five-point scale. Therefore, labels for which a picture had been presented were targets and labels presented without a corresponding picture were lures.

#### Results

Figures VIIIa-c display the distributions of parameter estimates from the DPSD and meta-*d*’ models. Dextroamphetamine at encoding did not impact recollection (*M* = .01, *SD* = .06, CI = [-.11, .13], *p* > .250), familiarity (*M* = .06, *SD* = .15, CI = [-.22, .37], *p* > .250), or metamemory (*M* = .11, SD = .11, *CI* = [-.10, .32], *p* = .142). Dextroamphetamine at retrieval did not impact recollection (*M* = .03, *SD* = .06, CI = [-.08, .14], *p* > .250), though it did enhance familiarity (*M* = .34, *SD* = .15, CI = [.05, .63], *p* = .010) and metamemory (*M* = .20, *SD* = .10, CI = [.00, .40], *p* = .023). Finally, dextroamphetamine at both encoding and retrieval did not impact recollection (*M* = .04, *SD* = .06, CI = [-.07, .16], *p* = .218), but there was a trend for a familiarity enhancement (*M* = .19, *SD* = .15, CI = [-.10, .50], *p* = .102) and a robust metamemory enhancement (*M* = .28, *SD* = .11, CI = [.07, .50], *p* = .004). Because the familiarity enhancement was most apparent in the retrieval group and nonexistent in the encoding group, the trending familiarity enhancement in the “both” group is likely to be driven by dextroamphetamine at retrieval alone. Although the metamemory enhancement was numerically greater in the “both” group than the retrieval group, suggestive of a state-dependent effect, the numerical increase of metamemory in the encoding group makes it difficult to differentiate state-dependency from potential additive enhancements of encoding and retrieval effects. All metamemory enhancements, including numerical enhancements in the encoding group, were greater for emotional compared to neutral material (see SM).

**Figure VIII.**
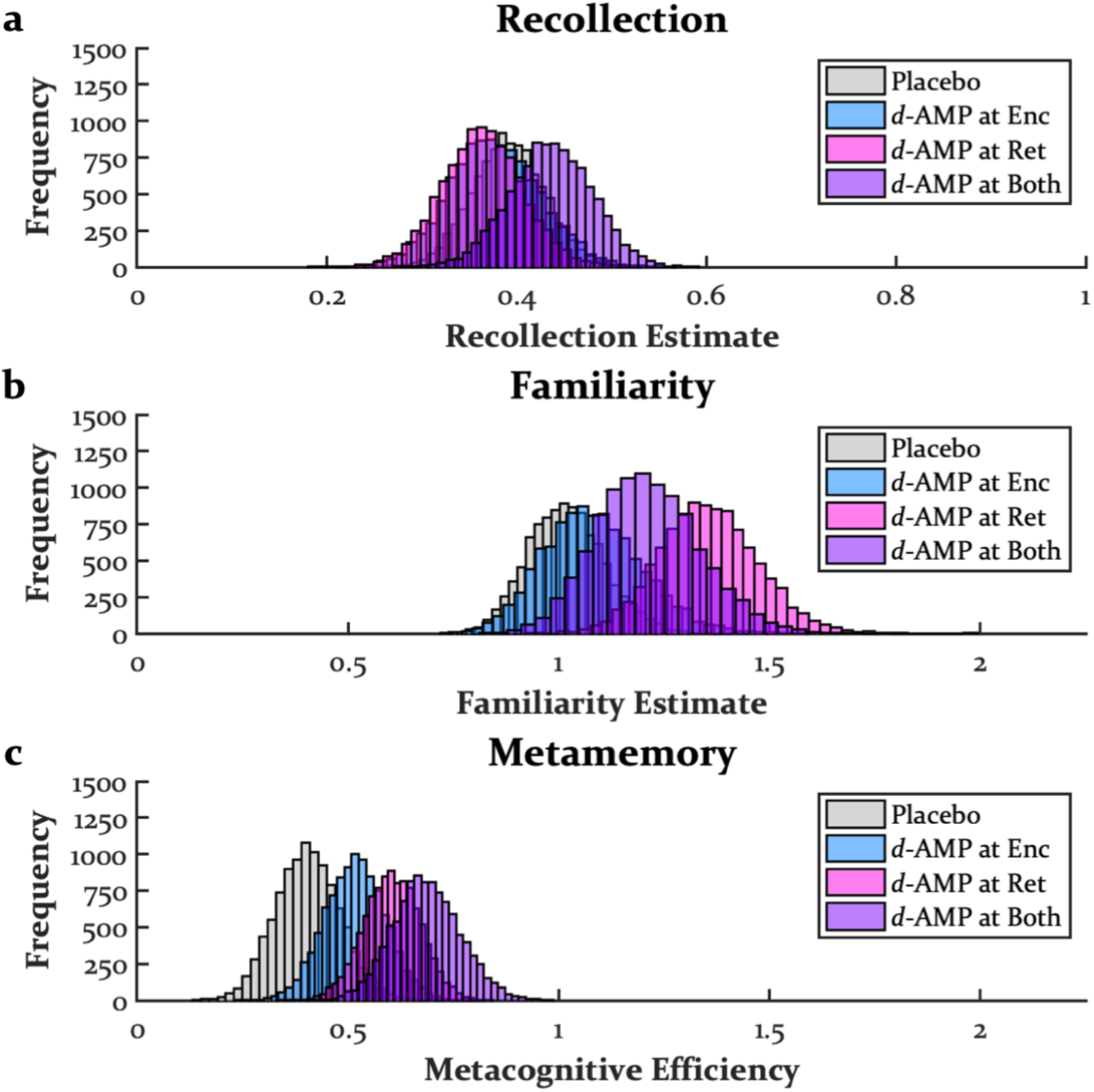
*The Effects Dextroamphetamine at Encoding, Retrieval, and Both Encoding and Retrieval* Dextroamphetamine at encoding had no impact on recollection, familiarity, or metamemory. Dextroamphetamine at retrieval did not impact recollection, but it enhanced familiarity and metamemory. Dextroamphetamine at both encoding and retrieval did not impact recollection, but there was a numerical enhancement of familiarity, consistent with dextroamphetamine at retrieval, and an enhancement of metamemory that was larger than dextroamphetamine either at encoding or retrieval alone. Doses used here were 20 mg. *d*-AMP = dextroamphetamine, Enc = encoding, Ret = retrieval, Both = both encoding and retrieval.

### Study 8: The Effects of Dextromethamphetamine on Encoding and Consolidation

The dataset from the current analysis comes from (Ballard *et al*., 2015). Dextromethamphetamine is pharmacologically similar to dextroamphetamine, but it is more rapidly absorbed and, therefore, better able to target post-encoding memory stabilization (i.e., consolidation). In the original analysis of the current dataset, dextromethamphetamine at encoding or consolidation was not found to impact memory on a picture recognition test. However, the prolonged duration of dextromethamphetamine effects caused poor sleep in several participants. A post hoc analysis found that those participants who did not report sleep difficulties exhibited memory enhancements for emotional pictures from dextromethamphetamine at encoding. Based on the post hoc nature of these findings and the absence of an effect of dextroamphetamine at encoding in the previous analysis, it might not be expected for dextromethamphetamine at encoding to impact recollection and familiarity. However, the numerical enhancement of metamemory from dextroamphetamine at encoding and the significant enhancement of dextroamphetamine at both encoding and retrieval suggest that dextromethamphetamine at encoding could enhance metamemory. It is unclear whether dextromethamphetamine at consolidation will impact recollection, familiarity, or metamemory.

#### Study Methods

This study is described in-depth elsewhere (Ballard *et al*., 2015). Briefly, a double-blind, mixed within- and between-subjects design was used in which 60 young adults (*M* = 23.5-years-old, range = 18-34) were randomized to one of 2 demographically similar groups: encoding (*N* = 29, 14 males) or consolidation (*N* = 31, 15 males). Each group received two doses of dextromethamphetamine (10 and 20 mg) and had its own placebo condition, essentially making this study two separate within-subjects experiments. Participants came in for 3 cycles of an encoding session followed 48 hours later by a retrieval session. The encoding group received each drug manipulation prior to encoding stimuli, and the consolidation group received each drug manipulation immediately post-encoding. Because memory consolidation is a time-dependent process in which recently encoded memories stabilize as more time passes (Wixted, 2004), tablets of dextromethamphetamine were crushed and suspended in a flavored syrup before oral administration to achieve a rapid onset. This was done for both the encoding and consolidation manipulations.

During the first session of each encoding/retrieval cycle, participants completed the encoding phase of a 60-trial picture recognition task. On each trial, participants were presented with a negative, neutral, or positive label (20 of each) during which they were to rate the picture’s valence (5-by-5 grid with negative and positive valence ratings on orthogonal axes) and arousal (9-point scale). During the second session, participants completed the retrieval phase of the picture recognition task. Participants were presented in random order with the 60 pictures they had seen previously (targets) along with 60 non-presented pictures (lures). On each trial, they were to indicate whether they had previously seen the picture during the encoding phase (yes/no) followed by a confidence rating on a 9-point scale and valence and arousal ratings, resulting in an 18-point scale. Because this was a large scale that was not fully utilized by all participants (a problem with larger scales; Mickes *et al*., 2007), we collapsed across 3 confidence bins, resulting in 6 confidence bins. After each confidence rating, participants rated the valence and arousal of each picture using the same scales that were used at encoding.

#### Results

Figures IXa-c display the distributions of parameter estimates from the DPSD and meta-*d*’ models for the dextromethamphetamine at encoding condition. Compared to the other datasets, familiarity estimates were larger in this study. Neither 10 mg (*M* = .10, *SD* = .11, CI = [-.07, .39], *p* = .159) nor 20 mg (*M* = .03, *SD* = .06, CI = [-.08, .16], *p* > .250) of dextromethamphetamine at encoding had an impact on recollection. The 10 mg dose did appear to enhance familiarity (*M* = .30, *SD* = .19, CI = [-.04, .71], *p* = .045), but without any trend for an enhancement of the 20 mg dose (*M* = .03, *SD* = .17, CI = [-.27, .38], *p* > .250), it is possible that this finding was spurious. In contrast, there were trends for both 10 mg (*M* = .08, *SD* = .05, CI = [-.03, .18], *p* = .072) and 20 mg (*M* = .11, *SD* = .08, CI = [-.05, .27], *p* = .096) of dextromethamphetamine to enhance metamemory, consistent with trends found in the previous analyses when dextroamphetamine was onboard during memory encoding.

**Figure IX.**
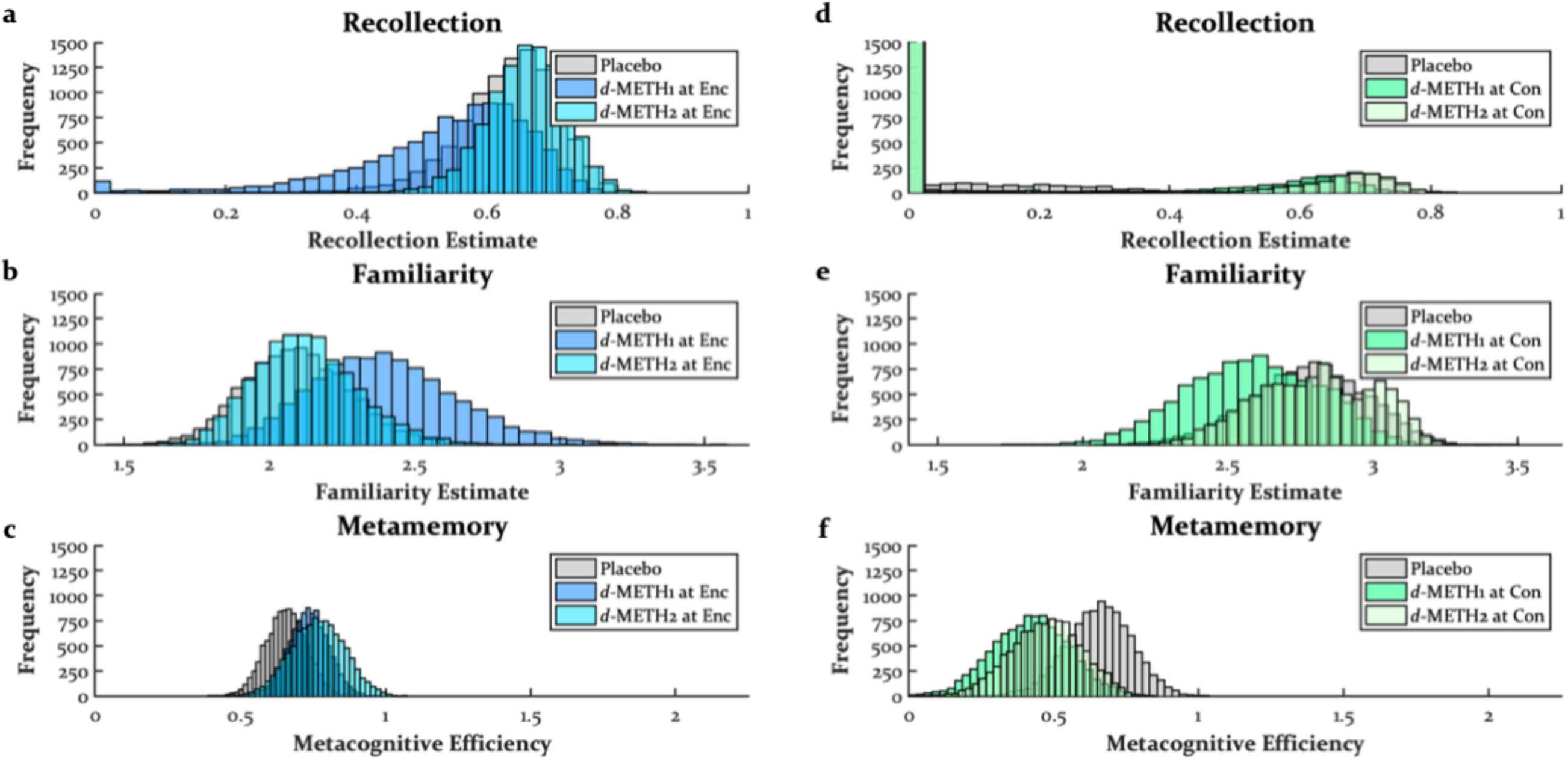
*The Effects Dextromethamphetamine at Encoding and Consolidation* Dextromethamphetamine at encoding had no impact on recollection, and the lower dose enhanced familiarity, though no such trend was found with the higher dose or with dextroamphetamine in Study 7. Both doses of dextromethamphetamine at encoding exhibited trends for enhancing metamemory consistent with Study 7. The consolidation group had unusually low recollection estimates (including in the placebo condition) and are therefore uninterpretable. Dextromethamphetamine at consolidation had no impact on recollection or familiarity, but both doses impaired metamemory. *d*-METH1 = 10 mg dextromethamphetamine, *d*-METH2 = 20 mg dextromethamphetamine, Enc = encoding, Con = consolidation. Note that recollection estimates were oddly biased toward zero, especially in the consolidation condition, and are therefore not reliable.

Figures IXd-f display the distributions of parameter estimates from the DPSD and meta-*d*’ models for the dextromethamphetamine at consolidation group. The familiarity estimates were larger in these data than the other datasets like the encoding group, but the distributions of recollection estimates were zero-inflated for all conditions, suggesting that these estimates are unreliable. Neither 10 nor 20 mg of dextromethamphetamine at consolidation had any impact on recollection (10 mg: *M* = .03, *SD* = .07, CI = [-.02, .27], *p* > .250; 20 mg: *M* = .02, *SD* = .08, CI = [-.08, .27], *p* > .250) or familiarity (10 mg: *M* = .19, *SD* = .17, CI = [-.26, .42], *p* = .115; 20 mg: *M* = .02, *SD* = .15, CI = [-.19, .43], *p* > .250). Interestingly, both 10 mg (*M* = .23, *SD* = .07, CI = [.09, .37], *p* = .001) and 20 mg (*M* = .18, *SD* = .08, CI = [.02, .34], *p* = .017) of dextromethamphetamine at consolidation impaired metamemory. This cannot be explained by the bias toward high confidence false alarms, as the placebo condition had metamemory estimates similar to the placebo condition in the encoding group. Moreover, such biases should be accounted for in signal detection analyses. This metamemory impairment is in sharp contrast to the previous findings that found stimulants at encoding, retrieval, or both encoding and retrieval to enhance metamemory.

### Study 9: The Effects of THC on Encoding

The dataset from the current analysis comes from (Doss, Weafer, *et al*., 2020). In the original analysis of these data, THC was found to robustly impair the encoding of the perceptual details of objects while sparing the influence of context on memory. When THC was administered at encoding, it made participants worse at discriminating between old objects and perceptually similar objects. Because this is a hippocampally-dependent task (Kirwan and Stark, 2007; Lacy *et al*., 2010), it might be expected for THC at encoding to impair recollection. During the retrieval phase, there was a context reinstatement manipulation such that these objects could be presented on the same scene they were paired with during the encoding phase or a different scene. Reinstating the encoding context at retrieval boosted true memory (as well as false memory for similar objects), but THC at encoding was not found to modulate context reinstatement (THC did, however, modulate the false memory effect in a post hoc analysis). Context reinstatement might drive both recollection (Diana *et al*., 2007) and familiarity (Doss, Picart, and Gallo, 2018), effects found here in this dataset (SM). Therefore, THC at encoding may have less of an impact on recollection compared to other drugs and perhaps even spare familiarity. No study has examined the effects of THC during encoding on metamemory, but one study found THC to impair metacognition during working memory decisions (Adam *et al*., 2020) using a measure of metacognition susceptible to the same artefacts as gamma.

#### Study Methods

This study is described in-depth elsewhere (Doss, Weafer, *et al*., 2020). Briefly, a double-blind, within-subjects design was used in which 24 young adults (*M* = 23.0-years-old, range = 18-29, 12 males) received a capsule containing placebo and THC (15 mg) approximately 155 minutes prior to encoding stimuli in 2 separate experimental arms. During the first session of each arm, participants completed the encoding phase of the Mnemonic Similarity Task-Doss version (MS-Doss; Doss, Picart, and Gallo, 2018). During the encoding phase of the MS-Doss, 80 object pictures superimposed over scenes were presented (e.g., black cat on the beach), and participants were to decide if the object belonged in the scene (yes/no).

Forty-eight hours later, participants completed the retrieval phase of the MS-Doss. Participants were again presented with object-scene pairings, but the object could be the original object (e.g., black cat), a similar object (e.g., tuxedo cat), or a completely new object for which an exemplar had not been presented (e.g., rollerblade). Moreover, the context could be reinstated for old and similar objects by presenting them on their scene from encoding (e.g., beach), or the context could be switched (e.g., forest). Participants were to decide whether the object (but not the scene) was “old,” “similar,” or “new” and then rate their confidence on a five-point scale. For this analysis, we collapsed the context conditions (see SM for analysis of these conditions), ignored the data from similar objects, and collapsed “similar” and “new” responses by summing them, resulting in a 10-point scale.

#### Results

Figures Xa-c display the distributions of parameter estimates from the DPSD and meta-*d*’ models. THC at encoding impaired recollection (*M* = .10, *SD* = .04, CI = [.03, .17], *p* = .003) and also marginally impaired familiarity (*M* = .15, *SD* = .10, CI = [-.03, .35], *p* = .059). Surprisingly, THC at encoding enhanced metamemory (*M* = .13, *SD* = .06, CI = [.02, .27], *p* = .011).

**Figure X.**
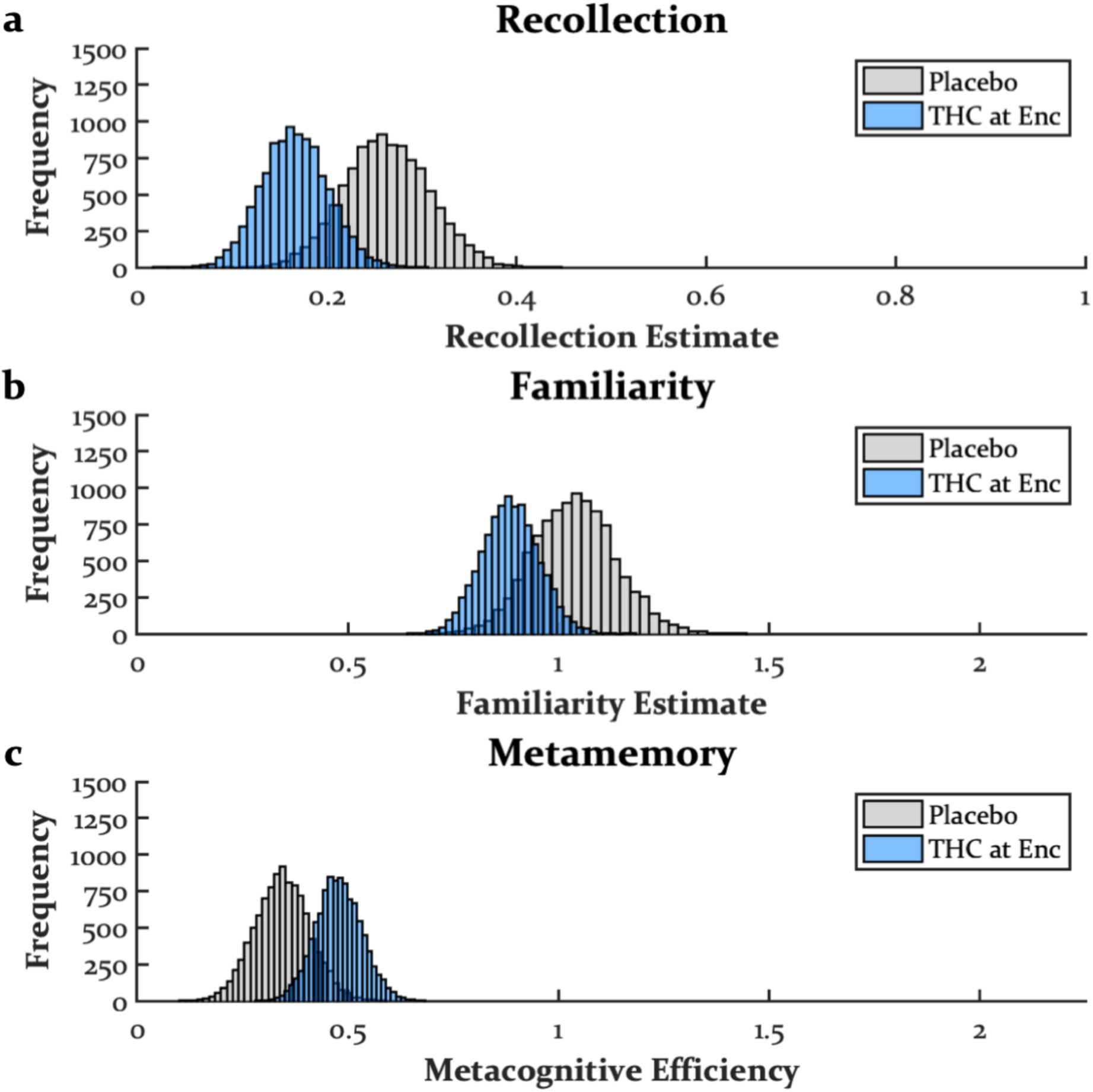
*The Effects Δ^9^-Tetrahydrocannabinol at Encoding* THC at encoding impaired recollection, trended on impairing familiarity, and enhanced metamemory. Dose used here was 15 mg. THC = Δ^9^-tetrahydrocannabinol, Enc = encoding.

### Study 10: The Effects of THC on Retrieval

The dataset from the current analysis comes from (Doss, Weafer, Gallo, *et al*., 2018b). In the original analysis, it was found that THC at retrieval robustly increased false alarms, including high confidence false alarms, and hit rates were unaffected. These findings were interpreted as an increase in false memories, a cognitive phenomenon outside of standard dual process models (but see Lampinen *et al*., 2006). Such an increase in high confidence false alarms would be expected to decrease estimates of recollection, as this would move the first point of the DPSD ROC curve away from the *y*-axis. However, this may not actually be an impairment in recollection, as others have found no impairment of THC at retrieval (with sober encoding) on verbal free recall (Ranganathan *et al*., 2017). No study has looked at THC’s effects during retrieval on familiarity. With THC at retrieval increasing false alarms, it is possible for metamemory to be attenuated. However, a boost in high confidence false memories may simply appear as a criterion shift in signal detection models in which case meta-*d*’ may not capture such an effect, as discrimination measures are bias-free.

#### Study Methods

This study is described in-depth elsewhere (Doss, Weafer, Gallo, *et al*., 2018b) and was the same participants as those in Study 9 (with one participant excluded for not following directions). Briefly, a double-blind, within-subjects design was used in which 23 young adults (*M* = 22.7-years-old, range = 18-29, 11 males) received a capsule containing placebo and THC (15 mg) prior to the retrieval phase (sober encoding) in 2 separate experimental arms. During the first session of each arm, participants completed the encoding phase of a 120-trial cued recollection task. On each trial, participants were presented with a negative, neutral, or positive label (40 of each valence), half of which were followed by a picture of that label. During the presentation of each label, participants were to rate its arousal on a five-point scale. On trials in which a picture was presented, they were again to make an arousal rating (five-point scale) for the picture.

Forty-eight hours later, participants came in for a second session. Participants consumed a capsule and 120 minutes later completed the retrieval phase of the cued recollection task. On each trial, participants were presented with a label from the encoding phase and asked whether they had seen a picture of that label (yes/no) followed by a confidence rating on a five-point scale. Therefore, labels for which a picture had been presented were targets and labels presented without a corresponding picture were lures.

#### Results

Figures XIa-c display the distributions of parameter estimates from the DPSD and meta-*d*’ models. The recollection distribution for THC at retrieval appeared to be somewhat bimodal with a peak shifted leftward from the placebo distribution but also a single peak at 0 (this was much more apparent when emotional conditions were modelled separately; see SM). Although this effect of THC at retrieval resulted in a significant attenuation of recollection (*M* = .17, *SD* = .08, CI = [.03, .33], *p* = .006), another interpretation is that this effect is an artefact of not modeling false memory. In contrast to recollection, there was no significant impact of THC at retrieval on familiarity (*M* = .08, *SD* = .11, CI = [-.12, .30], *p* = .214) or metamemory (*M* = .07, *SD* = .10, CI = [-.11, .27], *p* = .230).

**Figure XI.**
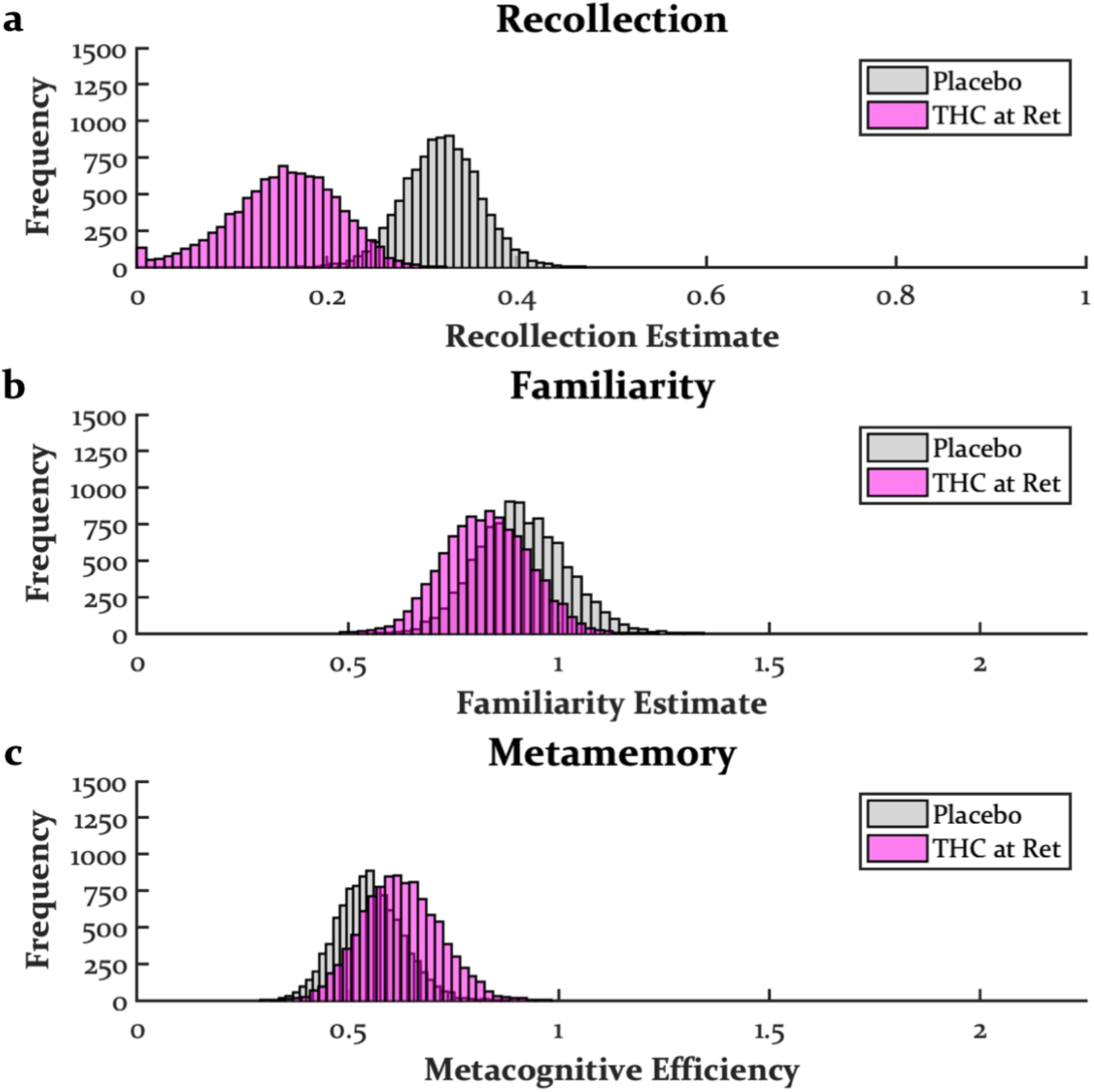
*The Effects Δ^9^-Tetrahydrocannabinol at Retrieval* THC at retrieval impaired recollection, though this was strictly due to an increase in false alarms, suggesting a false memory effect that is not captured in the dual process signal detection model. THC at retrieval had no impact on familiarity or metamemory. Dose used here was 15 mg. THC = Δ^9^-tetrahydrocannabinol, Ret = retrieval.

## Discussion

Assuming that the diverse altered states produced by different drugs are composed of a unique combination of cognitive processes, then a unique pattern of cognitive changes should be produced by different psychoactive drugs. In contrast to standard memory paradigms such as verbal free recall, the findings across the datasets reanalyzed here with computational models of memory confidence show unique patterns of effects across different classes of psychoactive drugs. Sedatives, dissociatives, and cannabinoids at encoding consistently impaired recollection and also familiarity at higher doses. Metamemory was generally unaffected by these drugs at encoding, though there was some evidence for dissociatives and cannabinoids to enhance metamemory. In contrast, psychedelics at encoding impaired recollection and showed a trend of enhancing familiarity but had no effect on metamemory. Stimulants at encoding, retrieval, or consolidation appeared to have little impact on recollection, though they perhaps enhanced familiarity at retrieval. Nevertheless, stimulants consistently impacted metamemory with enhancements when administered at encoding, retrieval, and especially both encoding and retrieval and impairments at consolidation. These findings, as well as those drug manipulations that have yet to be tested, are highlighted in Table II.

**Table II.**
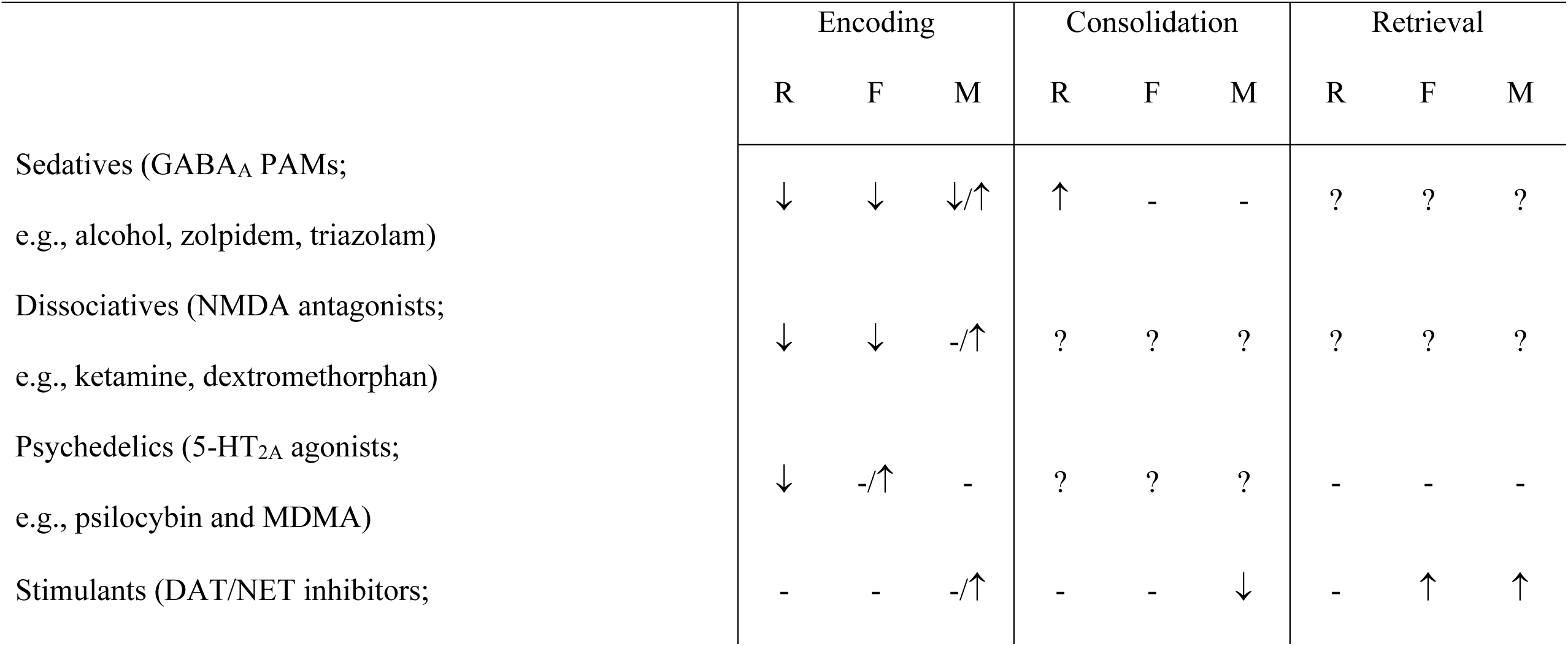
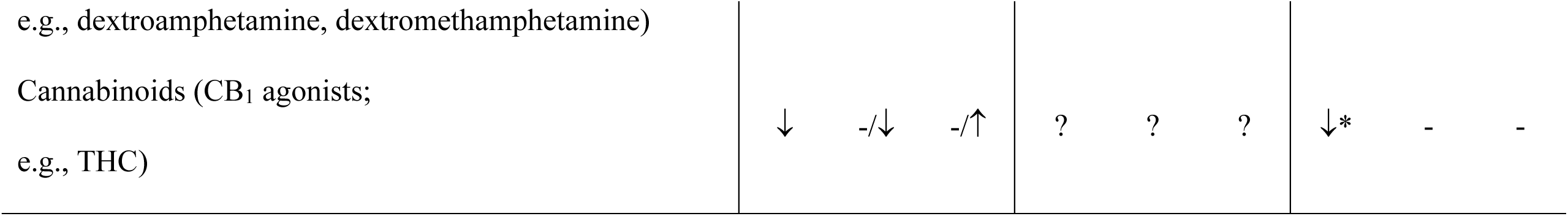
*Summary of Psychoactive Drug Effects* Up and down arrows indicate enhancements and impairments, respectively. A hyphen indicates no effect, and a question mark indicates an unknown effect. Both a hyphen and arrow indicate a trending effect, and both arrows indicate mixed findings. Asterisk indicates potentially inappropriate modeling of a false memory effect. R = recollection, F = familiarity, M = metamemory, PAM = positive allosteric modulator, MDMA = ±3,4-methylenedioxymethamphetamine, DAT = dopamine transporter, NET = norepinephrine transporter, THC = Δ^9^-tetrahydrocannabinol.

|  | Encoding |  |  | Consolidation |  |  | Retrieval |  |  |
| --- | --- | --- | --- | --- | --- | --- | --- | --- | --- |
|  | R | F | M | R | F | M | R | F | M |
| Sedatives (GABA<sub>A</sub> PAMs;<br>e.g., alcohol, zolpidem, triazolam) | ↓ | ↓ | ↓/↑ | ↑ | - | - | ? | ? | ? |
| Dissociatives (NMDA antagonists;<br>e.g., ketamine, dextromethorphan) | ↓ | ↓ | -/↑ | ? | ? | ? | ? | ? | ? |
| Psychedelics (5-HT<sub>2A</sub> agonists;<br>e.g., psilocybin and MDMA) | ↓ | -/↑ | - | ? | ? | ? | - | - | - |
| Stimulants (DAT/NET inhibitors; | - | - | -/↑ | - | - | ↓ | - | ↑ | ↑ |
| e.g., dextroamphetamine, dextromethamphetamine) |  |  |  |  |  |  |  |  |  |
| Cannabinoids (CB<sub>1</sub> agonists;<br>e.g., THC) | ↓ | -/↓ | -/↑ | ? | ? | ? | ↓* | - | - |

Although there were not multiple doses for every individual drug, a strength of this project is that a range of low to high doses of sedatives, dissociatives, psychedelics, and stimulants is represented across datasets. The analyzed data included less than to far greater than standard clinical or recreational doses of these compounds. Within a class of drugs, there were doses that evoked small physiological effects and could be mistaken for place and doses that evoked large physiological effects and difficulties in overall functioning (e.g., following instructions or getting to sleep; see original publications). Across studies, one can infer approximately where a given drug manipulation may be in terms of potency within a class of drugs. For example, one might consider 12.5 mg extended-release zolpidem used in Study 2 to be the lowest dose of a sedative considering it is a standard clinical dose and is released slowly. In contrast, .5 mg/70 kg of triazolam in Study 4 could be considered the highest dose of a sedative, as it is about 4 times greater than a common clinical dose (though .7 and .8 g/kg of alcohol in Study 1 could be comparable, as this would be the equivalent of approximately 4 or 5 drinks consumed in 10 minutes). Regardless of where exactly a given dose of a drug may be placed in terms of its relative potency, we generally observed modulation of the same processes within a drug class without any reliable nonlinearities, yet there were stark deviations between drug classes. For example, across doses, there were no trends for psilocybin and MDMA at encoding to enhance metamemory, whereas stimulants at encoding did appear to enhance metamemory. In contrast, across doses, there were no trends for dextroamphetamine and dextromethamphetamine at encoding to impair recollection, but psychedelics at encoding reliably impaired recollection.

Investigating the idiosyncratic patterns of impairments and enhancements from different drugs or classes of drugs could be useful in the prediction of different drug use and abuse patterns. Moreover, insights regarding the mechanistic make-up of unique subjective states may be borne out of discovering the specific representations impacted and operations executed by different drugs. This latter point, using drugs to understand the mind, has been touted by drug researchers (especially in the study of psychedelics; Yaden *et al*., 2021), yet this promise has been rather empty with most studies demonstrating the reverse; that is, understanding the effects of drugs through the lens of psychology.

Below we discuss similarities and differences between the mnemonic effects of different drugs classes and highlight particularly novel and interesting findings. Specifically, drugs thought to produce global impairments in cognition by inhibiting widely distributed receptors impaired both recollection and familiarity when administered at encoding. In contrast, these same drugs did not seem to impair metamemory using a more refined measure of metacognition than past work. Finally, with the recent resurgence of psychedelic research, the impairment of recollection but potential enhancement of familiarity when psychedelics were administered at encoding has both theoretical and clinical implications for these drugs. We conclude with gaps that remain in our knowledge and make suggestions for future work aiming to assess drug effects on episodic memory and cognition more broadly.

### Receptor Distribution may Determine Drug Effects on Familiarity

In addition to perirhinal cortex, familiarity appears to arise from the activity of brain areas such as occipitotemporal and anterior temporal cortices that process conceptual and perceptual features (Ranganath and Ritchey, 2012; Bastin *et al*., 2019). In contrast to recollection, for which the hippocampus appears to be necessary, familiarity-based encoding may be more robust to drug manipulations because of the multiple neural sources that can contribute to noetic consciousness. Nevertheless, it was found that most sedative manipulations at encoding impaired familiarity, and higher doses of dissociatives and 15 mg of THC at encoding also tended to impair familiarity. Considering that GABA_A_ (Waldvogel *et al*., 2010), NMDA (Kumar, 2015), and CB_1_ (Mackie, 2005) receptors are generally inhibitory and widely distributed throughout the brain, saturation of these receptors may impair recollection and familiarity through inhibition of hippocampal and widespread cortical processing.

Although sedatives, dissociatives, and cannabinoids impaired both recollection and familiarity, there are subjective differences in these amnestic states, perhaps because of different degrees to which each process is modulated. Whereas sedatives and dissociatives can cause periods of complete amnesia (“blackouts” and “K-holes,” respectively), such phenomena are not typically reported with THC. Although CB_1_ receptors are typically inhibitory, there are a population of excitatory CB_1_ receptors in entorhinal cortex (Wang *et al*., 2018), a region that receives projections from parahippocampal and perirhinal cortex and projects to the hippocampus. Thus, THC may enhance signaling to the hippocampus via these receptors, providing a buffer against complete amnestic episodes. Interestingly, 5-HT_2A_ receptors are also found in entorhinal cortex, though they are located on inhibitory interneurons (Deng and Lei, 2008; Bombardi, 2012). Whereas psychedelics may enhance familiarity by activating 5-HT_2A_ receptors on excitatory pyramidal neurons distributed throughout most of the cortex (Aghajanian and Marek, 1997; Martin and Nichols, 2016), psychedelics may impair recollection by shunting information flow to the hippocampus via activation of entorhinal 5-HT_2A_ receptors on inhibitory interneurons. Alternatively, activating 5-HT_2A_ receptors on inhibitory interneurons could disrupt entorhinal cortex’s gating of selective information to the hippocampus (Decurtis and Pare, 2004), thereby inundating hippocampal processing and resulting in catastrophic interference (similar to thalamic gating models of psychedelic drug action; Geyer and Vollenweider, 2008; Doss *et al*., 2021). In contrast to blackouts or K-holes, psychedelics impairing recollection, which is associated with autonoetic consciousness, while sparing familiarity, which is associated with noetic consciousness, is consistent with the phenomenological state of ‘ego dissolution’ from which memories are formed with a noetic quality (Yaden *et al*., 2017) but lack self-relevance (Conway, 2005; Letheby and Gerrans, 2017).

Because familiarity appears to be an aggregate signal of processing fluency throughout sensory and semantic networks (Johnston *et al*., 1985; Wang and Yonelinas, 2012; Ozubko and Yonelinas, 2014), differential inhibition of semantic activation or perceptual processing that results in familiarity impairments may also give rise to unique downstream effects (e.g., conceptually- vs. perceptually-mediated memory distortion; Doss *et al*., 2016, 2019; Doss, Picart, and Gallo, 2018) or even states of consciousness (e.g., noetic vs. anoetic; LeDoux and Lau, 2020). Some evidence suggests that semantic and perceptual processing are generally preserved under the acute effects of sedatives (reviewed in Curran, 1999)(Stewart *et al*., 1996; Hirshman *et al*., 1999; but see Boucart *et al*., 2002), whereas dissociatives (Morgan and Curran, 2006) and THC (Morgan *et al*., 2010; Schafer *et al*., 2012) may acutely impair and enhance semantic activation, respectively. Moreover, as hallucinogens, dissociatives might have idiosyncratic effects on perceptual processing, and this may be true of THC, which is also sometimes classified as a hallucinogen. Even within sedatives, there could be idiosyncratic perceptual effects that differentially impact processes supporting familiarity. In contrast to alcohol and benzodiazepines, zolpidem and other hypnotic non-benzodiazepine drugs are reported to be somewhat hallucinogenic, and sedatives that are GABA_A_ agonists (as opposed to GABA_A_ PAMs) such as muscimol and gaboxadol are hallucinogenic. Such findings might explain why zolpidem was the one manipulation at encoding to have equally or stronger effects on familiarity than recollection, though direct drug comparisons are needed.

### Metamemory is not Easily Impaired

Metamemory, measured here with metacognitive efficiency (i.e., meta-*d*’/*d*’), appeared to be more robust to modulation than observed with previous measures of metamemory, specifically, gamma. Whereas past work with gamma readily found impairments from sedatives and sometimes dissociatives at encoding (Bacon *et al*., 1998; Mintzer and Griffiths, 2003a; b, 2007; Carter *et al*., 2009; Mintzer *et al*., 2010; Kleykamp *et al*., 2012; Carter, Kleykamp, *et al*., 2013; Carter, Reissig, *et al*., 2013), we found little evidence for this. One possibility for the discrepancy between the findings reported here and prior findings using gamma to measure metamemory is because gamma does not consider response bias or overall levels of performance. It is unsurprising that drugs that drastically impair memory encoding through attenuation of both recollection and familiarity also impair gamma, which correlates strongly with memory accuracy. Such impairments of metamemory would also be expected from other metacognitive measures that correlate with overall performance, such as the area under type 2 ROC curves. Although it is possible for one’s memory and metamemory to be simultaneously impaired, it should be possible for one to have a poor memory and recognize some of these impairments (as was found with THC and maybe dissociatives). We did find that alcohol at encoding impaired metamemory for beverage-related stimuli, though it is unclear if this is specific to sedatives or if such an effect would occur from any amnestic drug coupled with a memory test that requires discriminating between stimuli that share overlapping conceptual and perceptual features.

In contrast to past work with sedatives at encoding that found consistent impairments of metamemory measured with gamma, dissociatives at encoding did not consistently impair gamma (Lofwall *et al*., 2006; Carter, Kleykamp, *et al*., 2013; Carter, Reissig, *et al*., 2013). Using the meta-*d*’ model, we instead found inconsistent enhancements when dissociatives were administered at encoding. It is possible that these enhancements from dissociatives were not always apparent or significant because memory was tested on the same day as drug administration when some acute drug effects may have persisted. A recent study found a trend for ketamine at retrieval to impair metamemory (measured with metacognitive efficiency; Lehmann *et al*., 2021), so any enhancement of metamemory from dissociatives at encoding might have been rendered null by lingering drug effects on retrieval. THC at encoding with retrieval 48 hours later (i.e., when drug effects have dissipated) was also found to enhance metamemory despite impairments of both recollection and familiarity. Although the idiosyncratic nature of the study design and our confidence binning procedure could have influenced these results, they demonstrate how metamemory measured with metacognitive efficiency can dissociate drug effects on memory from metamemory. Such a pattern of effects would be unlikely from other metacognitive measures such as gamma and the area under type 2 ROC curves on which the direction of effects is typically determined by overall memory performance. Others have noted similarities in the effects of cannabis and ketamine on memory, as well as how these drugs may model cognitive impairments in schizophrenia (Fletcher and Honey, 2006). Although impairments in recollection and familiarity are also observed in schizophrenia (Libby *et al*., 2013), the potential enhancements of metamemory by THC and ketamine at encoding contrast with impairments found in schizophrenia (Moritz *et al*., 2008; Peters *et al*., 2013; Eifler *et al*., 2015). These effects of THC and ketamine were not “hallucinogen-general,” as metamemory was unaffected by psychedelics, thereby questioning whether these drugs truly reveal the contents of one’s mind (as the etiology of “psychedelic” suggests). Nevertheless, retrospective metamemory is only one aspect of metacognition.

Interestingly, stimulants inconsistently enhanced memory, but they consistently impacted metamemory, a finding that again diverges from metacognitive measures correlated with overall performance. There was tendency for dextroamphetamine and dextromethamphetamine at both encoding and retrieval to enhance metamemory, consistent with 12 mg of methamphetamine (but not higher doses) enhancing metacognition of agency (Kirkpatrick *et al*., 2008). Perhaps most convincing was that when dextroamphetamine was administered at both encoding and retrieval, the metamemory enhancement was strongest. Enhancements of metamemory while drug effects are apparent during both encoding and retrieval may provide one with an online gauge for how well information is stored (e.g., while studying and testing oneself). Even if these drugs do not enhance actual memory encoding or retrieval, as was mostly found here, knowing how well information is stored in memory may be just as useful, allowing one to know how much more they should study. This could partially explain the high incidence of stimulant use by healthy college students studying for exams despite minimal cognitive enhancements in non-clinical populations (Bagot and Kaminer, 2014). In contrast, when dextromethamphetamine was administered at consolidation, there appeared to be an impairment of metamemory. This dataset was, however, somewhat problematic, as the recollection estimates were essentially zero (even in the placebo condition). Such findings are observed in patients with medial temporal lobe amnesia (Yonelinas *et al*., 2010) and false memory manipulations that result in several high confidence false alarms (similar to THC at retrieval). Although this impairment of metamemory should be replicated, the distributions of metamemory estimates were all within the range of the group who received dextromethamphetamine at encoding.

In contrast to the metamemory enhancements from stimulants at retrieval, haloperidol in low doses, which is thought to enhance dopaminergic transmission via antagonism of presynaptic dopamine 2 autoreceptors, was found to impair metamemory (using meta-*d*’ modeling) when administered at retrieval (Clos *et al*., 2019). This study implemented a one-day delay between encoding and retrieval, thus minimizing dopaminergic drug effects on consolidation, which could impair metamemory, as was found in the current report. Although one might speculate that these low doses of haloperidol may in fact attenuate dopaminergic activity, as is typically the rationale for using this drug as an antipsychotic, it was found that this dose of haloperidol increased striatal activity during memory retrieval. Moreover, haloperidol at retrieval also enhanced both recollection and familiarity, and similarly, we found dextroamphetamine at retrieval to enhance familiarity. One explanation for these diverging effects could come from the noradrenergic or even serotonergic effects of dextroamphetamine.

It is worth noting that retrospective metamemory decisions happen at retrieval. Thus, how a drug administered at encoding or consolidation can modulate these decisions when retrieval takes place sober (via proper temporal separation between encoding and retrieval) is unclear. Modulation of the actual memory traces could have some impact on subsequent metamemory, suggesting some non-independence between memory and metamemory, though there is clearly not a one-to-one mapping (i.e., high or low memory accuracy does not necessarily result in high and low metamemory). It might be the case that impairing certain memories or memory features could differentiate them more compared to the case in which one has an abundance of memories. This could result in a relative enhancement of what one knows about an event, as was the trend for dissociatives and THC. In contrast, perhaps some non-specific post-encoding memory enhancements, as would be expected when dextromethamphetamine is administered at consolidation, could have made memories for observed stimuli and other events taking place under stimulant intoxication more confusable. Finally, it is possible that suspecting one had a drug onboard during encoding could impact confidence judgements made on a subsequent day (while sober), possibly making one more cautious, which might improve metamemory independent from actual memory effects. Future work manipulating the content of encoded stimuli to be tested, the content of post-encoding stimuli that could interfere with previously encoded stimuli, and drug expectancy effects are needed to shed light on how metacognitive processes during memory retrieval can be modulated by drug effects that have since dissipated.

### Psychedelic Effects on Memory Support Dual Process Models of Memory

Whereas nearly all drug manipulations that impaired recollection also impaired familiarity at least numerically, we found psilocybin and MDMA at encoding to impair recollection while sparing or even enhancing familiarity. Although these enhancements were mostly trends, some evidence in both humans and mice have found psychedelics to enhance familiarity-like memory when administered at consolidation. Specifically, one study found memory enhancements from LSD administered after multiple rounds of encoding and retrieving object locations (Wießner *et al*., 2022), when memory performance may become more familiarity-dependent. Consistent with these findings, another study found post-encoding (4-bromo-3,6-dimethoxybenzocyclobuten-1-yl)methylamine (TCB-2), a psychedelic 5-HT_2A_ agonist, enhanced novel object recognition (Zhang *et al*., 2013), an analogue of familiarity-based memory dependent on perirhinal cortex. Thus, what was found here could be explained by drug effects persisting into the consolidation window, and a rapid-acting post-encoding psychedelic manipulation (e.g., inhaled DMT) might more precisely target familiarity. Such a facilitation of familiarity-based memories, thought to be supported by cortical regions involved in semantic processing, could disrupt crystallized semantic stores in the cortex, thereby driving new patterns of thought to improve psychiatric disorders such as depression (Carhart-Harris *et al*., 2021; Davis *et al*., 2020).

Despite MDMA’s strong facilitation of monoaminergic transmission, stimulants at encoding did not consistently enhance familiarity (or impair recollection), suggesting that the 5-HT_2A_ agonism of MDMA’s *R*-enantiomer could underlie the similar pattern of mnemonic effects as psilocybin. Consistent with this idea, the amnestic effects of pre-encoding MDMA on verbal free recall can be blocked by 5-HT_2A_ antagonism (van Wel *et al*., 2011). Nevertheless, it is possible that MDMA’s release of the neurohormone oxytocin could have also contributed to the pattern of effects found here, as one study found oxytocin at encoding to simultaneously impair recollection and enhance familiarity (Herzmann *et al*., 2013).

The possibility that 5-HT_2A_ agonists at encoding impair recollection but enhance familiarity is of theoretical relevance to dual process models of memory and perhaps cognition more broadly. One issue that dual process models have had trouble reconciling is that in spite of dual process models fitting well to the data, recollection and familiarity tend to correlate (Pratte and Rouder, 2011). It is typically observed that an effect on one process will be observed in the same direction, at least numerically, on the other process (as was found with most drug manipulations here). However, evolution may have placed constraints on when these processes can completely dissociate, as decoupling between memory systems may produce bizarre cognitive phenomena. That is, recollection may constrain the interpretation of increased familiarity, and an enhancement or preservation of familiarity along with a concurrent impairment in recollection may not necessarily be an overall enhancement *per se*. Semantic activation, which can drive familiarity, can result in feelings of insight (even in the absence of veridical insights; Grimmer *et al*., 2022), and d*éjà vu*, premonition (feeling that one can predict the future), and *presque vu* (feeling that one is on the verge of an insight) are illusory phenomena associated with conditions of high familiarity and low recollection (Cleary *et al*., 2012; Kostic *et al*., 2015; Cleary and Claxton, 2018). Anecdotally, users of psychedelics report having insights under the influence of psychedelics, yet it is unknown whether these insights are tangibly realized in a reliable or frequent fashion. It might be that psychedelics drive a feeling of insight and that these feelings can be misattributed to ideas or imagery conjured up during a psychedelic experience (cf. Laukkonen *et al*., 2022). Moreover, other altered states like daydreaming, sleeping, or even acute alcohol intoxication have all been found to facilitate insight problem-solving (Jarosz *et al*., 2012; Sio *et al*., 2013; Zedelius and Schooler, 2016), suggesting that simply being in different states might facilitate insights (cf. “diversifying experiences” in creative cognition; Damian and Simonton, 2014). Still, it is worth noting that stimulation of the right anterior temporal lobe, a region that supports familiarity (Ranganath and Ritchey, 2012; Bastin *et al*., 2019), facilitates insight problem-solving (Chi and Snyder, 2011; Salvi *et al*., 2020). Therefore, if psychedelics enhance familiarity through the anterior temporal lobe, then it may follow that they could also facilitate insights.

An alternative to strict dual process interpretations of memory could be that certain types of information during psychedelic experiences are more readily processed. For example, familiarity-based memory tends to be associated with objects or ‘entities’, whereas recollection-based memory tends to be associated with scenes, context, and novel bindings of disparate information (Diana *et al*., 2007; Ranganath and Ritchey, 2012; Bastin *et al*., 2019). Interestingly, psychedelics have been found to produce “entity encounters” (illusory social events; Winkelman, 2017; Griffiths *et al*., 2019), an effect potentially driven by activating an “entity representation core system” that supports familiarity (Bastin *et al*., 2019). In addition to the anterior temporal lobe’s involvement in familiarity-based memory, it is particularly important for face processing (Von Der Heide *et al*., 2013; Collins and Olson, 2014), and thus, a possible functional locus for the illusory social events and other phenomena produced by psychedelics.

Another possibility is that 5-HT_2A_ agonism alters the style of information processing such that mnemonic heuristics made at encoding like conceptual associations between otherwise disparate stimuli become less efficiently encoded while global matching based on conceptual and perceptual fluency is facilitated. Consistent with this assertion, some work has found psychedelics to enhance semantic and low-level visual processing (Spitzer *et al*., 1996, 2001). A related idea might be that 5-HT_2A_ agonists may alter the computational processes implemented by the brain. Whereas recollection is thought to rely on a threshold-like process, familiarity is thought to be a strength-based process. Thus, psychedelics may shift information processing from pattern separated autonoetic consciousness to a more continuous style of information processing based on stimulus novelty and familiarity. Indeed, psychedelic experiences are sometimes described as a blurring of the boundaries between the self and the rest of the world. A test of this hypothesis could be to explore how psychedelics differentially impact threshold and strength-based processing in other domains, such as perception using a change detection task (Aly and Yonelinas, 2012; Elfman *et al*., 2014). The same computational models can be applied to such data, and this would provide a test for whether psychedelics have any impact on another domain of metacognition.

### Considerations for Future Work

Although extensive, this review and reanalysis contains many gaps in our knowledge of drug effects, as we were limited by the data that we were able to acquire (data were only retrievable from the laboratories of the current authors). Drugs such as opioids, nicotine, GHB, deliriants, and atypical dissociatives were not studied here. Most of the studies with sedatives and dissociatives administered at encoding did not implement a proper delay between encoding and retrieval to control for drug effects on retrieval. However, these studies did test memory after most subjective effects had dissipated, and the effects were qualitatively similar to effects found when a ≥24-hour was implemented between encoding and retrieval. Moreover, many of the effects found here were robust to other differences in study design (e.g., verbal vs. pictorial stimuli, recognition vs. cued recollection memory tests, 6-point vs. 10-point confidence scales), suggesting some generalizability. Nevertheless, certain findings came from studies with relatively idiosyncratic designs (e.g., emotional and neutral scenes in the case of stimulants, a paradigm with context reinstatement in the case of THC at encoding) that require further replications. Another obvious gap is that most datasets analyzed here did not contain a manipulation of consolidation or retrieval. Although some studies in the past have applied drug manipulations to phases not explored here, in many cases, drug effects were not found (potentially explaining why future work manipulating that phase of memory with that specific drug was not continued). It is possible that more in-depth analyses such as those used here may be able to identify drug effects on memory that were otherwise missed. Finally, state-dependent drug effects are not always reliable, and deeper analyses of such data may reveal which mnemonic processes are in fact modulated. Although we believe it is important to parse selective effects of drugs on encoding, consolidation, and retrieval, it is true that these phases of memory are always interacting during drug experiences. Not only could state-dependent effects take place when memories are both encoded and retrieved during a single drug experience, there could be idiosyncratic drug interactions between how a memory is encoded (e.g., the degree to which it is supported by familiarity and/or recollection), how long it has had time to consolidate, and potential distortions in cognitive and metacognitive processes during retrieval. Exploring such interactions will be important for capturing unique phenomena produced by different psychoactive drugs.

Another interesting prospect for future work is how these drugs impact other metacognitive processes such as prospective metamemory. During memory encoding, could some drugs impair or enhance one’s predictions of their future memory? Prospective metamemory may be particularly impaired with sedatives, as blackouts from these drugs are not typically predictable by the individual. Sedatives, particularly benzodiazepines, are especially used as “roofies” (drugs administered without an individual’s consent, especially in cases of sexual assault). Lack of metacognitive awareness that one’s memories are slipping away into a cognitive abyss may make one particularly susceptible to blackouts. In contrast, prospective metamemory may be a metacognitive process that is enhanced by psychedelics, as “mystical experiences” driven by psychedelics are recollective experiences that are thought to be quite memorable (Griffiths *et al*., 2006). Therefore, perhaps information that survives the recollection impairment produced by psychedelics at encoding can be known with greater certainty.

This report highlights the utility of implementing confidence in tests of episodic memory to identify unique properties of different drugs, but other computational models can be implemented for different types of data to achieve this end. For example, considering the prominent use of verbal free recall in drug studies, a model of temporal context can be applied to these data based on the order of what is recalled (i.e., recall of one word can be used as a cue to recall a temporally proximate word; Howard and Kahana, 2002; Polyn *et al*., 2009). Additionally, we recommend for future work on drugs to use non-standard episodic memory tasks or stimulus manipulations, as these may reveal other unique drug effects. For example, using manipulations of context-dependency, it was found that sedatives at encoding may simply impair context-dependent memory (Reder *et al*., 2013). In contrast, THC at encoding can both impair or enhance context-dependency, sometimes resulting in a magnification of context-dependent memory distortions and other times abolishing it (Doss, Weafer, *et al*., 2020). Such findings have implications for context reinstatement procedures known to boost impoverished memory in “highwitness testimonies” and beg for the study of drug effects on specific mechanisms of false memory (e.g., Doss *et al*., 2016, 2019).

Although this report focused on episodic memory, most areas of cognition have progressed far beyond the standardized paradigms and analyses implemented in most clinical work (many of these tasks developed before computers were widespread). Other potentially interesting areas of study could be visual perception and attention. Whereas alcohol impairs low-level (contrast) but not high-level (motion) visual processing (Weschke and Niedeggen, 2012), the converse is true for psilocybin (Carter *et al*., 2004). Moreover, moderate to high doses of dissociatives have little to no effect on some measures of attention (Morgan *et al*., 2004; Lofwall *et al*., 2006; Honey *et al*., 2008; Carter, Kleykamp, *et al*., 2013; Carter, Reissig, *et al*., 2013) despite sedatives (Zoethout *et al*., 2011), psychedelics (Gouzoulis-Mayfrank *et al*., 2002; Carter *et al*., 2005), and cannabinoids (Broyd *et al*., 2016) consistently impairing attention. With many clinical researchers unable to dedicate their time to a deep understanding of the most modern analyses and paradigms across cognitive domains and most cognitive psychologists unable to run drug studies outside of a hospital, furthering our knowledge of drugs necessitates more interdisciplinary work between these research groups.

## Supporting information

Supplemental analyses of non-drug conditions.

## Acknowledgements

The authors would like to thank Jessica Weafer, Miriam Mintzer, and Michael Ballard for sharing their data. This work is dedicated to Boru Doss who provided invaluable support and passed during the writing of this manuscript.

## Author Contributions

Participated in research design: Doss, Samaha, Barrett, Gallo, Koen Conducted experiments: Doss, Barrett, Griffiths, de Wit, Gallo Contributed new reagents or analytic tools: Doss, Samaha, Koen Performed data analysis: Doss Wrote or contributed to the writing of the manuscript: Doss, Samaha, Barrett, Griffiths, de Wit, Gallo, Koen

## Abbreviations

5-HT_2A_: serotonin 2A receptor
ADHD: attention deficit hyperactivity disorder
AIDS: acquired immunodeficiency syndrome
CB1: cannabinoid 1 receptor
CI: confidence interval
DMT: *N*,*N*-dimethyltryptamine
DPSD: dual process signal detection
GABA: γ-aminobutyric acid
GHB: γ-hydroxybutyric acid
LSD: lysergic acid diethylamide
MDEA: 3,4-methylenedioxyethylamphetamine
MDMA: 3,4-methylenedioxymethamphetamine
MXE: methoxetamine
NMDA: *N*-methyl-d-aspartate
PAM: positive allosteric modulator
ROC: receiver operator characteristic
SM: Supplemental Material
TCB-2: (4-bromo-3,6-dimethoxybenzocyclobuten-1-yl)methylamine
THC: 1′^9^-tetrahydrocannabinol

## Footnotes

*This work was supported by the National Institute on Drug Abuse grants T32DA007209 (M.K.D.), R01DA03889 (R.R.G.), R01DA02812 (H.d.W). J.D.K. was supported by a National Institute on Aging grant (R56AG068149) during preparation of this manuscript. This work was also supported by the Heffter Research Institute, the Steven and Alexandra Cohen Foundation, Tim Ferriss, Blake Mycoskie, Matt Mullenweg, and Craig Nerenberg.

• M.K.D. is an advisor to Ocean Bio Ltd., VCENNA, Inc., and Arcadia Medicine. F.S.B. is an advisor to Wavepaths. R.R.G. is a board member of the Heffter Research Institute. H.d.W. is an advisor to Schedule I Therapeutics, Gilgamesh Pharmaceutics, and PharmAla Biotech. None of these companies had any involvement with the research presented here. J.S., D.A.G., and J.D.K. have no competing interests to declare.

• A subset of the analyses here were presented as posters at the College on Problems of Drug Dependence 81st annual meeting and the Psychonomic Society 60th annual meeting.

