## Supplemental analyses of non-drug conditions. for "Unique Effects of Sedatives, Dissociatives, Psychedelics, Stimulants, and Cannabinoids on Episodic Memory: A Review and Reanalysis of Acute Drug Effects on Recollection, Familiarity, and Metamemory"

### Supplemental Material

Manoj K. Doss[1], Jason Samaha[2], Frederick S. Barrett[1,3,4], Roland R. Griffiths[1,4], Harriet de Wit[5], David A. Gallo[6], & Joshua D. Koen[7]

[1]Department of Psychiatry and Behavioral Sciences, Center for Psychedelic & Consciousness Research

[2]Psychology Department, University of California, Santa Cruz

[3]Department of Psychological and Brain Sciences, Johns Hopkins University

[4]Department of Neuroscience, Johns Hopkins University School of Medicine

[5]Department of Psychiatry and Behavioral Neuroscience, University of Chicago

[6]Department of Psychology, University of Chicago

[7]Department of Psychology, University of Notre Dame

### Study 1: The Effects of Alcohol on Encoding and Consolidation

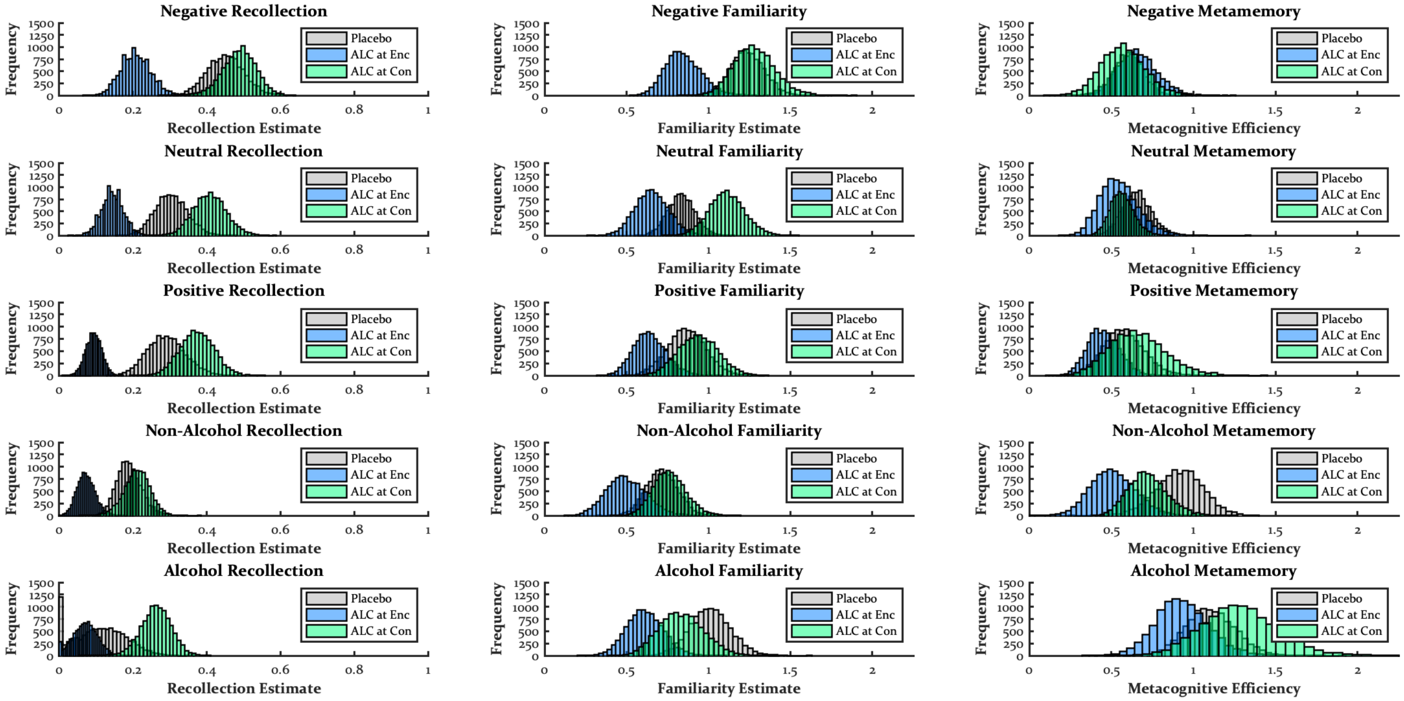


*Recollection Contrasts*

| | PLA vs. Enc | PLA vs. Con | Enc vs. Con |
|---|---|---|---|
| Negative | M = .24, SD = .06, CI = [.12, .36], p = .000 | M = .04, SD = .07, CI = [-.10, .16], p = .288 | M = .28, SD = .06, CI = [.16, .39], p = .000 |
| Neutral | M = .16, SD = .05, CI = [.06, .26], p = .001 | M = .10, SD = .06, CI = [-.02, .23], p = .046 | M = .26, SD = .05, CI = [.16, .37], p = .000 |
| Positive | M = .20, SD = .06, CI = [.09, .31], p = .000 | M = .08, SD = .07, CI = [-.05, .22], p = .118 | M = .28, SD = .05, CI = [.18, .38], p = .000 |

| | | | |
|---|---|---|---|
| Non-Alcohol | M = .12, SD = .04, CI = [.04, .20], p = .001 | M = .02, SD = .05, CI = [-.08, .11], p = .318 | M = .14, SD = .04, CI = [.05, .22], p = .001 |
| Alcohol | M = .04, SD = .08, CI = [-.10, .18], p = .323 | M = .15, SD = .08, CI = [-.00, .30], p = .029 | M = .19, SD = .05, CI = [.09, .29], p = .001 |

| | Negative vs. Neutral | Negative vs. Positive | Negative vs. Non-Alcohol | Negative vs. Alcohol | Neutral vs. Positive | Neutral vs. Non-Alcohol | Neutral vs. Alcohol | Positive vs. Non-Alcohol | Positive vs. Alcohol | Non-Alcohol vs. Alcohol |
|---|---|---|---|---|---|---|---|---|---|---|
| Placebo | M = .15, SD = .03, CI = [.08, .22], p = .000 | M = .16, SD = .05, CI = [.07, .25], p = .000 | M = .26, SD = .05, CI = [.17, .37], p = .000 | M = .34, SD = .08, CI = [.20, .49], p = .000 | M = .01, SD = .04, CI = [-.07, .10], p = .381 | M = .11, SD = .04, CI = [.04, .20], p = .001 | M = .19, SD = .07, CI = [.07, .33], p = .000 | M = .10, SD = .04, CI = [.03, .18], p = .003 | M = .18, SD = .08, CI = [.04, .33], p = .005 | M = .08, SD = .07, CI = [-.05, .21], p = .137 |
| ALC at Enc | M = .06, SD = .03, CI = [.01, .12], p = .006 | M = .11, SD = .03, CI = [.06, .18], p = .000 | M = .14, SD = .04, CI = [.07, .22], p = .000 | M = .14, SD = .05, CI = [.06, .23], p = .000 | M = .05, SD = .02, CI = [.01, .10], p = .005 | M = .07, SD = .03, CI = [.02, .13], p = .002 | M = .07, SD = .04, CI = [.00, .16], p = .018 | M = .02, SD = .03, CI = [-.03, .08], p = .220 | M = .02, SD = .04, CI = [-.04, .10], p = .264 | M = .00, SD = .03, CI = [-.06, .07], p = .485 |
| ALC at Con | M = .08, SD = .04, CI = [.00, .16], p = .021 | M = .11, SD = .04, CI = [.03, .20], p = .005 | M = .28, SD = .04, CI = [.19, .37], p = .000 | M = .23, SD = .05, CI = [.14, .33], p = .000 | M = .03, SD = .03, CI = [-.02, .10], p = .140 | M = .20, SD = .04, CI = [.13, .27], p = .000 | M = .15, SD = .05, CI = [.06, .25], p = .000 | M = .16, SD = .04, CI = [.08, .25], p = .000 | M = .11, SD = .05, CI = [.02, .21], p = .012 | M = .05, SD = .05, CI = [-.05, .15], p = .140 |

*Familiarity Contrasts*

| | PLA vs. Enc | PLA vs. Con | Enc vs. Con |
|---|---|---|---|
| Negative | M = .39, SD = .15, CI = [.09, .68], p = .006 | M = .04, SD = .16, CI = [-.26, .37], p = .402 | M = .43, SD = .17, CI = [.10, .76], p = .005 |
| Neutral | M = .18, SD = .13, CI = [-.08, .43], p = .085 | M = .28, SD = .13, CI = [.01, .54], p = .020 | M = .46, SD = .15, CI = [.17, .75], p = .001 |
| Positive | M = .22, SD = .15, CI = [-.07, .52], p = .074 | M = .07, SD = .17, CI = [-.25, .40], p = .335 | M = .30, SD = .16, CI = [-.01, .59], p = .027 |
| Non-Alcohol | M = .22, SD = .15, CI = [-.09, .51], p = .079 | M = .05, SD = .14, CI = [-.22, .32], p = .374 | M = .26, SD = .16, CI = [-.04, .57], p = .048 |
| Alcohol | M = .40, SD = .17, CI = [.07, .73], p = .007 | M = .18, SD = .18, CI = [-.19, .53], p = .169 | M = .23, SD = .17, CI = [-.11, .57], p = .096 |

| | Negative vs. Neutral | Negative vs. Positive | Negative vs. Non-Alcohol | Negative vs. Alcohol | Neutral vs. Positive | Neutral vs. Non-Alcohol | Neutral vs. Alcohol | Positive vs. Non-Alcohol | Positive vs. Alcohol | Non-Alcohol vs. Alcohol |
|---|---|---|---|---|---|---|---|---|---|---|
| Placebo | M = .40, SD = .09, CI = [.22, .58], p = .000 | M = .37, SD = .09, CI = [.18, .54], p = .000 | M = .52, SD = .11, CI = [.32, .73], p = .000 | M = .23, SD = .16, CI = [-.10, .53], p = .087 | M = .03, SD = .10, CI = [-.16, .25], p = .392 | M = .12, SD = .08, CI = [-.04, .28], p = .077 | M = .18, SD = .14, CI = [-.08, .46], p = .099 | M = .15, SD = .12, CI = [-.08, .38], p = .098 | M = .15, SD = .17, CI = [-.16, .50], p = .208 | M = .30, SD = .16, CI = [-.01, .61], p = .031 |
| ALC at Enc | M = .19, SD = .09, CI = [.01, .37], p = .022 | M = .21, SD = .09, CI = [.03, .39], p = .006 | M = .35, SD = .10, CI = [.15, .54], p = .001 | M = .24, SD = .11, CI = [.02, .47], p = .016 | M = .01, SD = .12, CI = [-.22, .25], p = .453 | M = .16, SD = .11, CI = [-.07, .37], p = .080 | M = .04, SD = .12, CI = [-.19, .27], p = .352 | M = .14, SD = .08, CI = [-.03, .29], p = .047 | M = .03, SD = .12, CI = [-.19, .26], p = .402 | M = .11, SD = .12, CI = [-.14, .34], p = .178 |
| ALC at Con | M = .17, SD = .16, CI = [-.14, .47], p = .145 | M = .34, SD = .16, CI = [.03, .69], p = .011 | M = .52, SD = .15, CI = [.23, .80], p = .000 | M = .44, SD = .15, CI = [.14, .75], p = .003 | M = .18, SD = .11, CI = [-.04, .38], p = .052 | M = .35, SD = .13, CI = [.09, .60], p = .004 | M = .28, SD = .15, CI = [-.02, .56], p = .035 | M = .17, SD = .11, CI = [-.04, .41], p = .060 | M = .10, SD = .14, CI = [-.20, .35], p = .226 | M = .07, SD = .15, CI = [-.19, .37], p = .320 |

*Metamemory Contrasts*

| | PLA vs. Enc | PLA vs. Con | Enc vs. Con |
|---|---|---|---|
| Negative | M = .05, SD = .15, CI = [-.24, .35], p = .382 | M = .04, SD = .16, CI = [-.28, .34], p = .401 | M = .09, SD = .18, CI = [-.27, .44], p = .314 |
| Neutral | M = .11, SD = .15, CI = [-.20, .39], p = .227 | M = .09, SD = .11, CI = [-.13, .32], p = .212 | M = .02, SD = .14, CI = [-.28, .27], p = .432 |
| Positive | M = .13, SD = .15, CI = [-.17, .43], p = .212 | M = .08, SD = .21, CI = [-.32, .50], p = .351 | M = .21, SD = .19, CI = [-.15, .62], p = .136 |
| Non-Alcohol | M = .41, SD = .20, CI = [.01, .78], p = .023 | M = .19, SD = .19, CI = [-.19, .54], p = .148 | M = .21, SD = .18, CI = [-.13, .56], p = .117 |
| Alcohol | M = .17, SD = .22, CI = [-.27, .60], p = .212 | M = .18, SD = .27, CI = [-.34, .74], p = .254 | M = .36, SD = .28, CI = [-.19, .93], p = .102 |

| | Negative vs. Neutral | Negative vs. Positive | Negative vs. Non-Alcohol | Negative vs. Alcohol | Neutral vs. Positive | Neutral vs. Non-Alcohol | Neutral vs. Alcohol | Positive vs. Non-Alcohol | Positive vs. Alcohol | Non-Alcohol vs. Alcohol |
|---|---|---|---|---|---|---|---|---|---|---|

| | | | | | | | | | | |
|---|---|---|---|---|---|---|---|---|---|---|
| Placebo | M = .05, SD = .10, CI = [-.15, .23], p = .300 | M = .03, SD = .17, CI = [-.31, .33], p = .409 | M = .31, SD = .17, CI = [-.06, .62], p = .046 | M = .50, SD = .17, CI = [.18, .83], p = .000 | M = .08, SD = .16, CI = [-.24, .38], p = .305 | M = .26, SD = .15, CI = [-.05, .53], p = .048 | M = .45, SD = .19, CI = [.09, .83], p = .009 | M = .34, SD = .19, CI = [-.05, .72], p = .042 | M = .53, SD = .17, CI = [.20, .87], p = .001 | M = .19, SD = .21, CI = [-.23, .60], p = .183 |
| ALC at Enc | M = .11, SD = .12, CI = [-.14, .34], p = .183 | M = .20, SD = .14, CI = [-.06, .48], p = .070 | M = .15, SD = .15, CI = [-.15, .45], p = .160 | M = .28, SD = .23, CI = [-.17, .74], p = .110 | M = .10, SD = .15, CI = [-.17, .42], p = .272 | M = .04, SD = .16, CI = [-.25, .39], p = .424 | M = .38, SD = .22, CI = [-.06, .80], p = .042 | M = .06, SD = .15, CI = [-.22, .35], p = .350 | M = .48, SD = .14, CI = [.20, .76], p = .001 | M = .42, SD = .17, CI = [.07, .76], p = .011 |
| ALC at Con | M = .01, SD = .13, CI = [-.24, .25], p = .484 | M = .09, SD = .16, CI = [-.22, .42], p = .297 | M = .15, SD = .11, CI = [-.07, .36], p = .085 | M = .72, SD = .20, CI = [.36, 1.13], p = .000 | M = .10, SD = .17, CI = [-.20, .45], p = .294 | M = .16, SD = .11, CI = [-.05, .40], p = .068 | M = .72, SD = .22, CI = [.32, 1.18], p = .000 | M = .06, SD = .16, CI = [-.24, .37], p = .348 | M = .63, SD = .21, CI = [.22, 1.07], p = .000 | M = .56, SD = .23, CI = [.15, 1.04], p = .003 |

### Study 2: The Effects of Zolpidem on Encoding

*First Memory Test*

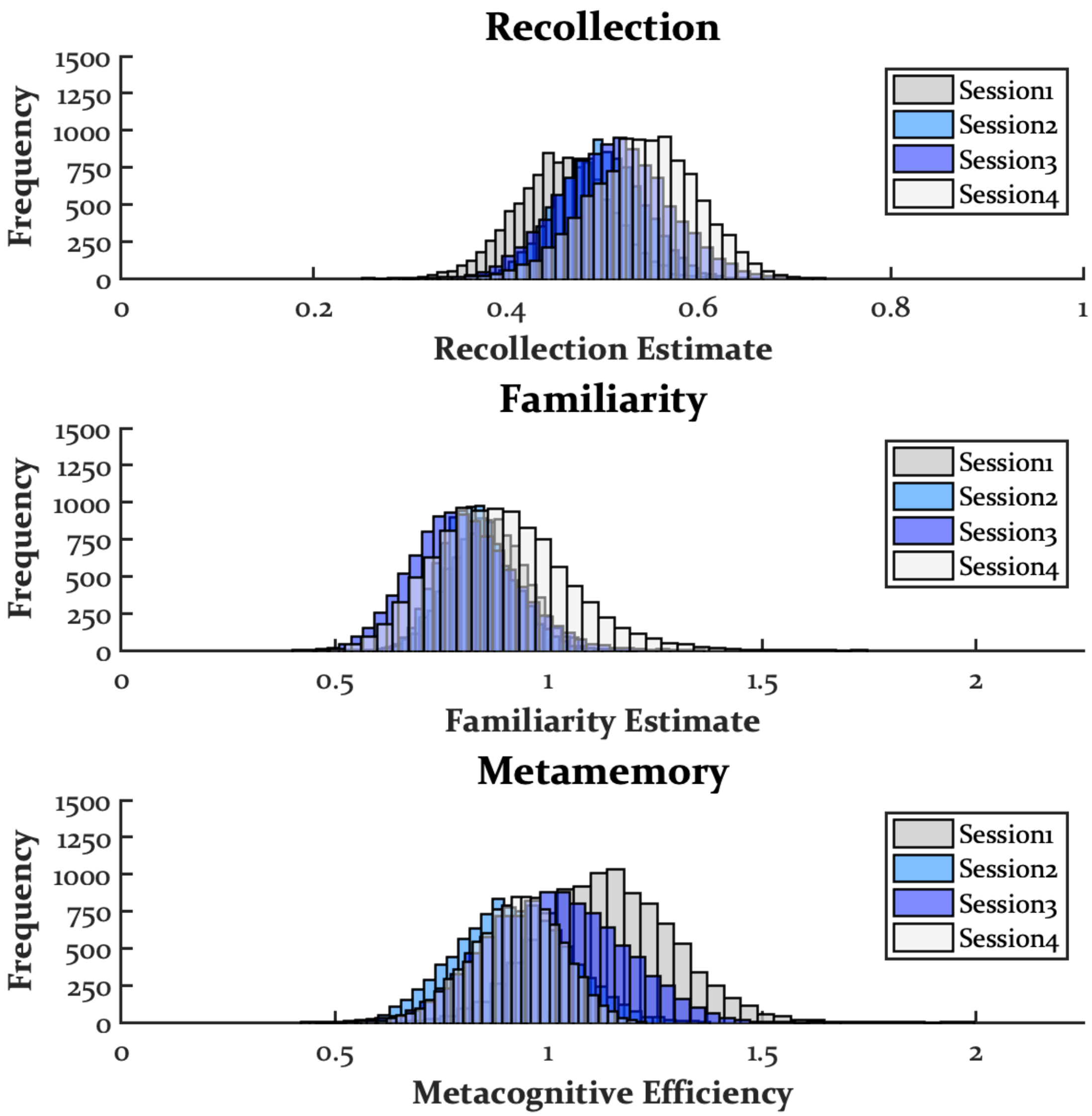
Recollection
Frequency
1500
1250
1000
750
500
250
0
0
0.2
0.4
0.6
0.8
1
Recollection Estimate
Session1
Session2
Session3
Session4
Familiarity
Frequency
1500
1250
1000
750
500
250
0
0
0.5
1
1.5
2
Familiarity Estimate
Session1
Session2
Session3
Session4
Metamemory
Frequency
1500
1250
1000
750
500
250
0
0
0.5
1
1.5
2
Metacognitive Efficiency
Session1
Session2
Session3
Session4

| | Session1 vs. Session2 | Session1 vs. Session3 | Session1 vs. Session4 | Session2 vs. Session3 | Sesison2 vs. Session4 | Session3 vs. Session4 |
|---|---|---|---|---|---|---|
| Recollection | M = .04, SD = .05, CI = [-.06, .13], p = .203 | M = .05, SD = .06, CI = [-.06, .16], p = .177 | M = .08, SD = .06, CI = [-.04, .19], p = .076 | M = .01, SD = .05, CI = [-.09, .12], p = .388 | M = .04, SD = .05, CI = [-.05, .13], p = .172 | M = .03, SD = .06, CI = [-.09, .13], p = .273 |
| Familiarity | M = .03, SD = .09, CI = [-.16, .21], p = .379 | M = .06, SD = .13, CI = [-.20, .33], p = .331 | M = .03, SD = .14, CI = [-.24, .33], p = .433 | M = .03, SD = .10, CI = [-.17, .23], p = .380 | M = .06, SD = .17, CI = [-.25, .41], p = .379 | M = .09, SD = .22, CI = [-.33, .54], p = .355 |
| Metamemory | M = .22, SD = .16, CI = [-.07, .54], p = .077 | M = .12, SD = .17, CI = [-.19, .46], p = .237 | M = .21, SD = .18, CI = [-.11, .57], p = .107 | M = .10, SD = .18, CI = [-.27, .44], p = .282 | M = .01, SD = .14, CI = [-.28, .28], p = .467 | M = .09, SD = .15, CI = [-.19, .39], p = .277 |

*Fourth Memory Test*

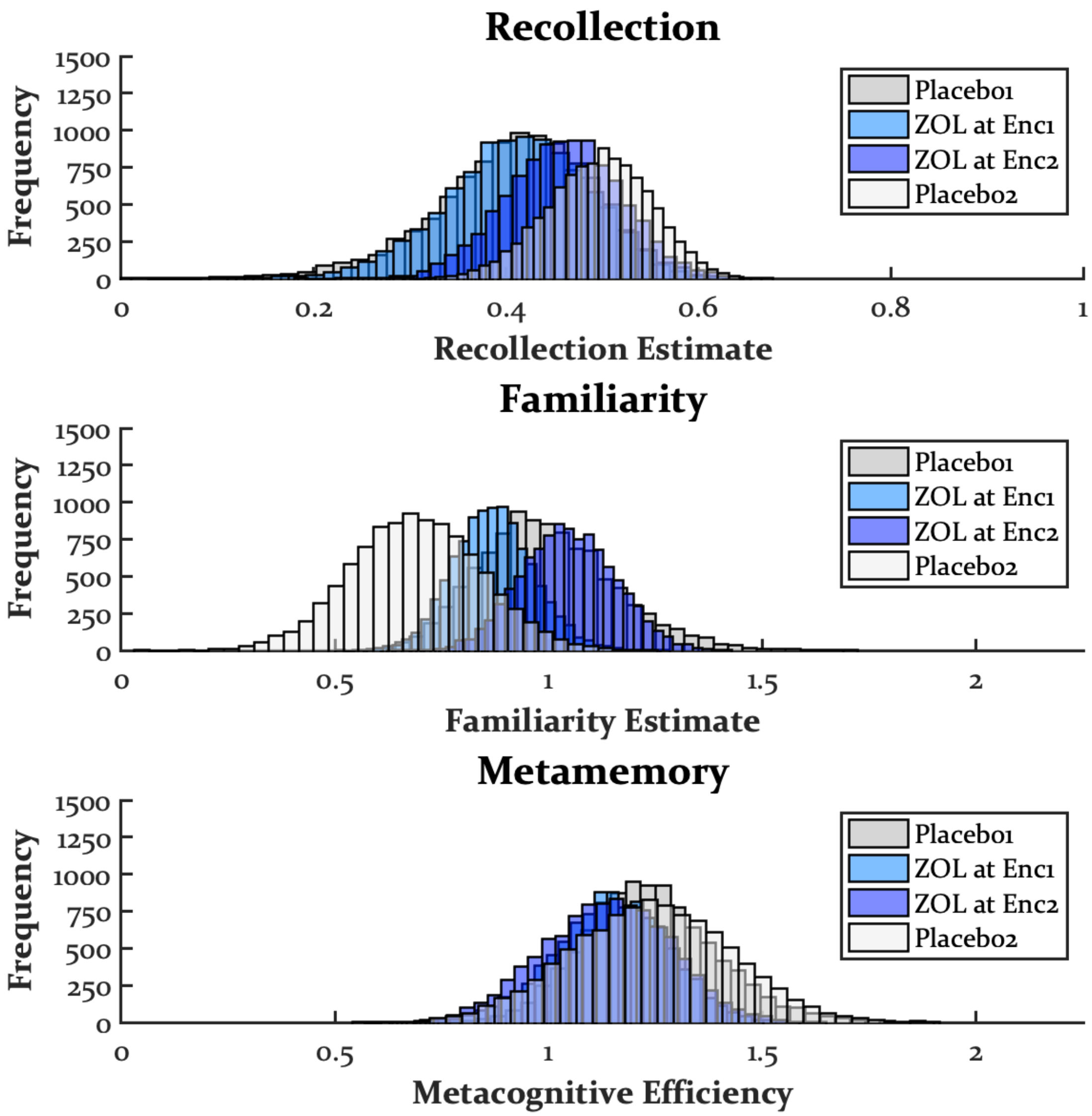

Recollection
Frequency
1500
1250
1000
750
500
250
0
0
0.2
0.4
0.6
0.8
1
Recollection Estimate
Placebo1
ZOL at Enc1
ZOL at Enc2
Placebo2
Familiarity
Frequency
0
0.5
1
1.5
2
Familiarity Estimate
Metamemory
Frequency
Metacognitive Efficiency

| | PLA1 vs. Enc1 | PLA1 vs. Enc2 | PLA1 vs PLA2 | Enc1 vs. Enc2 | Enc1 vs. PLA2 | Enc2 vs. PLA2 |
|---|---|---|---|---|---|---|
| Recollection | M = .01, SD = .10, CI = [-.16, .22], p = .492 | M = .05, SD = .06, CI = [-.04, .18], p = .148 | M = .09, SD = .07, CI = [-.02, .25], p = .065 | M = .05, SD = .06, CI = [-.08, .17], p = .232 | M = .08, SD = .06, CI = [-.04, .19], p = .085 | M = .04, SD = .04, CI = [-.05, .11], p = .169 |
| Familiarity | M = .12, SD = .16, CI = [-.17, .46], p = .224 | M = .05, SD = .15, CI = [-.26, .34], p = .358 | M = .31, SD = .24, CI = [-.13, .81], p = .093 | M = .17, SD = .14, CI = [-.10, .46], p = .111 | M = .18, SD = .17, CI = [-.17, .50], p = .147 | M = .36, SD = .19, CI = [-.01, .73], p = .028 |
| Metamemory | M = .09, SD = .19, CI = [-.27, .46], p = .312 | M = .10, SD = .18, CI = [-.24, .48], p = .286 | M = .00, SD = .20, CI = [-.39, .42], p = .510 | M = .01, SD = .17, CI = [-.34, .33], p = .468 | M = .09, SD = .17, CI = [-.25, .41], p = .287 | M = .10, SD = .14, CI = [-.17, .38], p = .231 |

### Study 6: The Effects of MDMA on Encoding and Retrieval

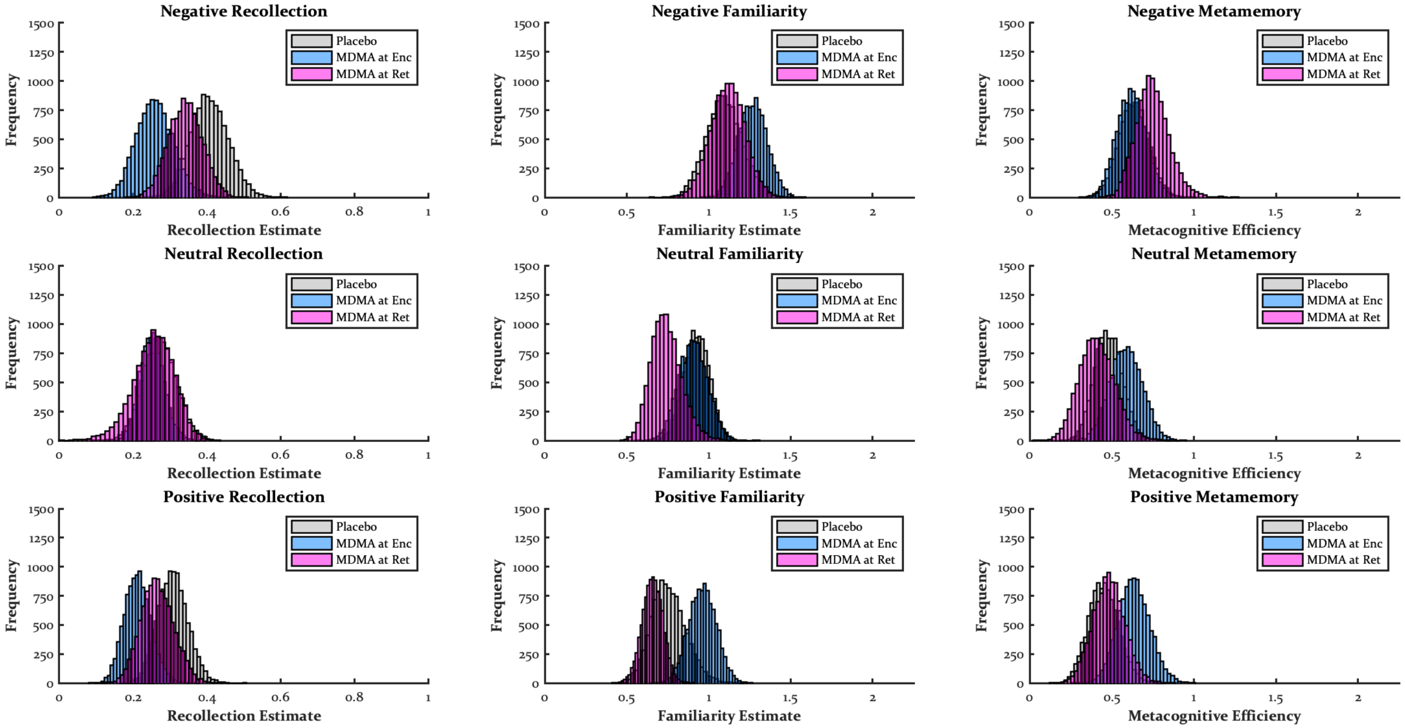


*Recollection Contrasts*

| | PLA vs. Enc | PLA vs. Ret | Enc vs. Ret |
|---|---|---|---|
| Negative | M = .15, SD = .07, CI = [.01, .29], p = .017 | M = .07, SD = .07, CI = [-.06, .20], p = .166 | M = .08, SD = .07, CI = [-.05, .21], p = .112 |
| Neutral | M = .02, SD = .06, CI = [-.09, .13], p = .350 | M = .02, SD = .07, CI = [-.12, .17], p = .411 | M = .00, SD = .06, CI = [-.13, .12], p = .463 |
| Positive | M = .10, SD = .05, CI = [-.01, .20], p = .033 | M = .04, SD = .06, CI = [-.07, .16], p = .236 | M = .05, SD = .06, CI = [-.06, .17], p = .167 |

| | Negative vs. Neutral | Negative vs. Positive | Neutral vs. Positive |
|---|---|---|---|

| | | | |
|---|---|---|---|
| Placebo | M = .13, SD = .04, CI = [.06, .22], p = .000 | M = .10, SD = .04, CI = [.02, .18], p = .005 | M = .04, SD = .03, CI = [-.02, .09], p = .122 |
| MDMA at Enc | M = .01, SD = .04, CI = [-.07, .08], p = .405 | M = .05, SD = .03, CI = [-.01, .10], p = .055 | M = .04, SD = .02, CI = [.00, .08], p = .015 |
| MDMA at Ret | M = .09, SD = .05, CI = [.01, .19], p = .017 | M = .08, SD = .04, CI = [-.00, .15], p = .027 | M = .01, SD = .04, CI = [-.06, .10], p = .432 |

*Familiarity Contrasts*

| | PLA vs. Enc | PLA vs. Ret | Enc vs. Ret |
|---|---|---|---|
| Negative | M = .17, SD = .14, CI = [-.10, .44], p = .101 | M = .02, SD = .15, CI = [-.29, .31], p = .436 | M = .15, SD = .13, CI = [-.11, .42], p = .125 |
| Neutral | M = .02, SD = .12, CI = [-.21, .25], p = .423 | M = .19, SD = .12, CI = [-.05, .41], p = .064 | M = .16, SD = .13, CI = [-.10, .40], p = .100 |
| Positive | M = .21, SD = .13, CI = [-.06, .45], p = .065 | M = .10, SD = .12, CI = [-.12, .35], p = .206 | M = .31, SD = .10, CI = [.10, .51], p = .001 |

| | Negative vs. Neutral | Negative vs. Positive | Neutral vs. Positive |
|---|---|---|---|
| Placebo | M = .17, SD = .10, CI = [-.03, .36], p = .045 | M = .34, SD = .10, CI = [.14, .53], p = .001 | M = .17, SD = .11, CI = [-.05, .39], p = .065 |
| MDMA at Enc | M = .36, SD = .08, CI = [.21, .52], p = .000 | M = .31, SD = .09, CI = [.12, .49], p = .001 | M = .06, SD = .08, CI = [-.09, .20], p = .218 |
| MDMA at Ret | M = .38, SD = .10, CI = [.16, .56], p = .000 | M = .46, SD = .10, CI = [.25, .66], p = .000 | M = .08, SD = .10, CI = [-.10, .28], p = .194 |

*Metamemory Contrasts*

| | PLA vs. Enc | PLA vs. Ret | Enc vs. Ret |
|---|---|---|---|
| Negative | M = .01, SD = .13, CI = [-.24, .26], p = .454 | M = .12, SD = .14, CI = [-.14, .39], p = .196 | M = .13, SD = .13, CI = [-.12, .40], p = .166 |
| Neutral | M = .11, SD = .13, CI = [-.15, .36], p = .203 | M = .07, SD = .14, CI = [-.20, .33], p = .298 | M = .18, SD = .14, CI = [-.11, .45], p = .112 |
| Positive | M = .19, SD = .13, CI = [-.07, .45], p = .078 | M = .03, SD = .13, CI = [-.23, .29], p = .401 | M = .15, SD = .14, CI = [-.11, .42], p = .126 |

| | Negative vs. Neutral | Negative vs. Positive | Neutral vs. Positive |
|---|---|---|---|

| | | | |
|---|---|---|---|
| Placebo | M = .15, SD = .11, CI = [-.06, .37], p = .086 | M = .19, SD = .12, CI = [-.06, .42], p = .066 | M = .03, SD = .12, CI = [-.19, .27], p = .397 |
| MDMA at Enc | M = .03, SD = .09, CI = [-.15, .21], p = .358 | M = .02, SD = .11, CI = [-.21, .24], p = .443 | M = .05, SD = .11, CI = [-.16, .28], p = .340 |
| MDMA at Ret | M = .34, SD = .11, CI = [.13, .56], p = .001 | M = .27, SD = .13, CI = [.02, .54], p = .016 | M = .07, SD = .14, CI = [-.21, .34], p = .296 |

### Study 7: The Effects of Dextroamphetamine on Encoding, Retrieval, and Both

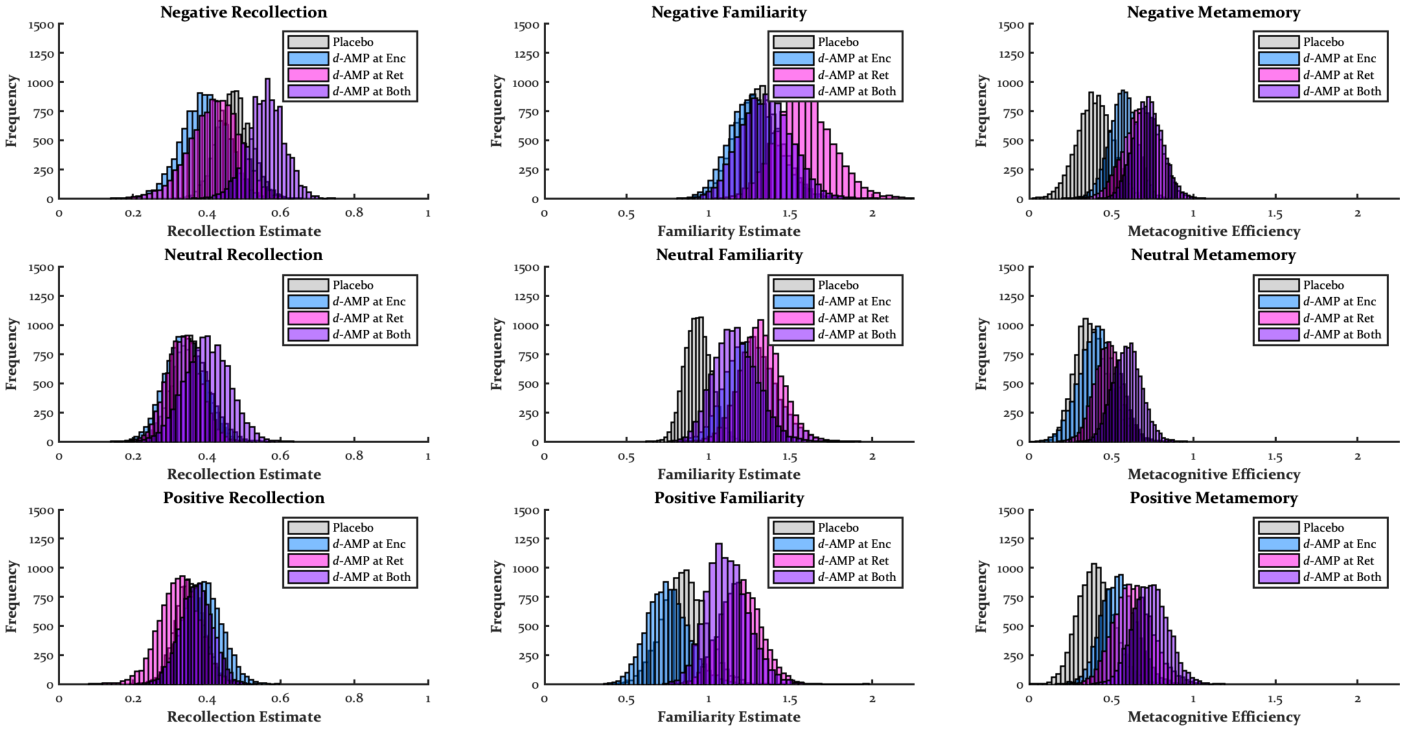


#### *Recollection*

| | PLA vs. Enc | PLA vs. Ret | PLA vs. Both | Enc vs. Ret | Enc vs. Both | Ret vs. Both |
|---|---|---|---|---|---|---|
| Negative | M = .08, SD = .08, CI = [-.06, .23], p = .135 | M = .05, SD = .08, CI = [-.10, .22], p = .264 | M = .09, SD = .07, CI = [-.05, .21], p = .096 | M = .03, SD = .09, CI = [-.15, .20], p = .369 | M = .17, SD = .08, CI = [.02, .32], p = .011 | M = .14, SD = .08, CI = [-.02, .31], p = .042 |
| Neutral | M = .01, SD = .07, CI = [-.14, .15], p = .438 | M = .02, SD = .06, CI = [-.11, .14], p = .399 | M = .05, SD = .08, CI = [-.10, .20], p = .271 | M = .01, SD = .07, CI = [-.13, .15], p = .470 | M = .06, SD = .08, CI = [-.11, .21], p = .244 | M = .06, SD = .08, CI = [-.08, .21], p = .202 |
| Positive | M = .03, SD = .06, CI = [-.09, .16], p = .304 | M = .03, SD = .07, CI = [-.09, .16], p = .334 | M = .02, SD = .06, CI = [-.10, .13], p = .381 | M = .06, SD = .08, CI = [-.08, .21], p = .202 | M = .02, SD = .07, CI = [-.12, .15], p = .408 | M = .05, SD = .07, CI = [-.09, .19], p = .250 |

| | Negative vs. Neutral | Negative vs. Positive | Neutral vs. Positive |
|---|---|---|---|
| Placebo | M = .12, SD = .03, CI = [.05, .19], p = .000 | M = .12, SD = .03, CI = [.06, .19], p = .000 | M = .00, SD = .03, CI = [-.07, .07], p = .498 |
| *d*-AMP at Enc | M = .05, SD = .05, CI = [-.05, .14], p = .147 | M = .01, SD = .06, CI = [-.12, .12], p = .462 | M = .04, SD = .04, CI = [-.03, .13], p = .144 |
| *d*-AMP at Ret | M = .09, SD = .06, CI = [-.05, .20], p = .092 | M = .10, SD = .06, CI = [-.03, .23], p = .059 | M = .01, SD = .06, CI = [-.08, .13], p = .433 |
| *d*-AMP at Both | M = .16, SD = .03, CI = [.09, .23], p = .000 | M = .19, SD = .05, CI = [.10, .28], p = .000 | M = .03, SD = .05, CI = [-.07, .14], p = .293 |

*Familiarity*

| | PLA vs. Enc | PLA vs. Ret | PLA vs. Both | Enc vs. Ret | Enc vs. Both | Ret vs. Both |
|---|---|---|---|---|---|---|
| Negative | M = .02, SD = .20, CI = [-.37, .40], p = .453 | M = .28, SD = .22, CI = [-.12, .72], p = .090 | M = .05, SD = .20, CI = [-.35, .45], p = .406 | M = .31, SD = .22, CI = [-.13, .75], p = .082 | M = .07, SD = .21, CI = [-.35, .48], p = .361 | M = .23, SD = .23, CI = [-.21, .70], p = .151 |
| Neutral | M = .32, SD = .17, CI = [.01, .66], p = .022 | M = .39, SD = .15, CI = [.10, .68], p = .004 | M = .23, SD = .15, CI = [-.06, .54], p = .067 | M = .06, SD = .20, CI = [-.33, .44], p = .369 | M = .10, SD = .20, CI = [-.29, .49], p = .311 | M = .16, SD = .18, CI = [-.20, .51], p = .193 |
| Positive | M = .10, SD = .18, CI = [-.25, .45], p = .289 | M = .34, SD = .18, CI = [-.01, .68], p = .028 | M = .26, SD = .18, CI = [-.08, .62], p = .068 | M = .44, SD = .17, CI = [.10, .79], p = .006 | M = .36, SD = .18, CI = [.02, .72], p = .019 | M = .08, SD = .17, CI = [-.27, .42], p = .316 |

| | Negative vs. Neutral | Negative vs. Positive | Neutral vs. Positive |
|---|---|---|---|
| Placebo | M = .37, SD = .13, CI = [.10, .61], p = .003 | M = .45, SD = .12, CI = [.22, .68], p = .000 | M = .08, SD = .11, CI = [-.13, .30], p = .219 |

| | | | |
|---|---|---|---|
| *d*-AMP at Enc | M = .02, SD = .12, CI = [-.22, .25], p = .425 | M = .53, SD = .13, CI = [.29, .78], p = .000 | M = .51, SD = .13, CI = [.26, .75], p = .000 |
| *d*-AMP at Ret | M = .27, SD = .12, CI = [.06, .52], p = .005 | M = .39, SD = .15, CI = [.11, .68], p = .003 | M = .13, SD = .13, CI = [-.12, .37], p = .163 |
| *d*-AMP at Both | M = .19, SD = .13, CI = [-.07, .44], p = .080 | M = .24, SD = .11, CI = [.04, .46], p = .009 | M = .05, SD = .10, CI = [-.14, .24], p = .309 |

*Metamemory*

| | PLA vs. Enc | PLA vs. Ret | PLA vs. Both | Enc vs. Ret | Enc vs. Both | Ret vs. Both |
|---|---|---|---|---|---|---|
| Negative | M = .17, SD = .13, CI = [-.09, .43], p = .099 | M = .29, SD = .15, CI = [-.00, .58], p = .026 | M = .33, SD = .14, CI = [.06, .60], p = .007 | M = .11, SD = .14, CI = [-.15, .38], p = .195 | M = .16, SD = .12, CI = [-.07, .40], p = .096 | M = .04, SD = .14, CI = [-.22, .32], p = .393 |
| Neutral | M = .03, SD = .15, CI = [-.29, .31], p = .412 | M = .10, SD = .14, CI = [-.18, .35], p = .223 | M = .21, SD = .14, CI = [-.07, .47], p = .065 | M = .07, SD = .14, CI = [-.20, .34], p = .300 | M = .19, SD = .14, CI = [-.08, .46], p = .088 | M = .11, SD = .12, CI = [-.12, .34], p = .173 |
| Positive | M = .14, SD = .15, CI = [-.15, .43], p = .175 | M = .23, SD = .16, CI = [-.09, .55], p = .074 | M = .33, SD = .15, CI = [.04, .63], p = .014 | M = .09, SD = .16, CI = [-.22, .39], p = .290 | M = .19, SD = .15, CI = [-.10, .49], p = .097 | M = .11, SD = .16, CI = [-.21, .41], p = .247 |

| | Negative vs. Neutral | Negative vs. Positive | Neutral vs. Positive |
|---|---|---|---|
| Placebo | M = .01, SD = .14, CI = [-.27, .27], p = .460 | M = .01, SD = .11, CI = [-.19, .22], p = .476 | M = .02, SD = .13, CI = [-.24, .28], p = .447 |
| *d*-AMP at Enc | M = .16, SD = .11, CI = [-.06, .39], p = .081 | M = .02, SD = .12, CI = [-.22, .25], p = .417 | M = .13, SD = .12, CI = [-.08, .38], p = .126 |
| *d*-AMP at Ret | M = .20, SD = .13, CI = [-.04, .46], p = .056 | M = .05, SD = .16, CI = [-.28, .34], p = .362 | M = .15, SD = .13, CI = [-.09, .42], p = .120 |
| *d*-AMP at Both | M = .13, SD = .07, CI = [-.01, .27], p = .032 | M = .01, SD = .09, CI = [-.16, .21], p = .451 | M = .14, SD = .09, CI = [-.04, .31], p = .064 |

**Study 8.1: The Effects of Dextromethamphetamine on Encoding**

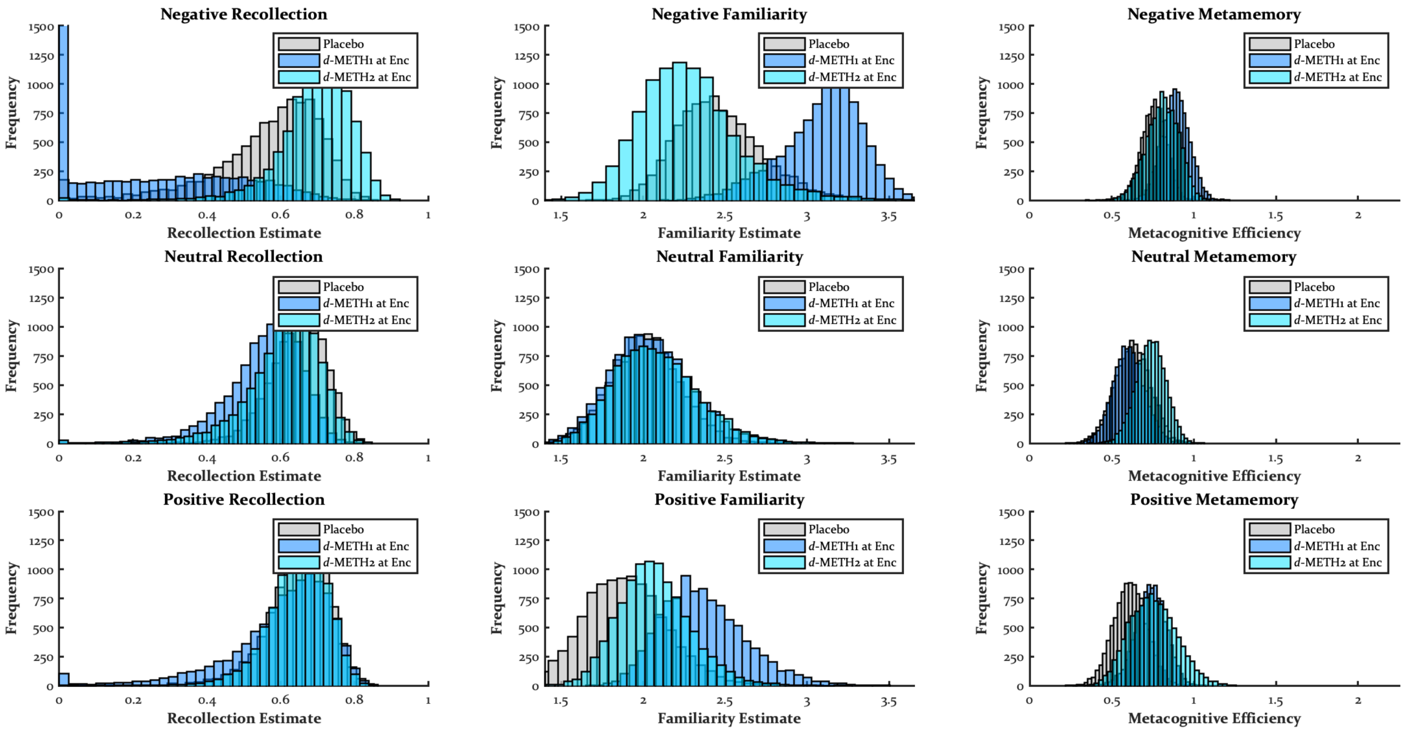


*Recollection Contrasts*

| | PLA vs. Enc1 | PLA vs. Enc2 | Enc1 vs. Enc2 |
|---|---|---|---|
| Negative | M = .40, SD = .23, CI = [-.06, .73], p = .050 | M = .13, SD = .16, CI = [-.10, .58], p = .149 | M = .54, SD = .23, CI = [.05, .82], p = .013 |
| Neutral | M = .09, SD = .11, CI = [-.10, .35], p = .186 | M = .04, SD = .11, CI = [-.14, .28], p = .349 | M = .05, SD = .11, CI = [-.19, .27], p = .284 |
| Positive | M = .04, SD = .14, CI = [-.17, .39], p = .390 | M = .00, SD = .11, CI = [-.20, .21], p = .494 | M = .05, SD = .15, CI = [-.18, .42], p = .408 |

| | Negative vs. Neutral | Negative vs. Positive | Neutral vs. Positive |
|---|---|---|---|

| | | | |
|---|---|---|---|
| Placebo | M = .08, SD = .15, CI = [-.14, .50], p = .305 | M = .08, SD = .15, CI = [-.14, .49], p = .285 | M = .00, SD = .08, CI = [-.14, .17], p = .532 |
| *d*-METH1 at Enc | M = .39, SD = .21, CI = [-.05, .69], p = .052 | M = .44, SD = .24, CI = [-.04, .77], p = .040 | M = .05, SD = .10, CI = [-.20, .23], p = .259 |
| *d*-METH2 at Enc | M = .09, SD = .12, CI = [-.14, .34], p = .166 | M = .05, SD = .10, CI = [-.16, .24], p = .258 | M = .04, SD = .13, CI = [-.19, .32], p = .368 |

*Familiarity Contrasts*

| | PLA vs. Enc1 | PLA vs. Enc2 | Enc1 vs. Enc2 |
|---|---|---|---|
| Negative | M = .63, SD = .33, CI = [-.07, 1.23], p = .037 | M = .19, SD = .35, CI = [-.52, .90], p = .284 | M = .82, SD = .34, CI = [.09, 1.41], p = .018 |
| Neutral | M = .04, SD = .24, CI = [-.41, .55], p = .457 | M = .06, SD = .25, CI = [-.39, .60], p = .419 | M = .02, SD = .25, CI = [-.44, .55], p = .480 |
| Positive | M = .48, SD = .23, CI = [.03, .95], p = .019 | M = .18, SD = .32, CI = [-.46, .80], p = .290 | M = .30, SD = .34, CI = [-.33, 1.00], p = .186 |

| | Negative vs. Neutral | Negative vs. Positive | Neutral vs. Positive |
|---|---|---|---|
| Placebo | M = .45, SD = .31, CI = [-.12, 1.11], p = .068 | M = .57, SD = .35, CI = [-.08, 1.28], p = .044 | M = .13, SD = .18, CI = [-.26, .43], p = .217 |
| *d*-METH1 at Enc | M = 1.04, SD = .24, CI = [.56, 1.51], p = .000 | M = .73, SD = .29, CI = [.13, 1.26], p = .009 | M = .32, SD = .19, CI = [-.09, .68], p = .058 |
| *d*-METH2 at Enc | M = .20, SD = .25, CI = [-.27, .74], p = .195 | M = .21, SD = .33, CI = [-.42, .92], p = .263 | M = .01, SD = .34, CI = [-.63, .71], p = .500 |

*Metamemory Contrasts*

| | PLA vs. Enc1 | PLA vs. Enc2 | Enc1 vs. Enc2 |
|---|---|---|---|
| Negative | M = .13, SD = .12, CI = [-.10, .38], p = .137 | M = .05, SD = .13, CI = [-.21, .29], p = .323 | M = .08, SD = .13, CI = [-.18, .33], p = .270 |
| Neutral | M = .02, SD = .09, CI = [-.14, .20], p = .418 | M = .11, SD = .12, CI = [-.12, .37], p = .191 | M = .13, SD = .11, CI = [-.07, .35], p = .110 |
| Positive | M = .13, SD = .10, CI = [-.07, .33], p = .107 | M = .14, SD = .13, CI = [-.11, .41], p = .144 | M = .01, SD = .13, CI = [-.25, .26], p = .451 |

| | Negative vs. Neutral | Negative vs. Positive | Neutral vs. Positive |
|---|---|---|---|

| | | | |
|---|---|---|---|
| Placebo | M = .13, SD = .13, CI = [-.12, .39], p = .159 | M = .15, SD = .12, CI = [-.09, .39], p = .120 | M = .01, SD = .10, CI = [-.19, .21], p = .428 |
| *d*-METH1 at Enc | M = .28, SD = .11, CI = [.08, .50], p = .004 | M = .15, SD = .12, CI = [-.07, .39], p = .096 | M = .13, SD = .09, CI = [-.05, .31], p = .076 |
| *d*-METH2 at Enc | M = .07, SD = .07, CI = [-.06, .22], p = .148 | M = .06, SD = .14, CI = [-.23, .31], p = .333 | M = .02, SD = .12, CI = [-.21, .26], p = .460 |

### Study 8.2: The Effects of Dextromethamphetamine on Consolidation

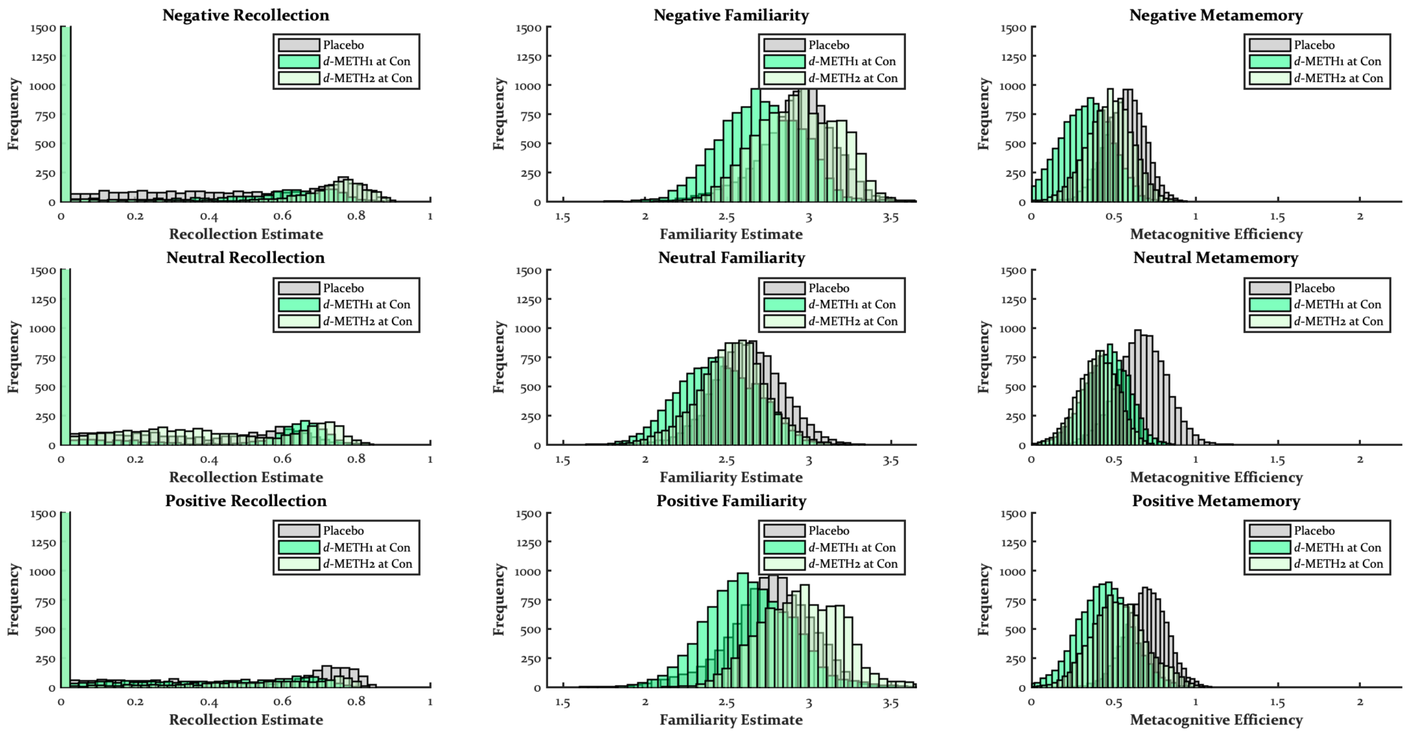


*Recollection Contrasts*

| | PLA vs. Con1 | PLA vs. Con2 | Con1 vs. Con2 |
|---|---|---|---|
| Negative | M = .07, SD = .16, CI = [-.01, .56], p = .283 | M = .05, SD = .15, CI = [-.10, .53], p = .321 | M = .02, SD = .10, CI = [-.05, .32], p = .413 |
| Neutral | M = .01, SD = .09, CI = [-.17, .27], p = .400 | M = .05, SD = .14, CI = [-.19, .43], p = .329 | M = .06, SD = .13, CI = [-.06, .45], p = .269 |
| Positive | M = .04, SD = .14, CI = [-.17, .49], p = .360 | M = .06, SD = .15, CI = [-.00, .54], p = .295 | M = .02, SD = .14, CI = [-.26, .42], p = .360 |

| | Negative vs. Neutral | Negative vs. Positive | Neutral vs. Positive |
|---|---|---|---|
| Placebo | M = .05, SD = .15, CI = [-.20, .48], p = .330 | M = .03, SD = .15, CI = [-.26, .50], p = .412 | M = .02, SD = .11, CI = [-.23, .30], p = .378 |
| *d*-METH1 at Con | M = .01, SD = .08, CI = [-.10, .24], p = .535 | M = .00, SD = .12, CI = [-.31, .30], p = .476 | M = .01, SD = .11, CI = [-.19, .35], p = .578 |
| *d*-METH2 at Con | M = .05, SD = .14, CI = [-.14, .46], p = .375 | M = .03, SD = .14, CI = [-.00, .55], p = .379 | M = .09, SD = .17, CI = [-.04, .56], p = .256 |

*Familiarity Contrasts*

| | PLA vs. Con1 | PLA vs. Con2 | Con1 vs. Con2 |
|---|---|---|---|
| Negative | M = .23, SD = .25, CI = [-.46, .49], p = .157 | M = .01, SD = .22, CI = [-.27, .59], p = .620 | M = .23, SD = .19, CI = [-.28, .52], p = .072 |
| Neutral | M = .19, SD = .18, CI = [-.23, .49], p = .131 | M = .11, SD = .21, CI = [-.31, .59], p = .277 | M = .09, SD = .21, CI = [-.43, .39], p = .258 |
| Positive | M = .12, SD = .27, CI = [-.67, .54], p = .196 | M = .20, SD = .26, CI = [-.09, .98], p = .143 | M = .32, SD = .24, CI = [-.04, .94], p = .031 |

| | Negative vs. Neutral | Negative vs. Positive | Neutral vs. Positive |
|---|---|---|---|
| Placebo | M = .29, SD = .19, CI = [-.18, .66], p = .069 | M = .17, SD = .21, CI = [-.35, .62], p = .144 | M = .12, SD = .18, CI = [-.31, .49], p = .180 |
| *d*-METH1 at Con | M = .26, SD = .15, CI = [.04, .68], p = .013 | M = .07, SD = .18, CI = [-.28, .50], p = .285 | M = .20, SD = .15, CI = [-.06, .62], p = .041 |
| *d*-METH2 at Con | M = .41, SD = .22, CI = [.10, .94], p = .008 | M = .02, SD = .21, CI = [-.27, .68], p = .543 | M = .43, SD = .26, CI = [.13, 1.16], p = .001 |

*Metamemory Contrasts*

| | PLA vs. Con1 | PLA vs. Con2 | Con1 vs. Con2 |
|---|---|---|---|
| Negative | M = .25, SD = .10, CI = [.06, .45], p = .006 | M = .09, SD = .12, CI = [-.13, .33], p = .211 | M = .15, SD = .14, CI = [-.13, .43], p = .138 |
| Neutral | M = .22, SD = .12, CI = [-.00, .46], p = .028 | M = .27, SD = .16, CI = [-.04, .58], p = .042 | M = .06, SD = .12, CI = [-.19, .30], p = .326 |
| Positive | M = .26, SD = .13, CI = [.01, .52], p = .016 | M = .17, SD = .11, CI = [-.05, .37], p = .061 | M = .09, SD = .15, CI = [-.18, .39], p = .272 |

| | Negative vs. Neutral | Negative vs. Positive | Neutral vs. Positive |
|---|---|---|---|
| Placebo | M = .10, SD = .15, CI = [-.19, .39], p = .240 | M = .13, SD = .10, CI = [-.07, .34], p = .100 | M = .03, SD = .12, CI = [-.20, .27], p = .410 |
| *d*-METH1 at Con | M = .13, SD = .12, CI = [-.10, .39], p = .139 | M = .12, SD = .13, CI = [-.14, .37], p = .166 | M = .01, SD = .13, CI = [-.23, .26], p = .475 |
| *d*-METH2 at Con | M = .08, SD = .12, CI = [-.16, .31], p = .254 | M = .06, SD = .11, CI = [-.15, .29], p = .304 | M = .13, SD = .14, CI = [-.14, .43], p = .174 |

**Study 9: The Effects of THC on Encoding**

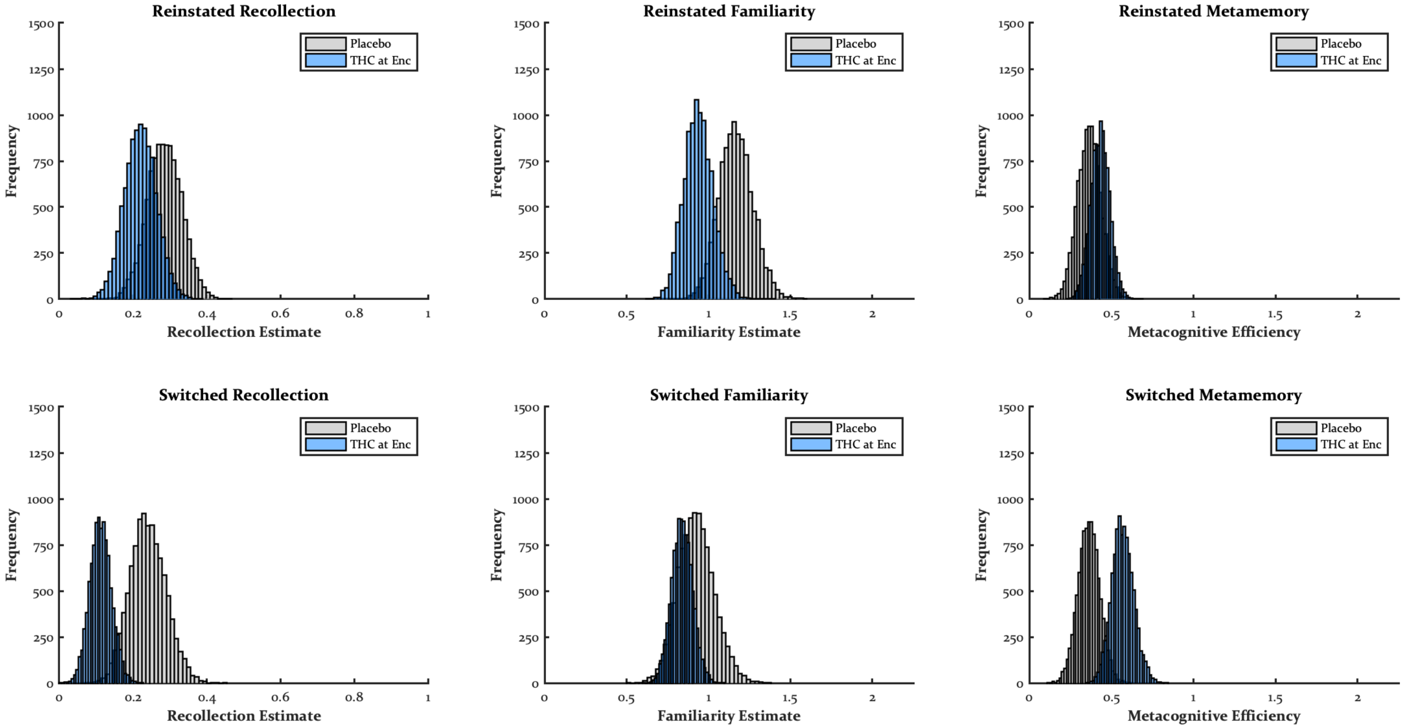


*Recollection Contrasts*

| | PLA vs. Enc |
|---|---|
| Reinstated | M = .07, SD = .04, CI = [-.02, .16], p = .065 |
| Switched | M = .13, SD = .03, CI = [.06, .19], p = .000 |

| | Reinstated vs. Switched |
|---|---|
| PLA | M = .07, SD = .04, CI = [-.02, .16], p = .065 |
| THC at Enc | M = .13, SD = .03, CI = [.06, .19], p = .000 |

*Familiarity Contrasts*

| | PLA vs. Enc |
|---|---|
| Reinstated | M = .23, SD = .10, CI = [.04, .42], p = .010 |
| Switched | M = .09, SD = .12, CI = [-.15, .33], p = .240 |

| | Reinstated vs. Switched |
|---|---|
| PLA | M = .24, SD = .09, CI = [.06, .40], p = .005 |
| THC at Enc | M = .10, SD = .07, CI = [-.02, .24], p = .052 |

*Metamemory Contrasts*

| | PLA vs. Enc |
|---|---|
| Reinstated | M = .07, SD = .06, CI = [-.05, .19], p = .132 |
| Switched | M = .21, SD = .08, CI = [.06, .37], p = .002 |

| | Reinstated vs. Switched |
|---|---|
| PLA | M = .00, SD = .05, CI = [-.10, .11], p = .485 |
| THC at Enc | M = .14, SD = .06, CI = [.01, .27], p = .014 |

**Study 10: The Effects of THC on Retrieval**

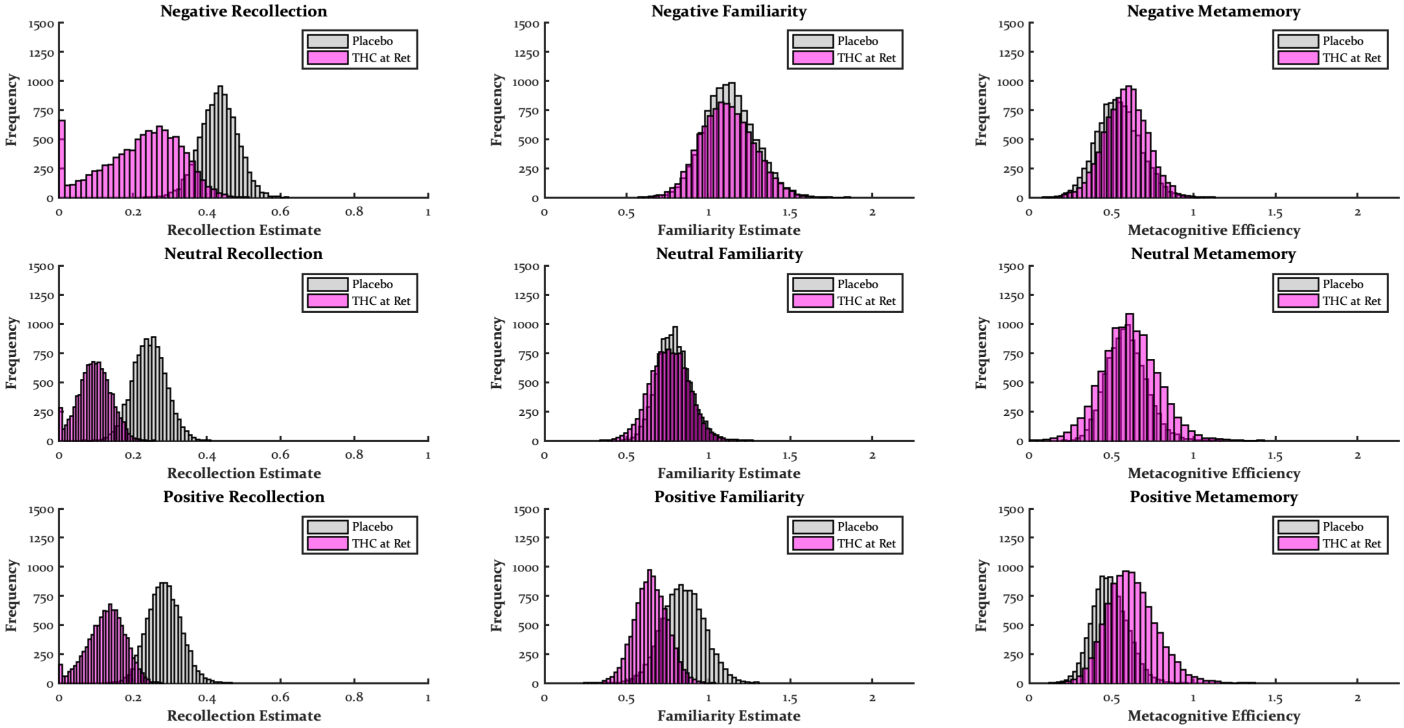


*Recollection Contrasts*

| | PLA vs. Ret |
|---|---|
| Negative | M = .21, SD = .13, CI = [-.00, .49], p = .026 |
| Neutral | M = .15, SD = .07, CI = [.02, .28], p = .010 |
| Positive | M = .16, SD = .07, CI = [.03, .30], p = .004 |

| | Negative vs. Neutral | Negative vs. Positive | Neutral vs. Positive |
|---|---|---|---|
| Placebo | M = .19, SD = .04, CI = [.11, .26], p = .000 | M = .15, SD = .05, CI = [.05, .24], p = .002 | M = .04, SD = .04, CI = [-.03, .11], p = .133 |

| | | | |
|---|---|---|---|
| THC at Ret | M = .13, SD = .09, CI = [-.06, .26], p = .107 | M = .09, SD = .08, CI = [-.09, .22], p = .152 | M = .03, SD = .03, CI = [-.03, .09], p = .152 |

*Familiarity Contrasts*

| | PLA vs. Ret |
|---|---|
| Negative | M = .01, SD = .19, CI = [-.36, .38], p = .462 |
| Neutral | M = .03, SD = .12, CI = [-.21, .27], p = .416 |
| Positive | M = .20, SD = .13, CI = [-.06, .45], p = .067 |

| | Negative vs. Neutral | Negative vs. Positive | Neutral vs. Positive |
|---|---|---|---|
| Placebo | M = .34, SD = .10, CI = [.15, .55], p = .001 | M = .28, SD = .15, CI = [-.01, .57], p = .030 | M = .06, SD = .10, CI = [-.14, .27], p = .285 |
| THC at Ret | M = .35, SD = .14, CI = [.10, .65], p = .001 | M = .47, SD = .15, CI = [.19, .77], p = .000 | M = .11, SD = .08, CI = [-.05, .28], p = .092 |

*Metamemory Contrasts*

| | PLA vs. Ret |
|---|---|
| Negative | M = .05, SD = .17, CI = [-.31, .37], p = .383 |
| Neutral | M = .02, SD = .19, CI = [-.35, .42], p = .473 |
| Positive | M = .13, SD = .16, CI = [-.18, .46], p = .203 |

| | Negative vs. Neutral | Negative vs. Positive | Neutral vs. Positive |
|---|---|---|---|
| Placebo | M = .05, SD = .17, CI = [-.28, .39], p = .390 | M = .05, SD = .16, CI = [-.25, .37], p = .384 | M = .10, SD = .12, CI = [-.14, .34], p = .212 |
| THC at Ret | M = .02, SD = .22, CI = [-.41, .47], p = .461 | M = .04, SD = .17, CI = [-.28, .38], p = .427 | M = .01, SD = .20, CI = [-.38, .41], p = .476 |
